# Activity-driven chromatin organization during interphase: compaction, segregation, and entanglement suppression

**DOI:** 10.1101/2024.01.22.576729

**Authors:** Brian Chan, Michael Rubinstein

**Author notes:** **Author Contributions:** BC and MR designed the research, analyzed data, and wrote the paper. BC performed the research. **Competing Interest Statement:** The authors declare no competing interests.

## Abstract

In mammalian cells, the cohesin protein complex is believed to translocate along chromatin during interphase to form dynamic loops through a process called active loop extrusion. Chromosome conformation capture and imaging experiments have suggested that chromatin adopts a compact structure with limited interpenetration between chromosomes and between chromosomal sections. We developed a theory demonstrating that active loop extrusion causes the apparent fractal dimension of chromatin to cross over between two and four at contour lengths on the order of 30 kilo-base pairs (kbp). The anomalously high fractal dimension *D* = 4 is due to the inability of extruded loops to fully relax during active extrusion. Compaction on longer contour length scales extends within topologically associated domains (TADs), facilitating gene regulation by distal elements. Extrusion-induced compaction segregates TADs such that overlaps between TADs are reduced to less than 35% and increases the entanglement strand of chromatin by up to a factor of 50 to several Mega-base pairs. Furthermore, active loop extrusion couples cohesin motion to chromatin conformations formed by previously extruding cohesins and causes the mean square displacement of chromatin loci during lag times (Δ*t*) longer than tens of minutes to be proportional to Δ*t*^1/3^. We validate our results with hybrid molecular dynamics – Monte Carlo simulations and show that our theory is consistent with experimental data. This work provides a theoretical basis for the compact organization of interphase chromatin, explaining the physical reason for TAD segregation and suppression of chromatin entanglements which contribute to efficient gene regulation.

**SIGNIFICANCE STATEMENT:** During interphase, cells must compact chromatin such that gene promoters and their regulatory elements frequently contact each other in space. However, cells also need to insulate promoters from regulatory elements in other genomic sections. Using polymer physics theory and computer simulations, we propose that the cohesin protein complex actively extrudes chromatin into topologically associated domains (TADs) with an anomalously high fractal dimension of *D* ≈ 4 while suppressing spatial overlap between different TADs. Our model suggests that the fast kinetics of active loop extrusion compared to the slow relaxation of chromatin loops maintains a dense chromatin organization. This work presents a physical framework explaining how cohesin contributes to effective transcriptional regulation.

## INTRODUCTION

Gene transcription during interphase can be regulated by spatial colocalization of promoters with enhancers or silencers located up to hundreds of kilo-base pairs (kbp) away (1) (see Fig. 1A). These distal regulatory elements frequently act on genes within the same topologically associated domain (TAD) (2–4). TADs are continuous sections of chromatin that preferentially associate in space, identified by squares along the main diagonal of Hi-C contact maps (5, 6). One model for TAD formation is loop extrusion, in which the cohesin protein complex threads chromatin into growing loops until stopped by the CCCTC-binding factor (CTCF) at TAD boundaries (7–10) (see Fig. 1B).

**Figure 1:**
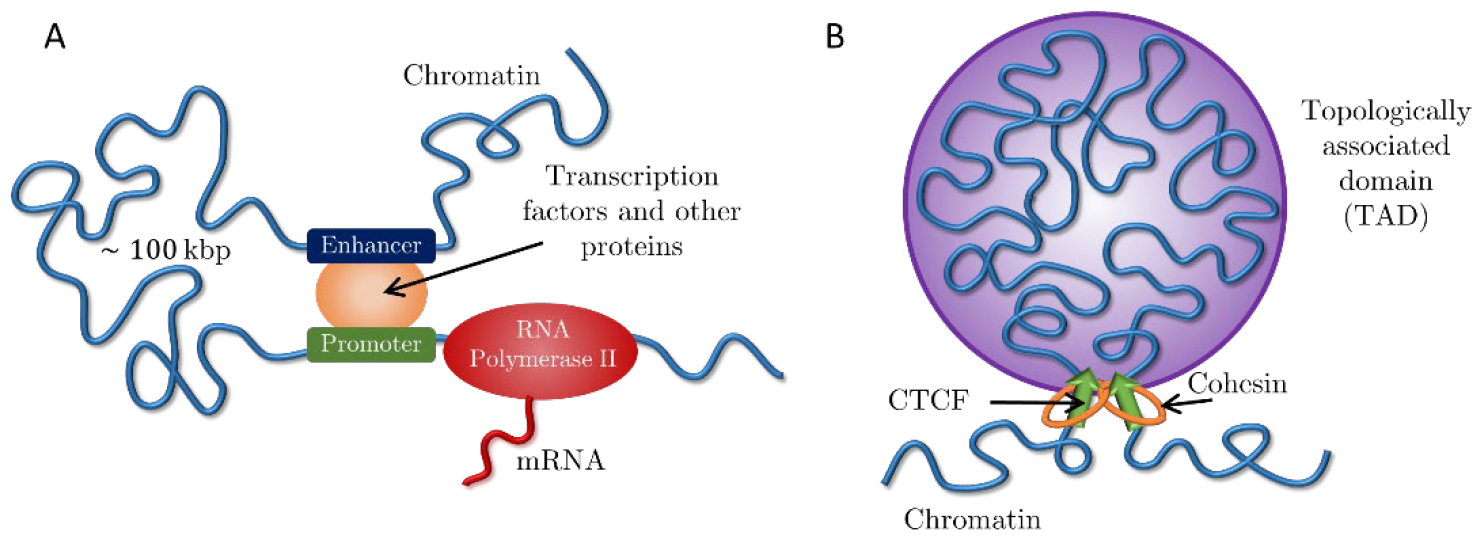
Active processes on chromatin. A) Eukaryotic transcription by RNA Polymerase II regulated by an enhancer-promoter interaction and transcription factors. B) Active loop extrusion by cohesin forms TADs anchored by CTCF proteins.

We consider *active* loop extrusion, in which cohesin uses ATP to move in biased directions toward domain boundaries (7–9, 11, 12). Active processes that consume energy are widespread throughout the cell and are known to modify the conformation and dynamics of different biopolymers, including chromatin. In general, ATP depletion decreases the diffusion constant of chromosomal loci (13) and eliminates their coherent dynamics, in which loci displacements correlate with those of other loci in close spatial proximity (14–16). A well-known example of an active process on chromatin is transcription, in which RNA Polymerase II (RNAPII) produces mRNA molecules from a DNA template (17–19) (see Fig. 1A). In contrast to loop extrusion, RNAPII translocates at slower rates (0.01 – 0.1 kbp per second) compared to cohesin (0.1 – 1 kbp per second) and does not hold two chromatin loci together (20–24).

To effectively modulate transcription, promoters and distal regulatory elements such as enhancers and silencers should frequently come into physical contact. In other words, the contact probability *P*(*s*) must decay slowly with the genomic distance *s* between them. Hi-C and Micro-C experiments have shown at least three power law scaling regimes for contact probabilities (see Fig. 2), each characterized by 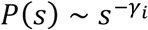, where ∼ indicates proportionality and the subscript *i* indicates different regimes. The three regimes typically have 1 ≤ *γ*_1_ ≤ 1.5, *γ*_2_ ≈ 0.75, and 1 ≤ *γ*_3_ ≤ 1.5 (5, 8, 25–32). The crossovers between these three regimes vary between experiments and are not well defined in part due to the ranges in observed scaling exponents. However, the crossover between *γ*_1_ and *γ*_2_ is typically on the order of 30—50 kbp, and the crossover between *γ*_2_ and *γ*_3_ is typically on the order of 200—500 kbp. Within a mean-field approximation *γ*_*i*_ ≈ 3/*D*_*i*_, where *D*_*i*_ is the fractal dimension (33). Fractal dimension describes how the mass *m* of an object grows with its physical size *r* such that *m* ∼ *r*^*D*^ (34). Mapping *γ* to fractal dimension yields 2 ≤ *D*_1_ ≤ 3 for the first regime, *D*_2_ ≈ 4 for the second, and 2 ≤ *D*_3_ ≤ 3 for the third. The second scaling regime occurs on genomic length scales on the order of a typical TAD (35, 36). Depleting chromatin-bound cohesins increases *γ*_2_, suggesting that loop extrusion compacts chromatin on the scale of TADs by increasing its fractal dimension (5, 25, 37). Furthermore, experimental evidence suggests that cohesins assist in efficient target search by transcription factors (TFs) and regulate transcriptional bursting probabilities (32, 38, 39), which hints at a connection between loop extrusion, a window of decreased *γ*, and transcriptional regulation.

**Figure 2:**
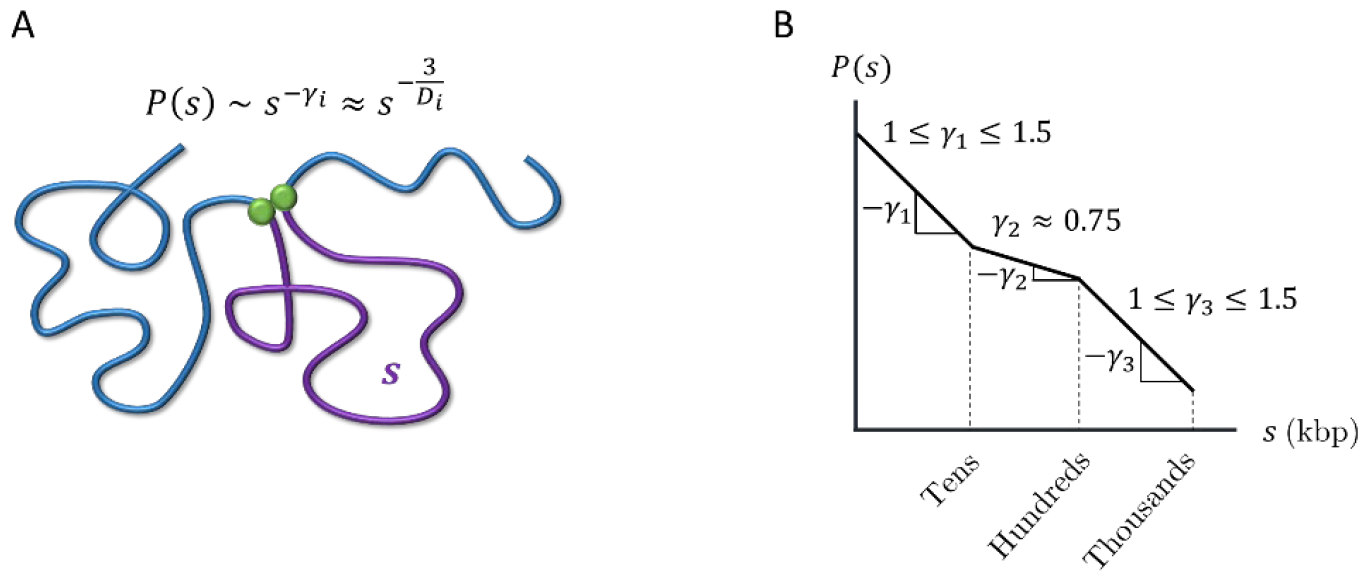
Contact probabilities in interphase chromatin. A) Schematic of a contact between two loci (green circles) separated by genomic distance *s* (purple curve). The contact probability between the two green loci *P*(*s*) is proportional to a power law 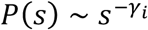 where the subscript *i* indicates a specific regime. *γ*_*i*_ ≈ 3/*D*_*i*_ within a mean-field approximation, where *D*_*i*_ is the fractal dimension. B) Schematic plot of the average contact probability *P*(*s*) for chromatin as a function of genomic separation *s* with three scaling regimes on a log-log scale.

Cells must ensure that regulatory elements do not act on off-target genes, such as those in other TADs. Super-resolution imaging suggests that neighboring TADs are spatially segregated (39– 41). We argue that the increase in fractal dimension due to active loop extrusion would suppress overlaps between chromatin sections, where we define the overlap parameter as the number of chromatin sections of similar genomic lengths sharing the same volume (34). The entanglement length is a characteristic scale along the polymer above which topological constraints that prevent strand crossing dominate dynamic properties (34). An estimate of this entanglement is the strand length for which the overlap parameter is ∼10. If chromatin were to adopt a random walk conformation with *D* ≈ 2 between entanglements, the entanglement genomic length in the absence of activity and looping *N*_*e,passive*_ would be 50—100 kbp (33, 42–44). However, this is in contrast to evidence that chromosomes and chromatin compartments do not intermingle extensively (45). We argue that extruded chromatin loops are *not equilibrated* and that the kinetics of active loop extrusion cause an increase of fractal dimension in compact loops and suppress entanglements by two orders of magnitude.

Several *equilibrium* polymer models are consistent with fractal dimensions larger than *D* ≈ 2 (see Fig. 3): i) a crumpled polymer with an exponential distribution of equilibrated loops (CPEL– Fig. 3A) (44, 46), ii) the fractal loopy globule (FLG) model for a melt of non-concatenated ring polymers (Fig. 3B) (47), and iii) a ring polymer in an array of fixed topological obstacles, which can be mapped onto double folded lattice animals (DFLA – Fig. 3C) (48, 49). In CPEL, looped and un-looped chromatin sections have *D* ≈ 2 (Fig. 3A). Loops shorten the effective contour length between genomic loci, causing a higher apparent fractal dimension on scales larger than the average loop length. Additional mass within loops contributes to entanglement dilution as in bottlebrush polymer systems (50). The FLG describes a crossover between fractal dimensions of *D* ≈ 2 and *D* ≈ 3 at the entanglement length smaller than a single ring such that loopy polymer sections maintain a constant degree of overlap with each other on all larger length scales (Fig. 3B) with large overlap parameter ∼10. In DFLA, polymers have *D* ≈ 2 on the smallest length scales as well (Fig. 3C). Double folds of ring polymers caused by their topological interactions with fixed networks produce lattice animal structures, resulting in *D* ≈ 4 above the scale of the typical fold or loop. This regime extends until density saturation, above which *D* ≈ 3.

**Figure 3:**
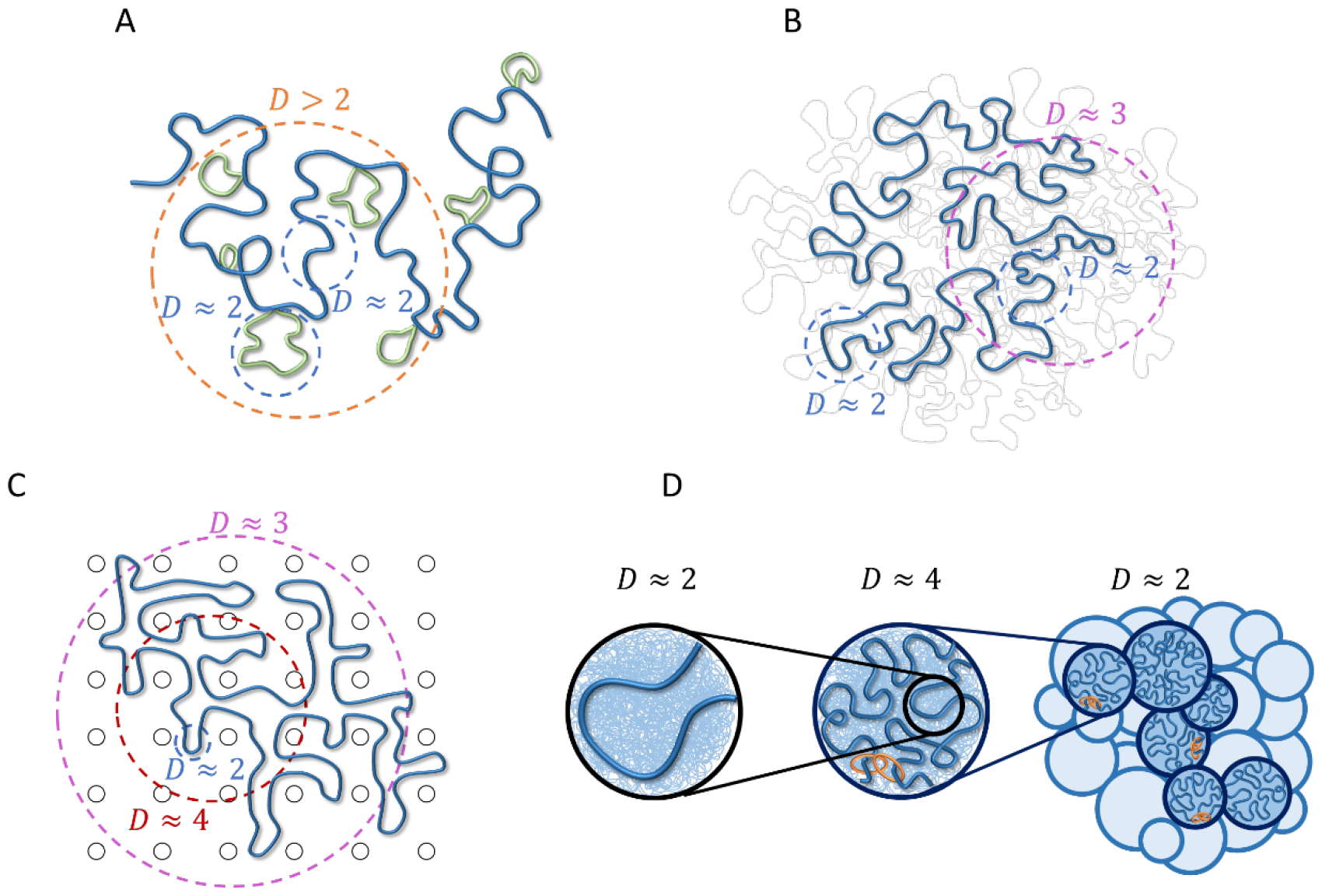
Polymer models relevant to chromatin organization consistent with fractal dimension *D* > 2. Dashed circles indicate typical length scales with a given fractal dimension. A) CPEL (44, 46). Looped and un-looped sections of the same polymer chain are drawn in green and blue, respectively. B) FLG (47). Fractal behavior is shown for one polymer ring of interest (thick blue curve). Thin gray curves represent other polymers in a melt. C) DFLA (48, 49). Fractal behavior is shown for one polymer ring of interest (thick blue curve). Black circles represent obstacles created by fixed networks. D) Active loop extrusion (this paper). Short, relaxed sections of chromatin with fractal dimension *D* ≈ 2 (left) pack together to form extruded loops with *D* ≈ 4 (center, where the most recently extruded section of a loop is shown in blue, the rest of the loop represented by the thin blue lines in the background, and orange rings represent cohesin). Extruded loops form a random walk (dark blue circles in the right-hand schematic) with *D* ≈ 2 surrounded by other chromatin sections (light blue circles). Models (A) – (C) are equilibrium, while (D) is non-equilibrium.

CPEL, FLG, and DFLA are all *equilibrium* models, while our model explicitly considers *non-equilibrium* activity. Our model of active loop extrusion suggests that a crossover from *D* ≈ 2 to *D* ≈ 4 occurs on genomic lengths smaller than a typical loop extruded by a loop extruding protein. We argue that this compaction is a non-equilibrium process that occurs due to the fast kinetics of loop extrusion in comparison with loop relaxation. A random walk conformation of looped sections with *D* ≈ 2 is recovered above the length scales of an average loop (see Fig. 3D). Compaction by loop extrusion suppresses entanglements. The range of the applicability of our model depends on the interplay between the extrusion velocity and chromatin relaxation rate, which have yet to be resolved experimentally. As described below, we suggest that active loop extrusion significantly modifies chromatin conformation given the observed cohesin extrusion speeds and chromatin dynamics.

The main parameters used to describe active loop extrusion are processivity and separation. Processivity *λ* is the average genomic length extruded by an unobstructed cohesin. Separation *d* is the average genomic length between chromatin-bound cohesins or the inverse of the linear density of cohesins. Both *λ* and *d* are predicted to be on the order of 200 kbp, suggesting limited loop nesting (7, 11, 51). Extrusion velocity *v*_*ex*_ is the average genomic length extruded per unit time and is equal to the processivity divided by the average residence time of an unobstructed cohesin *τ*_*res*_ · *v*_*ex*_ is on the order of 0.1 – 1 kbp per second and *τ*_*res*_ is on the order of 20 – 30 minutes (22–24, 52, 53).

In this work, we develop a scaling-level theory to describe chromatin organization due to active loop extrusion. We first describe the effect of active extrusion on relaxed chromatin with fractal dimension *D* ≈ 2. We test the predictions of our model by hybrid molecular dynamics—Monte Carlo (MD—MC) simulations of a single extrusion cycle on relaxed polymer chains in a theta-like solvent. Next, we extend our model to chromatin conformation regulated by steady-state extrusion (in which cohesins randomly bind and unbind along the genome) with and without TAD anchors. We compare our model to simulations of steady-state extrusion on 1 Mbp chromatin sections in a theta-like solvent. We then discuss how active loop extrusion kinetically suppresses overlaps of neighboring sections of chromatin, dilutes entanglements, and segregates TADs. Our model also describes how extrusion deforms chromatin outside of growing loops and the spatial dynamics of actively extruding cohesins and chromatin loci. This model explains the anomalous scaling exponent observed in contact probabilities *P*(*s*) ∼ *s*^−0.75^ and supports active loop extrusion as a mechanism for segregating TADs, thereby enhancing contacts between promoters and regulatory elements within the same TAD while suppressing off-target interactions.

## RESULTS

### Model description

Consider chromatin discretized into loci each with spatial size *b* and *z* base pairs (see Fig. 4). If unperturbed (without loop extrusion), chromatin has fractal dimension *D* = 2 for genomic lengths longer than *z*: a section with genomic length *s* has mean square size

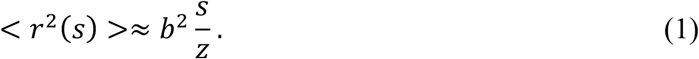

**Figure 4:**
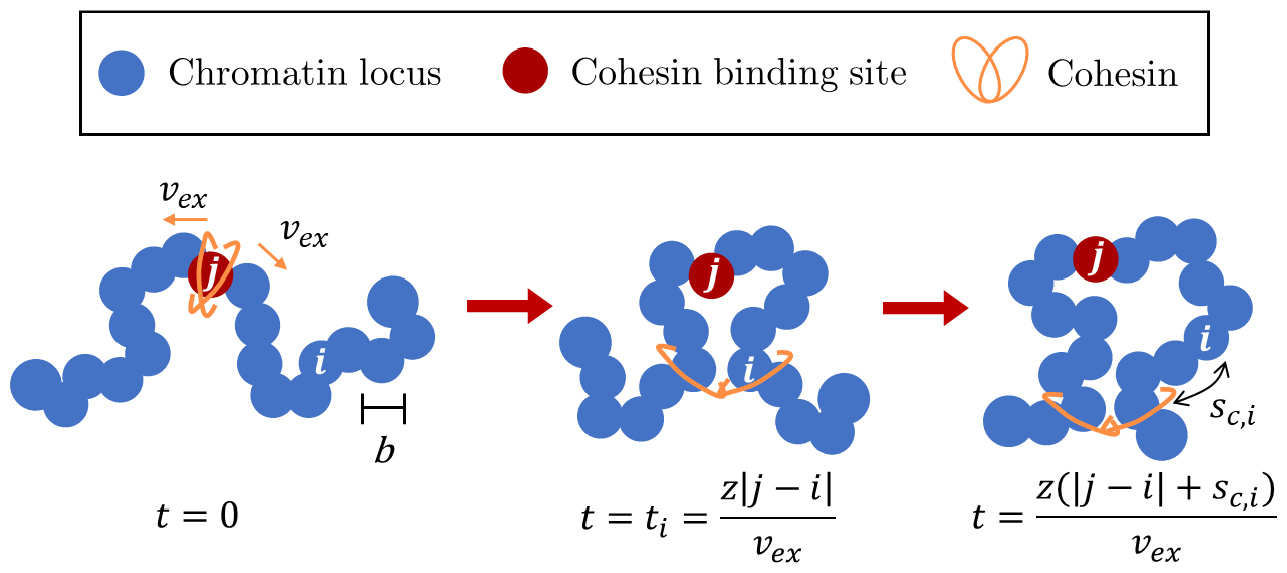
Schematic of active loop extrusion. Chromatin is discretized into loci with size *b* each representing *z* base pairs. Cohesin domains bind to locus *j* and extrude away from one another with average curvilinear velocity *v*_*ex*_ in units of the number of base pairs per unit time. Locus *i* is extruded at the time *t*_*i*_ = *z*|*j* − *i*|/*v*_*ex*_. At time *z*(|*j* − *i*| − *s*_*c,i*_)/*v*_*ex*_ > *t*_*i*_, locus *i* is *s*_*c,i*_ = *v*_*ex*_ (*t* − *t*_*i*_) base pairs away from the cohesin.

Throughout this work, ≈ indicates approximate equivalence within a factor on the order of unity and ∼ indicates proportionality. For definiteness we let each locus represent *z* = 2 kbp with *b* ≈ 50 nm (see Supporting Information (SI) for estimate), assuming the persistence length of chromatin is of equal or smaller size. While the persistence length of chromatin is not known, the principles of our model still hold for different choices of persistence length and locus size. In the absence of active forces, loci follow Rouse-like dynamics on length scales smaller than entanglement length with mean square displacement (MSD)

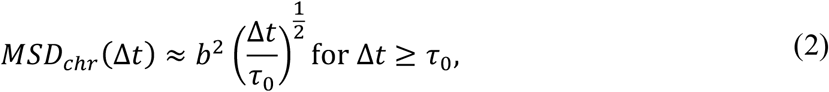

where Δ*t* is the lag time and *τ*_0_ is the locus diffusion time (34). The locus diffusion time is the time it takes a locus to move by thermal motion a distance on the order of its size. The relaxation time of a section with genomic length *s* and fractal dimension *D* = 2 is approximately

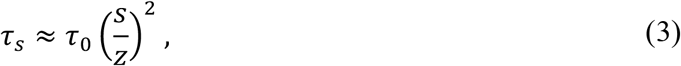

which is the time it takes for the section with genomic length *s* to move by thermal motion a distance on the order of its unperturbed size (34).

We model cohesin as having two domains that bind to chromatin (see Fig. 4). Domains extrude independently and in opposite directions along the chromatin contour with average curvilinear velocity *v*_*ex*_ in units of genomic length per unit time. The extrusion velocity *v*_*ex*_ and locus diffusion time *τ*_0_ define a dimensionless parameter that we call the “extrusion ratio”

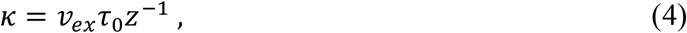

which is the ratio of rates of active and passive dynamics for a locus of *z* base pairs, like a Péclet number describing the ratio between flow and diffusion effects on some length scale. The extrusion ratio is the number of loci extruded per diffusion time of a locus. The genomic length unimpededly extruded during time *t* is *v*_*ex*_*t*. Since the two cohesin domains extrude independently, the average genomic length of the loop (loop length) at time *t* is

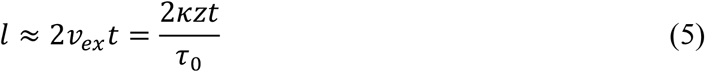

and the cohesin processivity is *λ* ≈ 2*v*_*ex*_ *τ*_*res*_ = 2*κzτ*_*res*_/*τ*_0_. Recall that cohesin processivity is defined as the average loop length extruded by unobstructed cohesin during the average residence time *τ*_*res*_ it is bound to chromatin (see Introduction).

Several experiments have observed chromatin loci with *MSD*_*chr*_(Δ*t*) ≈ *D*_*app*_Δ*t*^1/2^ consistent with Rouse-like dynamics where *D*_*app*_ ranges between ≈10^−3^ μm^2^ s^-1/2^ and ≈10^−2^ μm^2^ s^-1/2^ (54–59). For discretization of *z* = 2 kbp per locus each with size *b* ≈ 50 nm and *v*_*ex*_ ≈ 0.1 kbp per second, we estimate the extrusion ratio for cohesin (Eq. (4)) to be on the order of *κ* ≈ 0.003 − 0.3 (see SI). Note that the wide range of observed *D*_*app*_ causes a factor of 100 between the lower and upper estimates of the extrusion ratio. For definiteness, we use *κ* ≈ 0.2 to make biological estimates throughout the paper and provide ranges of estimated parameters in Table S2.

### Extrusion forms compact chromatin loops composed of overlapping relaxed sections

We start by describing the conformation of a single loop produced by active loop extrusion on a relaxed chromatin section with *D* ≈ 2. The main concept of our model is that chromatin sections longer than genomic length *g*_*min*_ ≈ *z*/*κ* relax slower than the time it takes a single cohesin domain to extrude them. The relaxation time of a chromatin section with genomic length *s* is proportional to *s*^2^, the square of its genomic length, while the time it takes to extrude it is proportional to *s* (see Eqs. (3) and (5) and Fig. 5A). The asymptotic behavior of the mean square size of a loop with genomic length *l* is

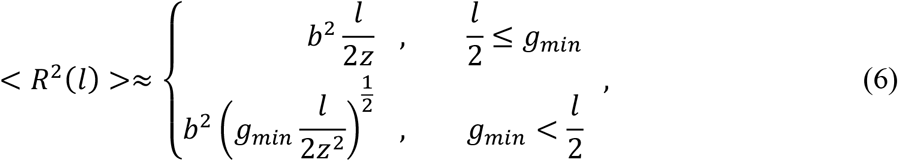

as plotted in Figure 5B. This scaling behavior could be impacted by the spatial mobility of cohesins (see SI). For a short loop with *l* ≤ 2*g*_*min*_, the entire loop is relaxed (see Eq (1)). Longer loops cannot relax, so their sizes are determined by subdiffusive Rouse-like dynamics with < *R*^2^(*l*) >∼ *t*^1/2^ ∼ *l*^1/2^. These loops are composed of multiple relaxed sections (see Fig. 5C), the smallest with genomic length *g*_*min*_ and the largest with genomic length

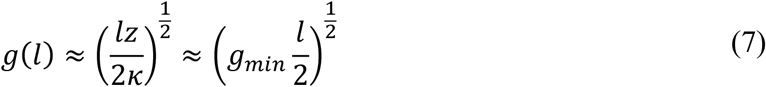

and mean square size *ξ*^2^(*l*) ≈ *b*^2^[*g*_*min*_*l*/(2*z*^2^)]^1/2^ (see SI for details). For an extrusion ratio of *κ* ≈ 0.2 and processivity *λ* = *l* ≈ 200 kbp, the smallest relaxed chromatin section has genomic length *g*_*min*_ ≈ 10 kbp and the largest has genomic length *g*(*λ*) ≈ 30 kbp.

**Figure 5:**
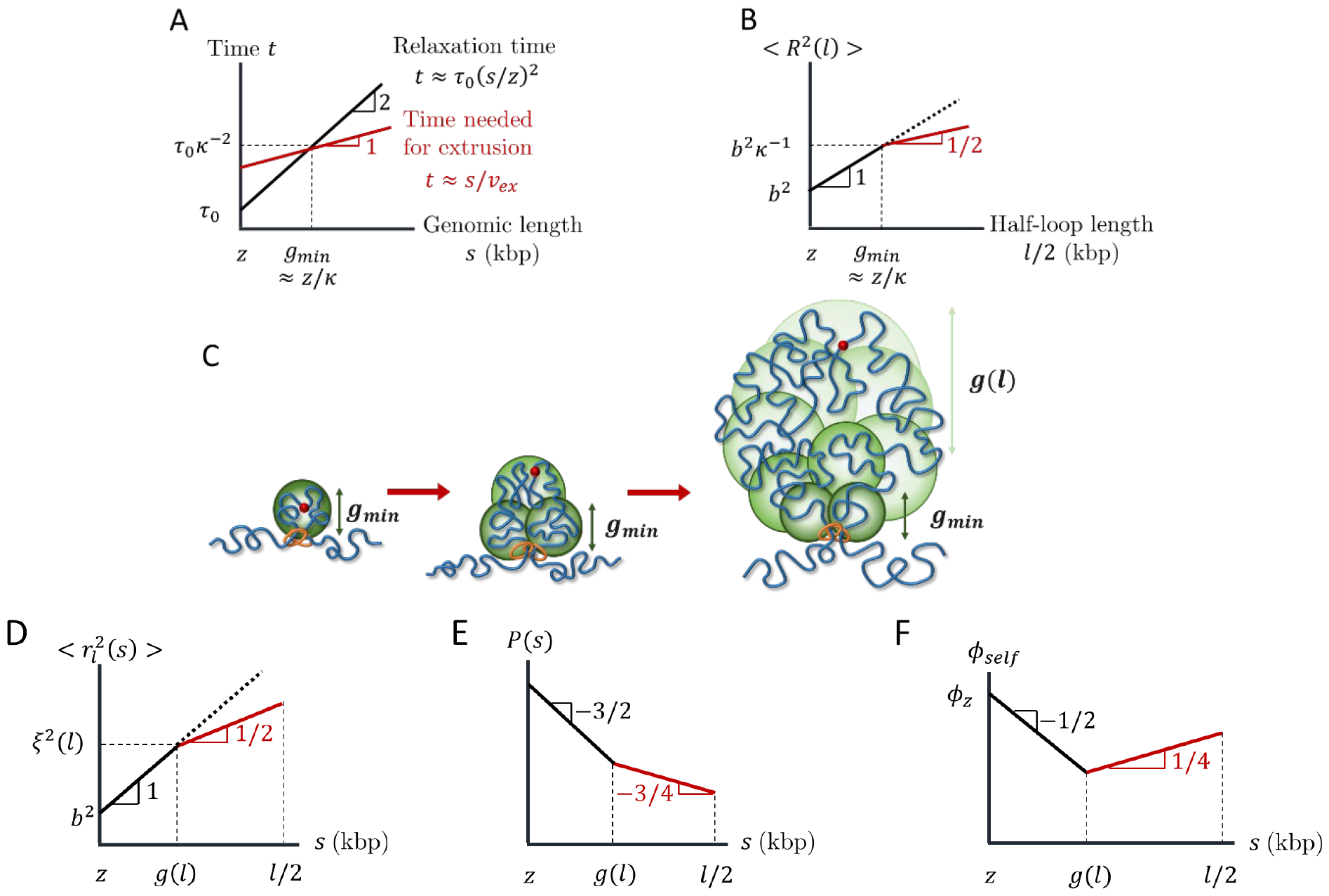
Structure of an actively extruded loop. All schematic plots are on log-log scales. A) Schematic plot comparing the relaxation time (black) and time needed for a single cohesin domain to extrude (red) a chromatin section with genomic length *s*. B) Schematic plot of the mean square size of a loop with genomic length *l*. The red line segment indicates the regime consistent with *D* = 4. The dotted line shows the mean square size of a chromatin section with genomic length *l*/2 without extrusion. C) Relaxed sections (green circles) in a growing loop overlap in space forming a fractal with dimensionality *D* = 4. The smallest relaxed section has the genomic length *g*_*min*_ ≈ *z*/*κ*. The genomic length of each relaxed section (green circles) grows from *g*_*min*_ next to the cohesin to *g*(*l*) ≈ (*g*_*min*_ *l*/2)^1/2^ ≈ [*lz*/(2*κ*)]^1/2^ next to the cohesin binding site (red circle). D) Mean square distance between two loci separated by a genomic distance *s* within a loop of genomic length *l* (see Eq. (8)). The average is taken over all loci pairs within the loop extruded by the same cohesin domain. The red line segment indicates the regime consistent with *D* = 4. The dotted line shows the mean square distance without extrusion. E) Average contact probability between loci separated by genomic length *s* within an extruded loop of length *l*. The red line segment indicates the regime consistent with *D* = 4. F) Volume fraction *ϕ*_*self*_ of a chromatin section of length *s* within its own pervaded volume where the section is part of a loop with length *l*. The red line segment indicates the regime consistent with *D* = 4.

Consider 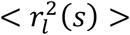, the mean square distance between any two loci separated by a genomic distance *s* extruded by the same cohesin domain in a loop with length *l*. The average is taken over all loci pairs separated by *s* within the loop. Like Equation (6), 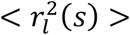 crosses over between power laws ∼ *s* and ∼ *s*^1/2^ because short genomic sections are relaxed while long sections are not:

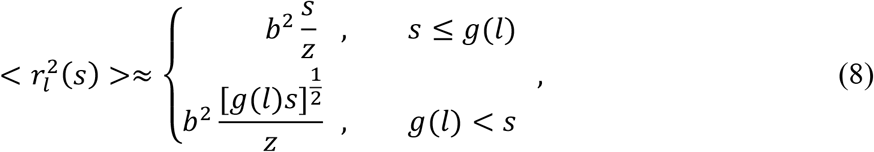

see Figure 5D. The crossover between the two power laws occurs at genomic lengths on the order of *g*(*l*) (Eq. (7)) because the largest relaxed sections with the largest number of loci pairs dominate the mean square distances averaged over all loop loci. Equation (8) is consistent with a fractal dimension of *D* ≈ 4 on genomic and spatial length scales longer than *g*(*l*) and *ξ*(*l*) respectively. Recall that within a mean-field approach, the contact probability function 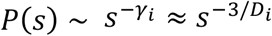 reflects the fractal behavior of the chromatin section of interest (see Introduction). Since compact loops transition between *D* ≈ 2 and *D* ≈ 4, the contact probability *P*(*s*) within a single extruded loop is proportional to

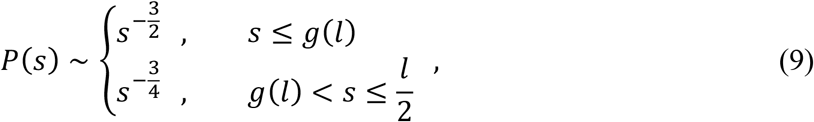

where *l*/2 is half of the loop length, after which contact probability increases (see Fig. 5E).

How can chromatin loops have a fractal dimension of four in three-dimensional space? Analogous to randomly branched ideal polymers, the fractal dimension below spatial dimension 3 on small length scales creates enough space within a loop’s pervaded volume for more compact structures with fractal dimension above 3 for a finite interval of larger length scales (34). Fig. 5F sketches the volume fraction *ϕ*_*self*_ for a section with genomic length *s* within its pervaded volume, where the section is part of a loop with genomic length *l*. We define volume fraction as the physical volume of chromatin (nucleosomes and linker DNA) of a genomic section with genomic length *s* divided by the approximately spherical volume spanned by the section (see SI). Smaller loop sections with *s* < *g*(*l*) are relaxed and therefore ideal with *D* ≈ 2. The volume fraction *ϕ*_*self*_ within these small sections initially decreases with section genomic length *s* because the genomic length of a section grows slower than its pervaded volume ∼ *s*^3/2^. When *D* ≈ 4, *ϕ*_*self*_ increases because the genomic length *s* and physical volume of the section grows faster than its pervaded volume ∼ *s*^3/4^. The volume fraction within a loop with genomic length *l* reaches *ϕ*_*self*_ ≈ 2*ϕ*_*z*_[*lz*^2^*g*(*l*)^−3^/2]^1/4^. *ϕ*_*z*_ is the volume fraction of chromatin within one locus, which we estimate to be ≈ 0.1 for *z* = 2 kbp (see SI). For *κ* = 0.2 and *z* = 2 kbp, *ϕ*_*self*_ within a loop with a typical genomic length *l* ≈ *λ* ≈ 200 kbp approaches ≈ 0.06, which is much smaller than unity. As such, a fractal dimension of four is possible for a wide range of genomic lengths.

To test our theory, we use hybrid MD—MC simulations to model active loop extrusion on a single bead-spring polymer chain in a theta-like solvent with each bead representing 1 kbp and Kuhn length ≈ 2*σ*_*LJ*_, where *σ*_*LJ*_ is the bead diameter. See Materials and Methods and SI for details. Relaxed polymer conformations in theta-like solvents are random walks with *D* ≈ 2 on scales longer than the Kuhn length, like dense solutions or melts of flexible linear polymers between the correlation and entanglement length scales, similar to conditions in the nucleus (34). The results of simulations of a single cohesin extruding a loop on an initially relaxed chain are consistent with Equations (8) and (9) (see Fig. 6).

**Figure 6:**
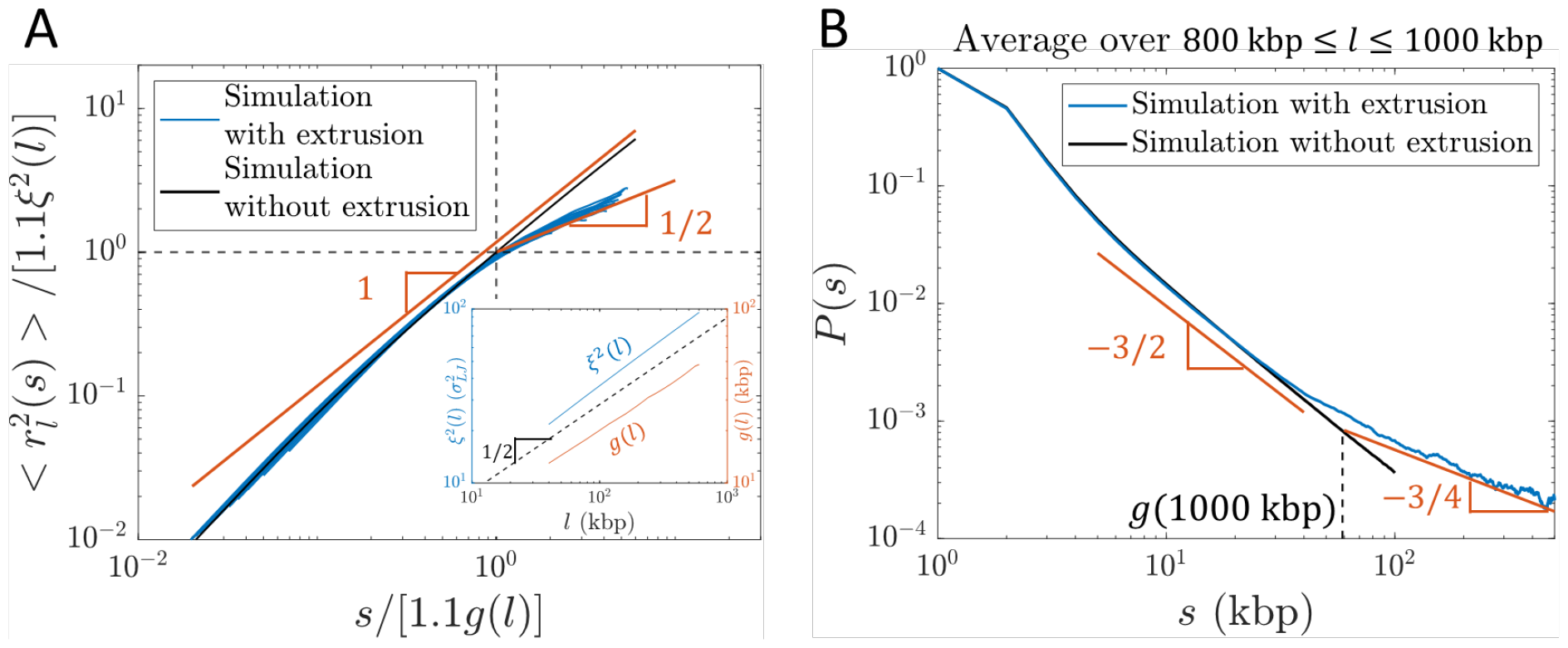
Internal structure of actively extruded loops from hybrid MD—MC simulations with *v*_*ex*_ ≈ 0.6 kbp per *τ*_0_ (*κ* ≈ 0.3) starting from a relaxed polymer chain with unperturbed fractal dimension *D* ≈ 2. A) Mean square distance between two loci extruded by the same cohesin domain separated by genomic distance *s* within a loop of genomic length *l* (see Eq. (8)). Each blue curve corresponds to a different loop length *l* = 40, 80, 120, …, 600 kbp. The red lines show the predicted scaling behavior. The abscissa and ordinate are scaled by 1.1*g*(*l*) and 1.1*ξ*^2^(*l*), respectively. *g*(*l*) and *ξ*^2^(*l*) were obtained from a simulation of relaxed chromatin *without* extrusion (see inset and SI for details). The factor of 1.1 shifts the crossover between scaling behaviors to *s*/[1.1*g*(*l*)] = 1 and 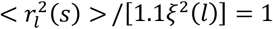 (dashed black lines). B) Average contact probability between loci separated by genomic length *s* within an extruded loop, averaged over loop lengths 800 ≤ *l* ≤ 1000. The dashed vertical line indicates the crossover to *P*(*s*) ∼ *s*^−3/4^, which is approximately *g*(1000 kbp) ≈ 60 kbp. The red lines show the predicted scaling behavior.

### Steady-state active loop extrusion without TAD anchors compacts chromatin on scales smaller than processivity

We now consider many cohesins that actively extrude in a steady state with average processivity *λ* and separation *d* without including TAD anchors. Four regimes of processivity and separation dictate chromatin compaction and loop nesting on genomic length scales shorter than the genomic entanglement length with activity *N*_*e,active*_ (see Fig. 7A and SI for details).

**Figure 7:**
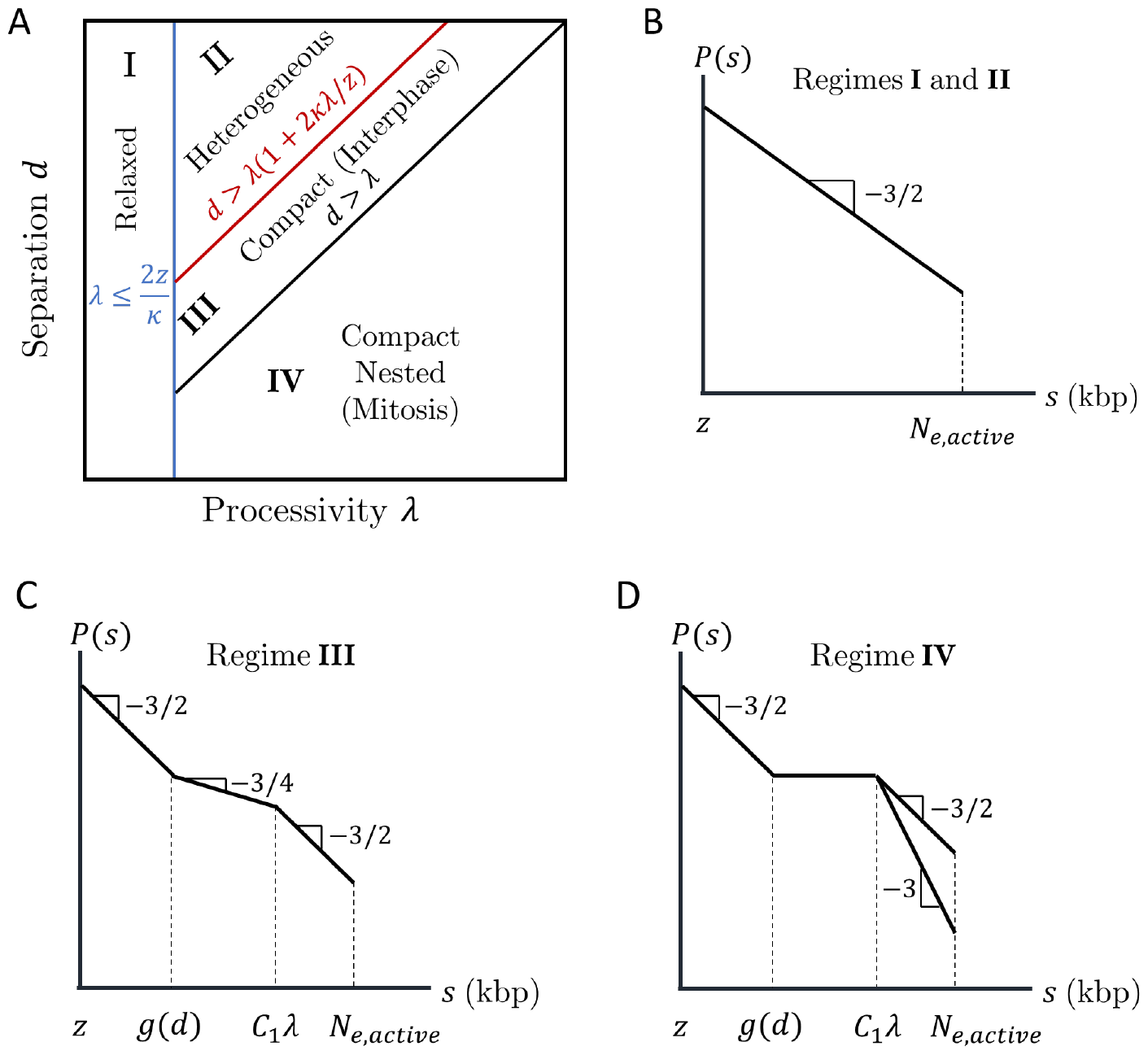
Four regimes of chromatin compaction and nesting. A) Diagram indicating the four regimes of chromatin compaction and nesting as functions of cohesin processivity and separation. B)—D) Schematic contact probabilities on log-log scales for the different regimes. In D), genomic separations of *C*_1_*λ* < *s* ≤ *N*_*e,active*_ may have *P*(*s*) ∼ *s*^−3/2^ or *P*(*s*) ∼ *s*^−3^ depending on whether cohesins can traverse each other.

Figures 7B—D show schematic plots of contact probabilities *P*(*s*) for each regime up to the entanglement genomic length. For all regimes, *P*(*s*) ∼ *s*^−3/2^ (*γ*_1_ ≈ 3/2) on short genomic length scales. In Regime I, the average processivity *λ* ≤ 2*g*_*min*_ ≈ 2*z*/κ is small enough for entire loops to relax during their extrusion process. In Regime II, loops are long enough to be compact, but have time to fully relax before another cohesin binds with *d* > *λ*(1 + 2*κλ*/*z*). The system is heterogeneous with compact and relaxed sections; however, the average chromatin conformation is relaxed. In Regime III, a loop extruded by cohesin only partially relaxes before another cohesin binds to this unrelaxed section with *λ*(1 + 2*κλ*/*z*) ≥ *d* > *λ*. In a steady state, cohesin separation controls the largest chromatin section that relaxes such that

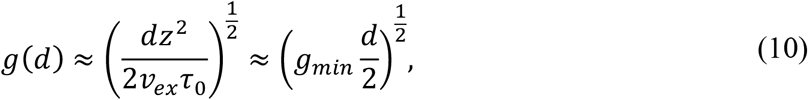

where *d*/(2*v*_*ex*_) is the average time between two cohesins binding within a chromatin section of length *λ*. Chromatin sections with shorter genomic lengths can fully relax before perturbation by another cohesin; chromatin sections with longer genomic lengths do not have enough time to relax. There is no loop nesting, but extrusion compacts the section from *D* ≈ 2 to *D* ≈ 4 such that *γ*_2_ ≈ 3/4 for *g*(*d*) ≤ *s* ≤ *C*_1_*λ*, where *C*_1_ is a constant on the order of unity. Extruding cohesins rarely simultaneously bind loci separated by much longer than *λ*; thus, for longer genomic lengths, chromatin is a random walk with *D* ≈ 2 of compact, extruded sections each of which has *D* ≈ 4 such that *γ*_3_ ≈ 3/2 (see Fig. 2D).

In Regime IV, *λ* ≥ *d* indicates significant loop nesting and the genomic section compacts into a globule with almost constant *P*(*s*) for *g*(*d*) ≤ *s* ≤ *C*_1_*λ*. On longer genomic length scales, chromatin could either be a random walk of compact sections as in Regime III or elongate due to excluded volume interactions between compact sections, which depends on whether cohesins can traverse each other. This regime can produce bottlebrush-like structures relevant to mitotic chromosomes formed by condensins and will be explored in future work.

There are too few cohesins per genomic length in Regimes I and II to compact chromatin. In Regime IV, chromatin adopts a nested structure that could form a bottlebrush not observed in interphase. Without entering the bottlebrush regime, chromatin compaction in Regime III is maximized when cohesin processivity and separation are approximately equal (by minimizing *g*(*d*) in Eq. (10) and maintaining *λ* ≤ *d*). Indeed, cohesin processivity and separation in mammalian cells are both predicted to be on the order of 200 kbp with limited nesting (7, 11, 51). The remainder of this work focuses on Regime III.

### TAD anchors can limit the range of extrusion-induced compaction

Compaction can also be limited by TAD anchors; since CTCF stops active loop extrusion, cohesins cannot directly bridge two loci in different TADs separated by lengths much larger than average TAD genomic length 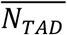 in the genomic section of interest. In this case, chromatin resembles a random walk of compact TADs with contact probabilities returning to *P*(*s*) ∼ *s*^−3/2^ for 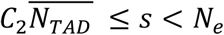 where *C*_2_ is a constant on the order of unity.

We simulate steady-state active loop extrusion on a 1 Mega-base pair (Mbp) chromatin section with and without TAD anchors with *λ* ≈ *d* ≈ 200 kbp and *v*_*ex*_ ≈ 0.3 kbp per *τ*_0_ (*κ* ≈ 0.15). CTCF sites were placed such that there were five consecutive TADs each with a genomic length of 200 kbp (see Fig. 8A). The simulated *P*(*s*) curves (see Fig. 8B) are consistent with the expected scaling behavior depicted for Regime III in Figure 7C. Without TADs, the crossover between *γ*_2_ ≈ 3/4 and *γ*_3_ ≈ 3/2 occurs at approximately *s* ≈ 400 kbp, suggesting that the constant *C*_1_ in Fig. 7C is approximately 2. With TAD anchors, the crossover occurs at the average TAD length 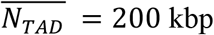. The peaks at *s* = 200 kbp and *s* = 400 kbp are due to the periodicity introduced by consecutive TADs.

**Figure 8:**
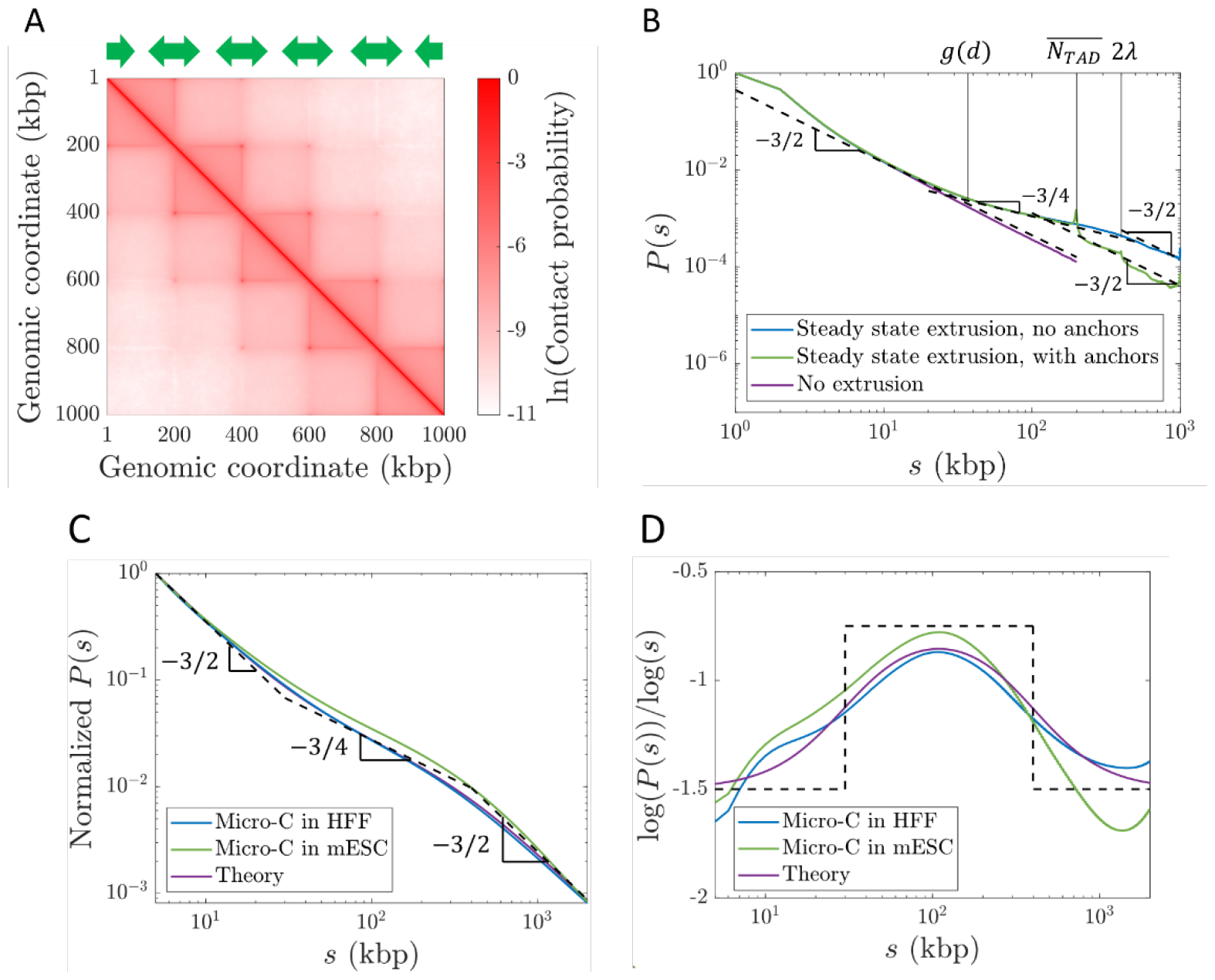
Contact probabilities from simulations with steady-state extrusion and public Micro-C data. A) Contact map from a simulation with 1000 kbp. CTCF sites were placed according to the green arrows such that there were five consecutive TADs of 200 kbp each. B) Contact probabilities from simulations with and without TADs. Dashed black lines show power laws *P*(*s*) ∼ *s*^−3/2^ and *P*(*s*) ∼ *s*^−3/4^. C) Genome-averaged contact probabilities normalized to *P*(5 kbp) = 1 from Micro-C in HFF (blue, (26)) and mESC (green, (32)) cells at 1 kbp resolution compared to theoretically expected scaling behavior (black dashed lines) and a smoothed theoretical crossover function (purple) *P*(*s*) ≈ *s*^−3/2^[1 + (*s*/30)^2^]^3/8^[1 + (*s*/400)^2^]^−3/8^. Fractal dimension is predicted to cross over from 2 to 4 at 30 kbp, and back to 2 at 400 kbp. D) Slopes on a log-log scale of curves in C).

### Experimental data are consistent with the predicted contact probability scaling

Next, we compare our theory with publicly available Micro-C data. We expect *P*(*s*) to follow Regime III in Figure 7C with crossovers between *γ*_1_ ≈ 3/2 and *γ*_2_ ≈ 3/4 at *s* ≈ *g*(*d*) and between *γ*_2_ ≈ 3/4 and *γ*_3_ ≈ 3/2 at 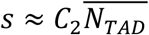. As discussed in the “Model Description” and SI, we choose an extrusion ratio of *κ* ≈ 0.2, indicating fast extrusion relative to chromatin relaxation within the range of observed chromatin loci mobilities. We reason that using genome-wide Micro-C data smooths out local variations in the lengths and genomic locations in TADs. To predict genome-wide *P*(*s*) in mammalian cells, we use 30 kbp as the crossover location between *γ*_1_ and *γ*_2_ and 400 kbp as the crossover location between *γ*_2_ and *γ*_3_ (using *C*_2_ ≈ *C*_1_ = 2). In Figure 8C we plot this predicted *P*(*s*) scaling (dashed black lines), a corresponding smoothed crossover function (purple curve), and two examples of genome-wide contact probabilities from Micro-C data (blue and green curves). Figure 8D shows that the slopes on a log-log scale of the two experimental data sets are consistent with our model, suggesting that our choice of *κ* ≈ 0.2 is reasonable. This scaling behavior is also consistent with other Hi-C and Micro-C experiments (5, 8, 25–27, 29–32). We note that specific genomic sections may have different average TAD lengths 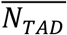, which could shift the crossover between *γ*_2_ and *γ*_3_.

### Active loop extrusion kinetically suppresses overlaps and dilutes entanglements

Another consequence of chromatin compaction due to active extrusion is the reduction of overlap between neighboring chromatin sections and entanglement dilution. We define the overlap parameter *O*(*s*) as the number of chromatin sections with genomic length *s* with the same mean square spatial size < *r*^2^(*s*) > that share the same approximately spherical volume with diameter ≈< *r*^2^(*s*) >^1/2^

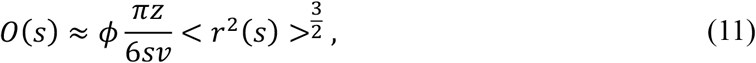

where *ϕ* ≈ 0.06 − 0.4 is the average chromatin volume fraction in the nucleus and *v* is the physical volume occupied by a locus, which for *z* ≈ 2 kbp we estimate to be ≈ 7.5×10^3^ nm^3^ (see SI). If a chromatin section with genomic length *s* has a fractal dimension of *D* between length scales *b* and < *r*^2^(*s*) >^1/2^, the overlap parameter is *O*(*s*) ≈ [*πb*^3^/(6*v*)]*ϕ*(*s*/*z*)^3/*D*−1^. Entanglements occur above a critical overlap parameter *O*_*KN*_ on the order of 10—20 as conjectured by Kavassalis and Noolandi (60). If chromatin has *D* ≈ 2 without loop extrusion, the entanglement genomic length is approximately *N*_*e,passive*_ ≈ 100 kbp for an average chromatin volume fraction *ϕ* ≈ 0.15 and *O*_*KN*_ = 10. Previous estimates suggest an entanglement genomic length without extrusion nor extensive looping of 50—100 kbp (33, 42–44). Our estimate of *N*_*e,passive*_ ≈ 100 kbp is also consistent with Hi-C data showing a contact probability scaling of *P*(*s*) ∼ *s*^−0.94^ for 100 kbp < *s* < 500 kbp in cohesin-depleted cells (25).

With active loop extrusion, the overlap parameter *decreases* with genomic length when *D* ≈ 4, which we predict to occur on genomic length scales between ≈ *g*(*d*) and 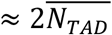 (see Fig. 9). The mean square size of genomic sections longer than *g*(*d*) is smaller with active loop extrusion compared to the passive case with random walk statistics up to entanglements. As such, longer genomic sections are required to reach *O*_*KN*_ such that

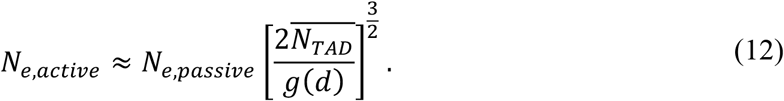

**Figure 9:**
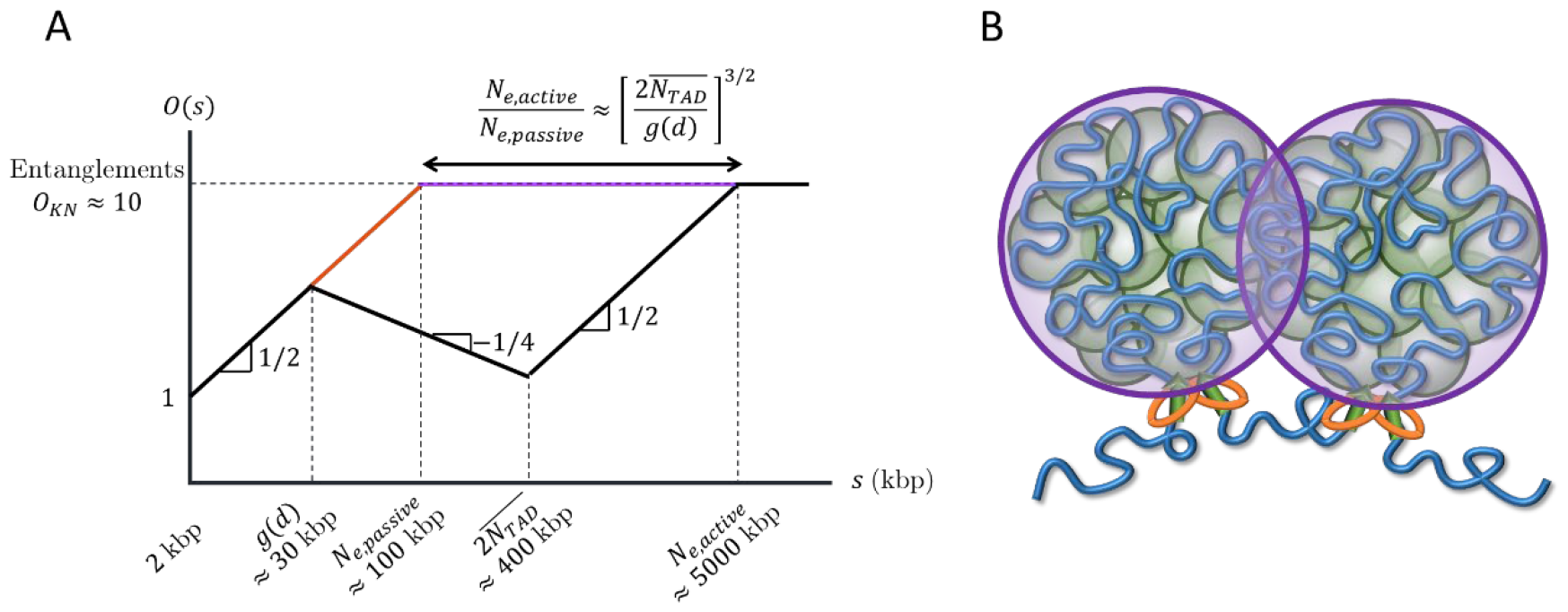
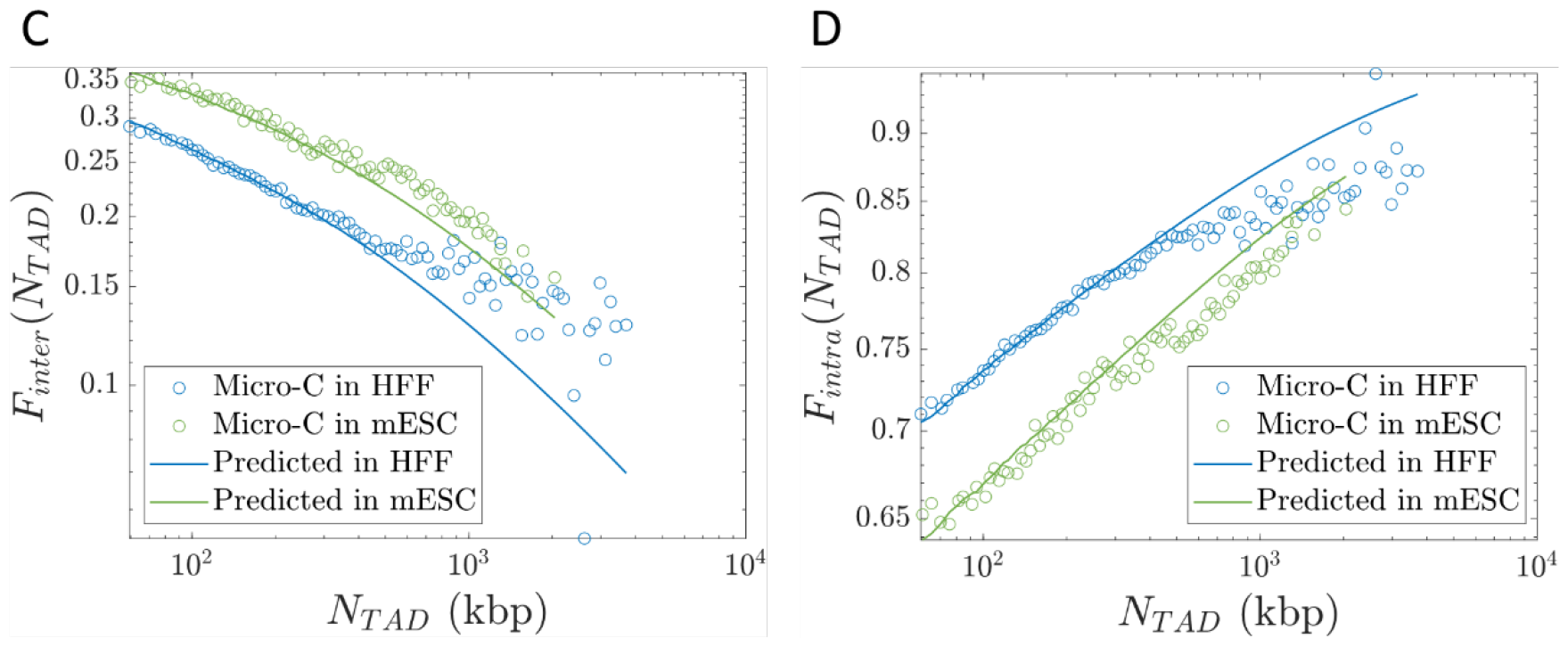
Overlaps between chromatin sections. A) Schematic plot of the overlap parameter *O*(*s*) for active loop extrusion (black), random walks (red), and fractal loopy globules (purple) as a function of genomic length on a log-log scale. The black and purple curves merge for *s* ≥ *N*_*e,active*_. The horizontal dashed line indicates the onset of entanglements. We use the volume fraction of chromatin in the nucleus *ϕ* = 0.15 and *O*_*KN*_ = 10. B) Chromatin sections within TADs (green circles) overlap and have increased contact probabilities. Neighboring TADs (purple circles) are mostly segregated apart from a narrow interface. C) Inter-TAD fraction of all intra-chromosomal contacts made by loci in a TAD as a function of TAD length. D) Intra-TAD fraction of all intra-chromosomal contacts made by loci in a TAD as a function of TAD length.

With 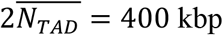 and *g*(*d*) = 30 kbp, active loop extrusion can increase *N*_*e*_ by up to a factor of approximately 50. If the passive entanglement genomic length is 50—100 kbp, active loop extrusion could dilute entanglements up to 2.5—5 Mbp. In Table S3 we provide ranges for the entanglement genomic lengths for different chromatin volume fractions *ϕ* and *O*_*KN*_.

### Active loop extrusion segregates TADs and enhances intra-TAD contacts

The increase in fractal dimension to *D* ≈ 4 due to active loop extrusion is consistent with TAD segregation. Loop extrusion decreases TAD overlap compared to random walks with *D* ≈ 2 and fractal loopy globules with *D* ≈ 3. In the fractal loopy globule model, the overlap parameter remains constant at *O*_*KN*_ ≈ 10 above the entanglement genomic length, which with *N*_*e,passive*_ ≈ 50 − 100 kbp indicates relatively strong overlap of genomic loci between TADs. If this were the case, regulatory elements would frequently affect promoters in different TADs. On the other hand, the decrease in overlap parameter with active loop extrusion by a factor of [*s*/*g*(*d*)]^3/2^ for 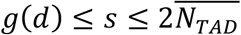 (see Fig. 9) indicates that genomic contacts between TADs are suppressed compared to contacts within TADs. Active loop extrusion suppresses the number of overlapping TADs compared to the passive case (without extrusion) by up to a factor of 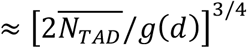. For 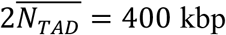 and *g*(*d*) = 30 kbp, this reduction factor is approximately 7. For *ϕ* = 0.15 and *O*_*KN*_ = 10, the overlap parameter at 400 kbp reduces from ≈ 16 to ≈ 3. Note that because our theory is on a scaling level and the degree of overlap between genomic sections depends on the shape of their pervaded volumes (*e*.*g*., spherical, or ellipsoidal), our estimates can differ from real biological systems by a factor of two.

Our model suggests that while contacts between TADs are suppressed, contacts between sections within the same TAD are enhanced. Loci in each TAD mostly contact other loci separated by less than 400 kbp within the same TAD, apart from a narrow zone of interaction at the interface between neighboring TADs (see Fig. 9B). Future simulations will further investigate this effect. Analysis of Micro-C data shows that out of all intra-chromosomal contacts made by loci in a TAD of length *N*_*TAD*_, the inter-TAD fraction *F*_*inter*_(*N*_*TAD*_) (contacts between loci in the TAD and other loci on the same chromosome but not in the same TAD) is ≲ 0.35 and monotonically decreases with TAD length (see Fig. 9C). Conversely, the intra-TAD fraction *F*_*intra*_(*N*_*TAD*_) (contacts between loci in the TAD and other loci within the same TAD) is ≳ 0.65 and monotonically increases with TAD length (see Fig. 9D). Predictions using our theory agree with Micro-C data in HFF and mESC cells (see Figs. 9C—D). See SI for details. As such, active loop extrusion facilitates colocalization of promoters and regulatory elements within the same TAD while limiting erroneous contacts between TADs.

### Active loop extrusion extends chromatin adjacent to loops

Cohesin pulls on chromatin directly adjacent to loops, causing two tension fronts that propagate into the unextruded sections (see Fig. 10A). We use the term “leg” to denote the largest chromatin section that adapts to this pulling force. Active extrusion suppresses the relaxation modes of the legs, as cohesin translocation is ballistic. Cohesin translocation rather than polymer relaxation controls leg conformation. Cohesin reels in chromatin legs, each with genomic length *g*_*leg*_(*t*), and unravels their conformations. Loci separated by more than *g*_*leg*_(*t*) past cohesin are not “aware” of extrusion. The mean square size of a 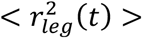 is approximately the mean square size of a genomic length *v*_*ex*_*t* + *g*_*leg*_(*t*) before extrusion started such that 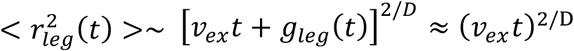 with corrections due to partial relaxation. Each leg is extended such that its genomic length is 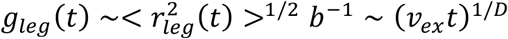 (see SI).

**Figure 10:**
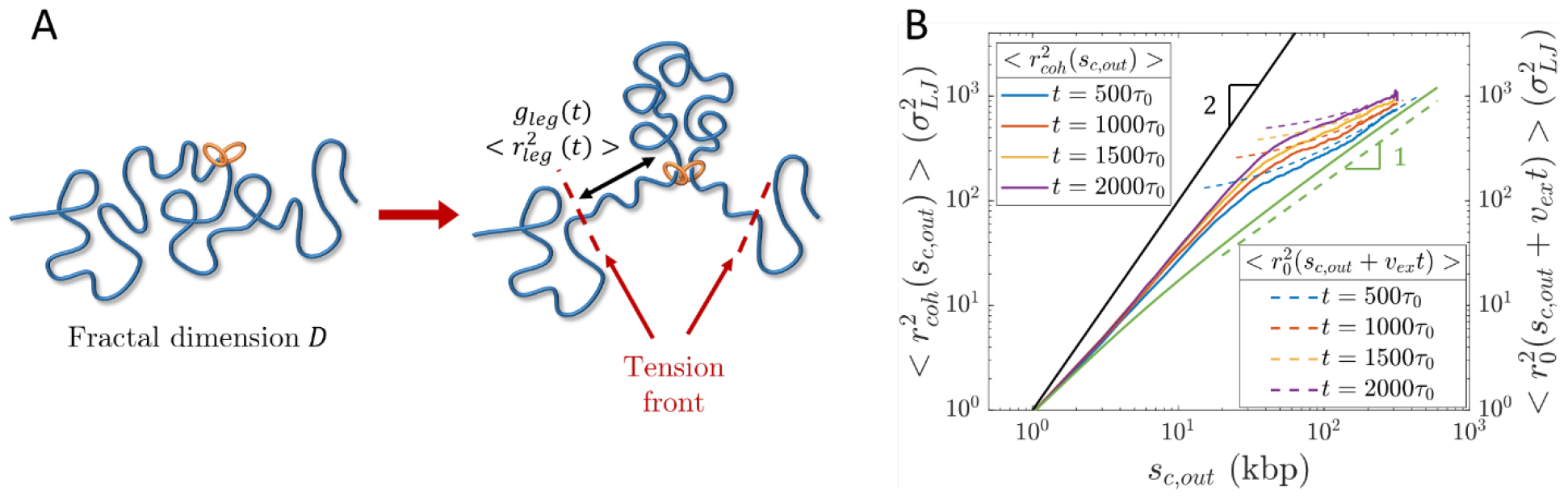
Conformation of legs produced by active loop extrusion. A) Schematic of legs, each with genomic length *g*_*leg*_ (*t*) ∼ (*v*_*ex*_*t*)^1/*D*^ and mean square size 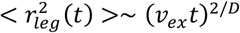. The left-hand schematic shows the initial chromatin conformation with fractal dimension *D* at the time of cohesin binding. The right-hand schematic shows leg extension at time *t*. B) Mean square distance between cohesin and a locus *s*_*c,out*_ away outside of its extruded loop at different times after binding (solid lines) compared to 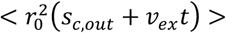, the unperturbed mean squared sizes of sections with genomic lengths *s*_*c,out*_ + *v*_*ex*_ *t* (dashed lines), from hybrid MD—MC simulations of single cohesins actively extruding a relaxed polymer chain (*D* ≈ 2) with *v*_*ex*_ ≈ 0.3 kbp per *τ*_0_ (*κ* ≈ 0.15). The solid black line is the predicted limiting behavior for full polymer extension. The solid green curve is the unperturbed mean square sizes of sections with genomic length *s*_*c,out*_. The dashed green line is the predicted scaling behavior of these unperturbed sizes without extrusion.

Consider 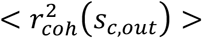, the mean square distance between cohesin and a locus *s*_*c,out*_ base pairs outside of the loop. The limiting behavior is a straight array of loci with 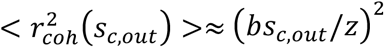. For genomic distances longer than the leg length, 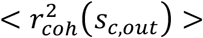 is approximately equal to 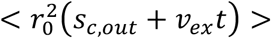, the mean square distance between the cohesin binding site and the locus of interest before extrusion started. Hybrid MD—MC simulations of a single cohesin extruding a loop on an initially relaxed polymer chain are consistent with this result (see Fig. 10B).

### Active loop extrusion causes anomalous dynamics of actively extruding cohesins and chromatin loci

The two tension fronts (one tension front produced per leg) localize an actively extruding cohesin in the volume between them. A cohesin’s trajectory follows the midpoint between the two tension fronts (see Fig. 11A) and fluctuates around it by ≈ *b*^2^(*t*/*τ*_0_)^1/2^. Hybrid MD—MC simulations agree with this result (see Figs. 11B and S2). Multiple unnested cohesins attract each other in space because they exert tension on the same intervening genomic sections. After the cohesins meet, they travel together in space; this effect is more pronounced if the cohesins cannot traverse one another (see SI). This “self-focusing” effect of neighboring cohesins further maintains the compact structure of actively extruded chromatin. Future work will explore additional details, including the nested cohesin case.

**Figure 11:**
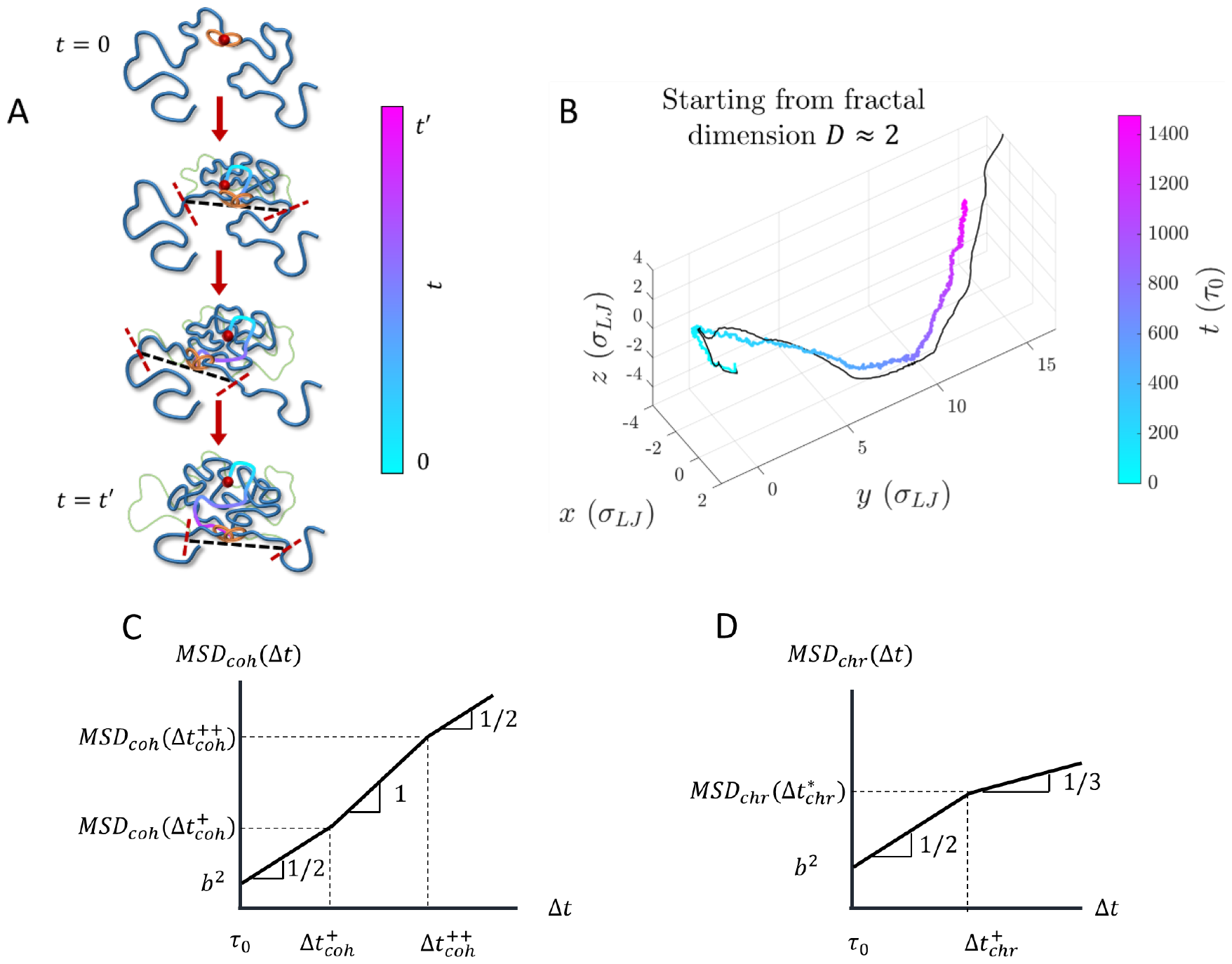
Dynamics of cohesin and chromatin loci during active extrusion. A) Cohesin trajectory (gradient curve) after binding to chromatin at its binding site (red circle) at time *t* = 0. The thick blue curve represents chromatin at a given moment, and the thin green curve represents the initial conformation. The gradient from cyan to magenta indicates time. Dashed black lines connect the tension fronts (indicated by dashed red lines) at different times, which propagate through the section with time. B) Average cohesin trajectory (gradient curve) from a fixed binding site and the corresponding smoothed trajectory of midpoints between two tension fronts (black curve) from 200 simulations starting from the same initial conformations with *v*_*ex*_ ≈ 0.3 kbp per *τ*_0_ (*κ* ≈ 0.15) starting from *D* ≈ 2. C) MSD of actively extruding, chromatin-bound cohesins on a log-log scale. D) MSD of chromatin loci affected by active extrusion on a log-log scale.

The mean square displacement (MSD) of cohesin *MSD*_*coh*_ (Δ*t*) is coupled on short time scales to the Rouse modes of the smallest relaxed chromatin section *g*_*min*_ and is proportional to Δ*t*^1/2^ for lag times 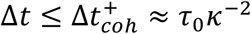. Cohesin trajectories then follow tension fronts dictated by the chromatin conformation such that *MSD*_*coh*_ ∼ Δ*t*^2/*D*^. Upon steady-state extrusion, the cohesin MSD for 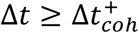 crosses over between scaling behaviors of ∼ Δ*t* and ∼ Δ*t*^1/2^ at 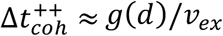 (see Fig. 11C and Fig. S6). 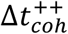 is approximately the time it takes a cohesin domain to extrude a genomic length *g*(*d*) in the process of active loop extrusion. Both crossover times 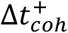 and 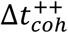 depend on the *D*_*app*_ of the chromatin section of interest, where the chromatin locus MSD is *MSD*_*chr*_(Δ*t*) ≈ *D*_*app*_Δ*t*^1/2^ (see SI). For *κ* ≈ 0.2, a locus discretization of *z* = 2 kbp per locus, and locus size *b* = 50 nm, we predict 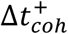 to be on the order of 100 seconds and Δ*t*^++^ to be on the order of 300 seconds.

In the Rouse model of polymer dynamics, the MSD of monomers in a polymer with fractal dimension *D* scales as ∼ Δ*t*^2/(*D*+2)^ (34). The change in chromatin fractal dimension due to active loop extrusion on scales below cohesin processivity causes *MSD*_*chr*_(Δ*t*) to cross over between scaling behaviors of ∼ Δ*t*^1/2^ and ∼ Δ*t*^1/3^ (see Fig. 11D and Fig. S7). The crossover time 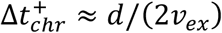 does not depend on *D*_*app*_ (see SI). In the absence of other activity for *d* ≈ 200kbp and *v*_*ex*_ ≈ 0.1 kbp/s, we predict the crossover to occur at lag times on the order of tens of minutes. Consistent with this prediction, the two-point MSD of the relative position of two loci has been observed to deviate from a ∼ Δ*t*^1/2^ scaling to a weaker time dependence after ≈15 minutes, though this could be due to the MSD approaching twice the mean square size of the intervening genomic section (54). In the SI we discuss the MSD of chromatin loci for longer lag times.

## DISCUSSION

### Summary

We present a theory and accompanying hybrid MD—MC simulations of chromatin organization and dynamics during interphase driven by active loop extrusion. Extrusion produces compact loops composed of overlapping relaxed chromatin sections (see Fig. 5). We show that within extruded loops, chromatin conformation can have the fractal dimension of two on length scales smaller than the sizes of these relaxed sections. The loops are much more compact with fractal dimensions of four on larger length scales, with the crossover between these regimes at chromatin strands of ≈ 30 kbp. We suggest that conformations of chromatin strands longer than TADs are random walks of looped sections with topological interactions appearing on scales of several Mbps. The predicted contact probability scaling behavior is consistent with publicly available experimental data (see Figs. 8C—D). Our model suggests that active loop extrusion increases the entanglement genomic length of chromatin by almost two orders of magnitude and segregates TADs (see Fig. 9). We also predict the MSD for both actively extruding chromatin-bound cohesins (Fig. 11C) and chromatin loci (Fig. 11D). This work provides a theoretical explanation for the compact, largely unentangled structure of chromatin during interphase.

### Active loop extrusion is an effective mechanism for transcriptional regulation

Contacts between loci that may be separated by hundreds of kbp can up- or down-regulate genes (1). Active loop extrusion is an effective way of bringing genomic loci within physical proximity and segregating TADs, which may contribute to their insulating properties, consistent with TAD segregation observed with microscopy (39–41). At genomic separations of ≈ 400 kbp, active loop extrusion increases contact probabilities by factors of approximately 7 and 3 compared to random walks and fractal loopy globules, respectively. This model ensures that *cis*-regulatory elements mostly encounter promoters within the same TAD separated by up to 400 kbp due to the local minimum of overlap parameter on the order of unity at this scale (see Fig. 9). Another consequence of chromatin compaction is the facilitation of transcription factor binding to chromatin (TF binding) within TADs. Active loop extrusion increases the volume fraction of a TAD within its pervaded volume. While a TF explores this volume, the frequency of encounters with its binding sites in the TAD increases. This is consistent with experimental evidence that cohesin depletion can impair TF binding, particularly for inducible TFs like the glucocorticoid receptor (61), by reducing the TF target search efficiency (32).

### Cohesin affects large-scale chromatin organization and dilutes entanglements

Cohesin-mediated chromatin compaction on scales shorter than 400 kbp reduces spatial overlaps between genomic sections, thus suppressing genomic entanglements (see Fig. 9A). We predict the entanglement genomic length of chromatin with active loop extrusion to be approximately 5 Mbp. Our model is consistent with experiments showing that interphase chromatin is largely unentangled (45). Furthermore, recent work observed more disordered chromatin organization and anomalous genomic contacts on genomic length scales of several Mbps upon cohesin degradation (62), indicating that cohesin regulates chromatin conformation in a wide range of genomic lengths.

### TAD anchors can limit chromatin compaction

As seen in our computer simulations, TAD anchors can control the crossover between contact probability scaling exponents *γ*_2_ ≈ 3/4 and *γ*_3_ ≈ 3/2. Thus, the *P*(*s*) profile can vary for genomic sections with different distributions of TADs. If the average TAD length 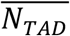 is greater than *λ*, we expect a negligible impact on *P*(*s*) compared to without TAD anchors (*i*.*e*., Fig. 7C and the blue curve in Fig. 8B). Consecutive TADs each with length 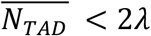 truncate the regime associated with *γ*_2_ such that the crossover to *γ*_3_ shifts to 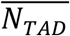 (see Fig. 8B). If 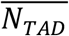 is shorter than both the cohesin processivity and separation, the location of the first crossover between *γ*_1_ ≈ 3/2 and *γ*_2_ ≈ 3/4 can also shift to 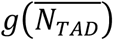 (see Eq. (7)). This crossover could be impacted by how long TAD anchors are held together by cohesins compared to the average residence time of unimpeded cohesin.

### Active loop extrusion parameters control contact probabilities

Extrusion velocity, cohesin processivity, and cohesin separation modulate contact probabilities both within loops formed by single cohesins and within sections regulated by multiple cohesins. In both cases, the first crossover in *P*(*s*) between *γ*_1_ ≈ 3/2 and *γ*_2_ ≈ 3/4 can be tuned by extrusion velocity (see Eq. (10)). Faster extrusion decreases the genomic length of chromatin sections that can relax, which increases compaction within each loop. When multiple cohesins organize a genomic section, decreasing the average cohesin separation can also shift this crossover to shorter genomic length scales (see Eq. (10)), ideally by increasing cohesin binding frequency. More frequent cohesin binding results in shorter chromatin sections that relax between binding events. Cohesin processivity controls the crossover in *P*(*s*) between *γ*_2_ ≈ 3/4 and *γ*_3_ ≈ 3/2 (as well as TAD anchors, as discussed above). Longer processivities (achieved, for example, by extending cohesin residence times) increase the genomic separation between loci that can be held together by active extrusion. If experiments could develop fine control over the parameters of active loop extrusion in particular TADs, this work could help tune the contact probabilities between specific loci of interest.

### Model assumptions

One assumption of our model is that active loop extrusion is the main regulatory mechanism of chromatin organization. Recent studies have carefully examined the role of RNAPII in shaping chromatin organization, showing that specific chromatin loops and genomic sections are impacted by active transcription (27, 63–65). However, RNAPII degradation does not affect genome-wide contact probabilities *P*(*s*) on genomic length scales shorter than 10 Mbp (27, 64). Furthermore, transcription is known to occur through bursting kinetics, with inactive periods on the order of hours (18, 66, 67). As a result, although transcription and other processes may affect chromatin organization at specific genomic locations, we reason that on average, genome organization is predominantly shaped by cohesin-mediated active loop extrusion.

We also assume that the crossovers between scaling regimes in *P*(*s*) and overlap parameter *O*(*s*) are well-characterized by the average cohesin processivity and separation. The averages and distributions of these parameters could vary with genomic context, which could shift the crossover locations to different genomic length scales. Our computer simulations suggest that *P*(*s*) averaged over a 1 Mbp section is well described by the average *λ* and *d*. More detailed simulations will confirm whether this is true for *O*(*s*) as well. The crossover genomic lengths should not impact the asymptotic scaling behaviors of both *P*(*s*) and *O*(*s*) (*ex*., the *γ*_*i*_ exponents). Additionally, we assume that almost all contacts made by a given locus are intra-chromosomal. Some process(es) must ensure that chromosomes are confined to localized volumes since it is well-known that chromosomes segregate into territories (68). We leave the investigation of the phenomena that could lead to such segregation as an open question for future work.

### Comparison with other models

As discussed in the Introduction, the CPEL, FLG, and DFLA models for chromatin organization are also consistent with fractal dimensions *D* > 2 (44, 46–49). In contrast to these three equilibrium models, we directly couple active, ATP-dependent extrusion *kinetics* to chromatin conformation and dynamics. The fractal dimension *D* ≈ 4 between the scales of approximately 30 kbp and 400 kbp is a direct result of activity. With respect to CPEL, the looped sections with *D* ≈ 2 could be interpreted as analogous to relaxed sections with genomic length *g*(*d*) in our model. However, in CPEL the relaxed loop lengths are exponentially distributed, whereas in our model, the *extruded* loop lengths are exponentially distributed while the *relaxed* sections within an extruded loop are uniformly distributed (see SI). In contrast to FLG, our model explains the anomalous fractal dimension of *D* ≈ 4. We suggest that chromatin sections longer than *N*_*e,active*_ maintain constant overlap like in FLG (see Fig. 9). Finally, to compare with DFLA, the polymer sections between fixed obstacles with *D* ≈ 2 may be analogous to the relaxed sections with genomic length *g*(*d*) in our model.

### Conclusion

In conclusion, our model explains how active loop extrusion compacts individual chromatin loops, forms segregated TADs, and dilutes chromatin entanglements. This work suggests a crucial role of loop extruding proteins in maintaining effective regulation of transcription by distal elements. Future experiments informed by our model may be able to control contact probabilities within genomic sections of interest.

## MATERIALS AND METHODS

Hybrid MD—MC simulations were used to model active loop extrusion on linear polymer chains. Chains in theta-like solvent were simulated using the Kremer-Grest bead-spring model (69). A finite extensible nonlinear elastic (FENE) potential connected bonded beads. Non-bonded beads interact via a shifted and truncated Lennard-Jones pairwise potential. We represent cohesin as a switchable FENE bond that can move between binding partners. The FENE bond representing cohesin changes partners according to MC. The spatial trajectories of each bead were evolved in time by MD. Cohesin binding and translocation kinetics were updated by MC. MD was performed using the Large-scale Atomic/Molecular Massively Parallel Simulator (LAMMPS) package (70). MC steps were implemented in C and coupled to MD. For most simulations, chains were initiated as random walks and equilibrated for at least ten times their relaxation times before starting extrusion. At least 100 replicates were run of each condition for single extrusion cycle simulations. The steady-state simulations were run for at least 10 times the end-to-end vector autocorrelation decay time. Contact probabilities in steady-state simulations are consistent across different initial chain conformations (see SI). Experimental contact probability plots were extracted from Micro-C data using cooltools (71). See SI for extended methods.

## ACKNOWLEDGEMENTS AND FUNDING

MR acknowledges funding support by the National Science Foundation under grant number EFMA-1830957 and the National Institutes of Health under grant numbers P01-HL164320 and 5P01-HL108808.

## Notes

### Competing Interest Statement

The authors have declared no competing interest.

