## Supporting Information for "Activity-driven chromatin organization during interphase: compaction, segregation, and entanglement suppression"

#### Authors and Affiliations

Brian Chan[a] and Michael Rubinstein[a, b, c, d, e, 1]

[a] Department of Biomedical Engineering, Duke University, Durham, North Carolina, 27708, United States
[b] Thomas Lord Department of Mechanical Engineering and Materials Science, Duke University, Durham, North Carolina, 27708, United States
[c] Department of Physics, Duke University, Durham, North Carolina, 27708, United States
[d] Department of Chemistry, Duke University, Durham, North Carolina, 27708, United States
[e] World Premier International Research Center Initiative – Institute for Chemical Reaction Design and Discovery, Hokkaido University, Sapporo, 001-0021, Japan

**This file includes:**

Supporting text
Figures S1 to S13
Tables S1 to S3
SI References

**Table of contents:**

I. **Estimates of model parameters**
II. **Theory and extended data**
   a. Description of an extruded chromatin loop
   b. Steady-state extrusion regimes
   c. Entanglement suppression
   d. Description of legs and cohesin trajectories
   e. Mean square displacement of cohesins and chromatin loci
III. **Extended methods**
   a. Hybrid molecular dynamics – Monte Carlo (MD—MC) simulations
   b. Contact probability plots from Micro-C data
   c. Inter-TAD and intra-TAD contacts
IV. **Supplementary tables**
   a. Table S1: Description of notations used in main text.
   b. Table S2: Range of parameter estimates given a range of extrusion ratios.
   c. Table S3: Range of the estimated passive and active entanglement genomic lengths.
V. **SI references**

### I. Estimates of model parameters

Locus discretization $z$, $b$, $v$, and $\phi_z$

We discretize chromatin into loci each with genomic length $z$ and spatial size $b$. We assume that in the absence of activity (no extrusion), chromatin conformation follows random walk statistics on length scales longer than $b$ such that the root-mean-square size of a section with genomic length $s$ is proportional to $s^{1/2}$. The loci size $b$ is assumed to be equal to or larger than the Kuhn segment of human chromatin. The exact value of Kuhn length of chromatin is not known with the estimates ranging from one to tens of kbp corresponding to tens to hundreds of nanometers (nm) (1–5). The lower end of this range represents the 10-nm fiber model of chromatin, while the upper end represents the 30-nm fiber model.

As the 10-nm fiber model is more relevant to interphase chromatin in live cells (6, 7), we choose a discretization of $z = 2$ kbp. The locus size is approximately $b = 50$ nm using a linear density of ≈40 bp/nm (1). This locus discretization is on the order of or larger than Kuhn length estimates for human chromatin (3–5). $b = 50$ nm is five times shorter than the length of a 2 kbp straight array of 10 nucleosomes and 500 bp of double stranded linker DNA. Each nucleosome has a diameter of ≈10 nm and each base pair has length of ≈0.34 nm. The physical space occupied by each locus is approximately

$$v \approx 10\left(\frac{4}{3}\pi 5.5^3 + 50 * 0.34\pi\right)\mathrm{nm}^3 \approx 7.5\mathrm{x}10^3\mathrm{nm}^3\,, \quad \text{(S1)}$$

where we use approximately spherical nucleosomes with radii of ≈5.5 nm each with ≈150 bp, and ≈50 bp of double stranded DNA with diameter of ≈2 nm. The volume fraction of chromatin (double stranded DNA and nucleosomes) within one 2 kbp locus is approximately $\phi_z \approx v/[4\pi(b/2)^3/3] \approx 0.1$.

Locus diffusion time $\tau_0$ and extrusion ratio $\kappa$

As described in the main text, chromatin locus dynamics typically exhibit mean square displacements $MSD_{chr}(\Delta t) \approx D_{app}\Delta t^{1/2}$ with $D_{app}$ between ≈$10^{-3}$ μm$^2$ s$^{-1/2}$ and ≈$10^{-2}$ μm$^2$ s$^{-1/2}$ (8–13). Using Eq. (2) in the main text, the locus diffusion time is

$$\tau_0 \approx \left(\frac{b^2}{D_{app}}\right)^2, \quad \text{(S2)}$$

which for $b = 50$ nm, ranges between $\tau_0 \approx 0.06$ s and $\tau_0 \approx 6$ s. Using a typical cohesin processivity of $\lambda = 200$ kbp and a residence time of approximately $\tau_{res} \approx 25$ min, the extrusion velocity is $v_{ex} \approx 0.1$ kbp/s (14–20). The extrusion ratio $\kappa \approx v_{ex}\tau_0/z$ (Eq. (4) in the main text) thus ranges from $\kappa \approx 0.003$ to $\kappa \approx 0.3$. We choose $\kappa \approx 0.2$ corresponding to $\tau_0 \approx 4$ s and $D_{app} \approx 1.25\mathrm{x}10^{-3}$ μm$^2$ s$^{-1/2}$.

Average chromatin volume fraction in the nucleus

Human diploid cells have approximately $6.2\mathrm{x}10^9$ base pairs of chromatin (21). Assuming chromatin has on average 150 base pairs per nucleosome and 50 base pairs per linker DNA, the chromatin volume is approximately

$$v_{chrom} \approx 6.2\mathrm{x}10^9\mathrm{bp}\left(\frac{1\text{ nucleosome and 1 linker}}{200\mathrm{bp}}\right)\left(\frac{\frac{4}{3}\pi 5^3 + 50(0.34)(\pi)}{1\text{ nucleosome and 1 linker}}\right)\mathrm{nm}^3 \quad \text{(S3)}$$
$$\approx 2.3\mathrm{x}10^{11}\mathrm{nm}^3 \approx 230\mu\mathrm{m}^3.$$

Nuclei of typical human cells have diameters on the order of 10—20 μm (1, 22), meaning the average chromatin volume fraction ranges between $\phi \approx 0.06$ and $\phi \approx 0.4$. This estimate does not consider other proteins and molecules that may be complexed with chromatin.

### II. Theory and extended data

#### a. Description of an extruded chromatin loop

Theory

Consider a chromatin section with fractal dimension $D \approx 2$ on short genomic length scales in the absence of activity. As described above, we discretize chromatin into loci, each with spatial size $b$ and representing $z$ base pairs. A chromatin section with genomic length $s$ has mean square size $\approx b^2(s/z)$ in the absence of activity. The relaxation time of the section is $\approx \tau_0(s/z)^2$, where $\tau_0$ is the locus diffusion time. Let a cohesin bind locus $j$ at time $t' = 0$ and extrude with velocity $v_{ex}$ in units of genomic length per unit time (see Fig. 4 in the main text). The extrusion ratio $\kappa = \tau_0 v_{ex} z^{-1}$ is the number of loci extruded per diffusion time of a locus (Eq. (4) in the main text). $s_{xy}$ and $< r_{xy}^2(s_{xy}) >$ are the genomic distance and mean square spatial distance between loci $x$ and $y$ respectively, where the same cohesin domain extruded both loci and locus $x$ was extruded after locus $y$.

The cohesin extrudes locus $i$ at time $t_i$. Let us track the position of locus $i$ relative to cohesin at time $t$. $s_{ci} \approx (t - t_i)v_{ex}$ is the genomic distance between locus $i$ and cohesin (see Fig. S1a). For early times such that $(t - t_i)v_{ex} \leq g_{min} \approx z/\kappa$, locus $i$ is part of the smallest relaxed section of the extruded loop. For these early times, the relaxation time $\tau_0(s_{ci}/z)^2$ of a section with genomic length $s_{ci}$ is shorter than or equal to $t - t_i$. The mean square distance between $i$ and cohesin is the relaxed size of $s_{ci}$, approximately $\approx b^2(s_{ci}/z)$. For longer times, locus $i$ follows Rouse-like subdiffusive dynamics such that the mean square distance between $i$ and cohesin is $\approx b^2[(t - t_i)/\tau_0]^{1/2} \approx b^2[s_{ci}/(\kappa z)]^{1/2}$. Thus, the mean square distance between locus $i$ and cohesin as a function of the genomic length between them is

$$< r_{ci}^2(s_{ci}) > \approx \begin{cases} b^2\left(\frac{s_{ci}}{z}\right), & s_{ci} \leq g_{min} \\ b^2\left(\frac{s_{ci}}{\kappa z}\right)^{\frac{1}{2}}, & s_{ci} > g_{min} \end{cases}, \quad \text{(S4)}$$

see Fig. S1b. For $s_{ci} > g_{min}$, the mean square distance between locus $i$ and cohesin is *smaller* than the unperturbed size of a section with genomic length $s_{ci}$. This crossover broadens as the mobility of cohesin increases. With cohesin as the reference point, the extruded loop crosses over between fractal dimensions of $D \approx 2$ and $D \approx 4$ at genomic distances of $\approx g_{min}$ and mean square spatial sizes of $\approx b^2\, g_{min}/z \approx b^2/\kappa$. By replacing $s_{ci}$ with $l/2$, Eq. (S4) estimates the mean square size of an extruded loop with length $l$ because $l/2$ is the genomic distance between the cohesin and its binding site (see Eq. (6) in the main text).

The largest chromatin section that can relax around locus $i$ has genomic length on the order of

$$g_i \approx \left(\frac{s_{ci} z}{\kappa}\right)^{\frac{1}{2}} \quad \text{(S5)}$$

with mean square size

$$\xi_i^2 \approx b^2 \frac{g_i}{z} \approx b^2 \left(\frac{s_{ci}}{\kappa z}\right)^{\frac{1}{2}}. \quad \text{(S6)}$$

$g_i$ is found by equating the time since *i* was extruded $t - t_i \approx s_{ci}/v_{ex}$ to $\tau_0(g_i/z)^2$, the relaxation time of a section with genomic length $g_i$. The genomic length of these relaxed sections increases with $s_{ci}$. The largest relaxed section in a loop with genomic length *l* has genomic length

$$g(l) \approx \left(\frac{lz}{2\kappa}\right)^{\frac{1}{2}} \approx \left(\frac{g_{min} l}{2}\right)^{\frac{1}{2}}, \quad \text{(S7)}$$

where $l/2$ is the half-loop length, or the genomic distance between cohesin and the cohesin binding site. The mean square size of a section with genomic length $g(l)$ is

$$\xi^2(l) \approx b^2 \frac{g(l)}{z} \approx b^2 \left(\frac{l}{2\kappa z}\right)^{\frac{1}{2}} \approx b^2 \left(\frac{g_{min} l}{2}\right)^{\frac{1}{2}}. \quad \text{(S8)}$$

In Figure S1c, we sketch the genomic lengths of the relaxed sections in a loop of length $l$ as a function of $s_{ci}$. Consider two loci *i* and *m* extruded by the same cohesin domain where locus $i$ was extruded before locus $m$ (see Fig. S1a). The genomic distance between the two loci is $s_{mi} \approx (t_i - t_m)/v_{ex}$. The mean square spatial distance between *i* and *m* is approximately

$$< r_{mi}^2(s_{mi}) > \approx \begin{cases} b^2 \left(\frac{s_{mi}}{z}\right), & s_{mi} \le g_i \\ b^2 \left(\frac{s_{ci}}{\kappa z^3}\right)^{\frac{1}{4}} s_{mi}^{\frac{1}{2}}, & s_{mi} > g_i \end{cases}, \quad \text{(S9)}$$

where $s_{ci}$ is the genomic distance between locus $i$ and cohesin (see Fig. S1d). The mean square distance for $s_{mi} \le g_i$ is the relaxed distance, while the mean square distance for $s_{mi} > g_i$ is smaller than the relaxed distance. Equation (S9) is consistent with a crossover in fractal dimension between $D \approx 2$ and $D \approx 4$ at genomic distances of $\approx g_i$ and mean square spatial sizes of $\approx \xi_i^2$.

Now consider the mean square distance between the cohesin binding site locus *j* and a locus *i* within a loop of genomic length *l* (see Fig. S1a). The genomic distance between them is $s_{ij}$. The longest relaxed chromatin section around the cohesin binding site has a genomic length of approximately $g(l)$ (see Eq. (S7)). As such, the mean square distance between *j* and *i* is approximately

$$< r_{ij}^2(s_{ij}) > \approx \begin{cases} b^2 \left(\frac{s_{ij}}{z}\right), & s_{ij} \le g(l) \\ b^2 \frac{[s_{ij} g(l)]^{\frac{1}{2}}}{z}, & s_{ij} > g(l) \end{cases}, \quad \text{(S10)}$$

see Fig. S1e. As discussed in the main text, the mean square distance between any two loci separated by a genomic distance *s* extruded by the same cohesin domain crosses over between scaling behaviors of $\sim s^1$ to $\sim s^{1/2}$ (see Eq. (8) and Figs. 5D and 6A in the main text). The crossover occurs at genomic lengths on the order of $g(l)$ because the relaxed sections with length $g(l)$ have the most loci pairs, thus dominating the overall average.

Consider a genomic section with genomic length $s$ that is part of an extruded loop with genomic length $l$. The section's mean square size is $< r_l^2(s) >$ and is given by Eq. (8) in the main text. We define the section's self-volume fraction as

$$\phi_{self}(s) \approx \frac{\frac{s}{z} v}{\frac{4}{3}\pi \left[\frac{< r_l^2(s) >^{\frac{1}{2}}}{2}\right]^3}, \tag{S11}$$

where $v$ is the volume of one locus given by Eq. (S1). The numerator is the physical volume of chromatin in a genomic section of length $s$, and the denominator is the approximately spherical volume with diameter $< r^2(s) >^{1/2}$ spanned by the section. The self-volume fraction of one locus is $\phi_{self}(s = 2\text{kbp}) = \phi_z$, as described in Section I. The self-volume fraction at $s = l/2$ (half the loop length) is approximately $\phi_{self}(s = l/2) \approx \phi_z[lz^2 g(l)^{-3}/2]^{1/4}$ for loops longer than $g_{min}$. The self-volume fraction of the entire loop is then approximately two times $\phi_{self}(s = l/2)$ because the spatial size of the entire loop is the same as a section with length $s = l/2$ but with twice the volume of chromatin.

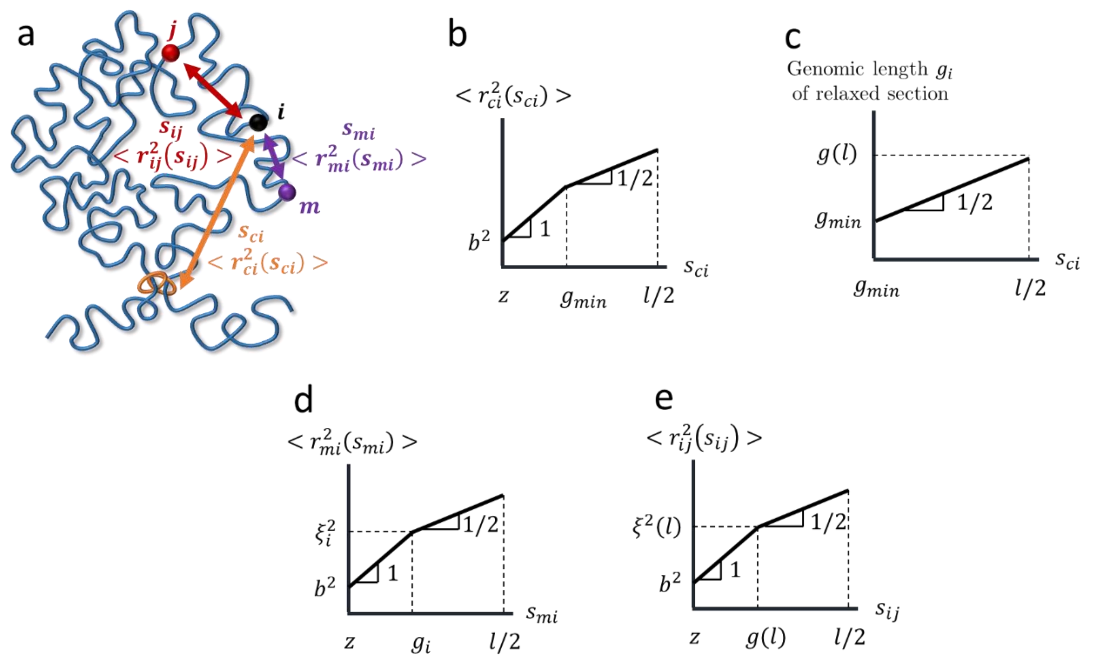


**Figure S1:** Internal structure of a single loop formed by active loop extrusion. All schematic plots are on log-log scales. a) Schematic of a chromatin loop (blue curve) actively extruded by cohesin (orange rings). Locus $j$ is the cohesin binding site. Loci $i$ and $m$ were extruded by the same cohesin domain. b) Schematic plot on a log-log scale of the mean square distance between cohesin and $i$ separated by genomic distance $s_{ci}$. Numbers next to lines indicate slopes on a log-log scale. c) Genomic length of the largest relaxed section around a locus $i$ that is a genomic distance $s_{ci}$ away from cohesin. d) Mean square distance between $i$ and $m$ that were extruded by the same cohesin domain and are separated by a genomic distance $s_{mi}$. The average is taken over all such pairs of loci in the loop. e) Mean square distance between $i$ and the cohesin binding site $j$ located a genomic distance $s_{ij}$ away.

Simulations

As described in the main text and in the "Extended methods" (Section III), to test our theory we run hybrid molecular dynamics – Monte Carlo (MD—MC) simulations of single extrusion cycles, where one cohesin extrudes one chromatin section once. Specifically, we examine the mean square distances between different loci within an extruded loop to validate Equations (S4), (S9), and (S10) as sketched in Figures S1b, d, and e, respectively. We also validate Equation (8) in the main text, sketched in Figure 5D in the main text. Instead of the sharp, clear power law scaling regimes in our theory, we expect broad crossovers over several decades in simulation and experimental data. The data in this section is averaged over 100 simulations of a single cohesin extruding a relaxed chromatin section with $D \approx 2$ using $v_{ex} \approx 0.6$ kbp per $\tau_0$ ($\kappa \approx 0.3$).

Figure S2 shows the mean square distances between cohesin and a locus $i$ within an extruded loop as a function of their genomic separation $s_{ci}$ (see Fig. S1a) from simulations with cohesins that are tethered to their initial points in space (immobile, purple curves) or free to move in space (mobile, blue curves). The black curve represents the mean square size of a section with length $s_{ci}$ without extrusion, while the red lines are the power laws predicted by our scaling theory. For small genomic separations $s_{ci}$, the curves with extrusion behave the same as without extrusion (black curve). The crossover between the unperturbed mean square distances and a power law $\sim s_{ci}^{1/2}$ for the immobile cohesin case is consistent with Eq. (S4) and Fig. S1b. The curves representing mobile cohesins have broader crossovers than their immobile counterparts.

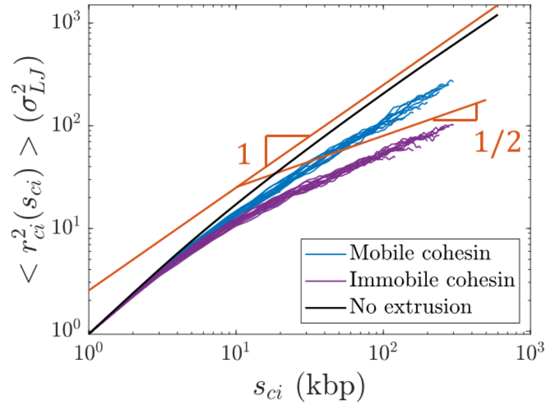


**Figure S2:** Mean square distance between cohesin and a locus $i$ within the loop as a function of their genomic separation $s_{ci}$, corresponding to Figure S1b. Each curve is a different loop length $l$ =40, 80, 120, …, 600 kbp.

Figure S3a shows the mean square distance $< r_{mi}^2(s_{mi}) >$ between two loci $m$ and $i$ within the same extruded loop as a function of the genomic separation between the two loci $s_{mi}$ (see Fig. S1a). Both loci were extruded by the same cohesin domain and locus $i$ was extruded before locus $m$. Each curve represents a different reference locus $i$, each with a different genomic distance to cohesin $s_{ci}$. The mean square distances for small separation $s_{mi}$ is approximately the same as the mean square size without extrusion (black curve). $< r_{mi}^2(s_{mi}) >$ then becomes smaller than the mean square size without extrusion. The crossover occurs at longer genomic lengths for longer $s_{ci}$, and corresponds to the largest relaxed section around each locus $i$ (green circles in Fig. 5C in the main text) with predicted genomic length $g_i$ (Eq. (S5)) and mean square size $\xi_i^2$ (Eq. (S6)).

To obtain $g_i$ and $\xi_i^2$ from simulation, we compare $MSD_{passive}$, the mean square displacement of beads in a simulated polymer *without* extrusion, to internal sizes $< r_{passive}^2(s) >$ *without* extrusion. $< {r_{passive}}^2(s) >$ is the mean square size of a section with $s$ beads averaged over different sections within the polymer chain. For each $s_{ci}$, we obtain $g_i$ from simulation as the largest section with $MSD_{passive}(s_{ci}/v_{ex}) \geq < r^2_{passive}(s) >$. This is the largest section that relaxes during time $s_{ci}/v_{ex}$, or the time since locus $i$ was extruded. The mean square size $\xi_i^2$ was obtained as $< r^2_{passive}(g_i) >$, the relaxed, unextruded sizes of sections with genomic length $g_i$. $g_i$ and $\xi_i^2$ were fitted to power laws such that $g_i \approx (2.8 \pm 0.3) s_{ci}^{(0.50\pm0.02)}$ and $\xi_i^2 \approx (4.52 \pm 0.02) s_{ci}^{(0.53\pm0.01)}$, close to the predicted scaling behavior of $\sim s_{ci}^{1/2}$ (see Eqs. (S5) and (S6)). Errors represent 95% confidence intervals.

Figure S3b shows the same data as Figure S3a, but with the abscissa and ordinate normalized by $g_i$ and $\xi_i^2$ from simulation, respectively, which collapses the data onto a universal curve. To find the crossover location, we fit the normalized data to two power laws: $< r^2_{mi}(s_{mi}) >/\xi_i^2 \approx (0.994 \pm 0.005) * (s_{mi}/g_i)^{(1.02\pm0.02)}$ for $0.5 \leq s_{mi}/g_i \leq 0.9$ and $< r^2_{mi}(s_{mi}) >/\xi_i^2 \approx (1.17 \pm 0.01) * (s_{mi}/g_i)^{(0.55\pm0.01)}$ for $2 \leq s_{mi}/g_i \leq 7$. The errors represent 95% confidence intervals. The intersection of the two curves is at $s_{mi}/g_i \approx 1.4 \pm 0.1$ and $< r^2_{mi}(s_{mi}) >/\xi_i^2 \approx 1.43 \pm 0.06$, where the errors represent the range of intersection between the simultaneous, functional bounds of the fitted power laws at 95% confidence levels.

The mean square distance between loci $m$ and $i$ in our simulations is consistent with a crossover between scaling regimes $\sim s^1_{mi}$ and $\sim s^{1/2}_{mi}$ at genomic length on the order of $1.4 g_i$ and mean square size on the order of $1.4\xi_i^2$, in agreement with Eq. (S9). Figure S3c shows the crossover genomic lengths $1.4 g_i$ (solid red curve) and mean square sizes $1.4\xi_i^2$ (solid blue curve), where $g_i$ and $\xi_i^2$ are obtained from simulation as described in the previous paragraph. We note that the fitted power law in the second regime for $2 \leq s_{mi}/g_i \leq 7$ has a stronger dependence on $s_{mi}$ than the theoretical scaling behavior of $\sim s^{1/2}_{mi}$. This could be due to a broad crossover between two scaling regimes and the relatively narrow range of fitting (less than a decade).

An alternative method to collapse the data in Figure S3a is to individually fit each blue curve to find where they deviate from the relaxed, unperturbed mean squared sizes (the black curve). We demonstrate that this method agrees with the method described in the previous paragraph for several of the blue curves in Figure S3a. For each curve, we fit the large $s_{mi}$ regime to a power law and find the intersection with the relaxed, unperturbed data. The red and blue circles in Figure S3c show the genomic lengths and mean square sizes of the intersection, respectively. The individually fitted crossover points are consistent with $1.4 g_i$ (solid red curve in Fig. S3c) and $1.4\xi_i^2$ (solid blue curve in Fig. S3c) as described in the previous paragraph.

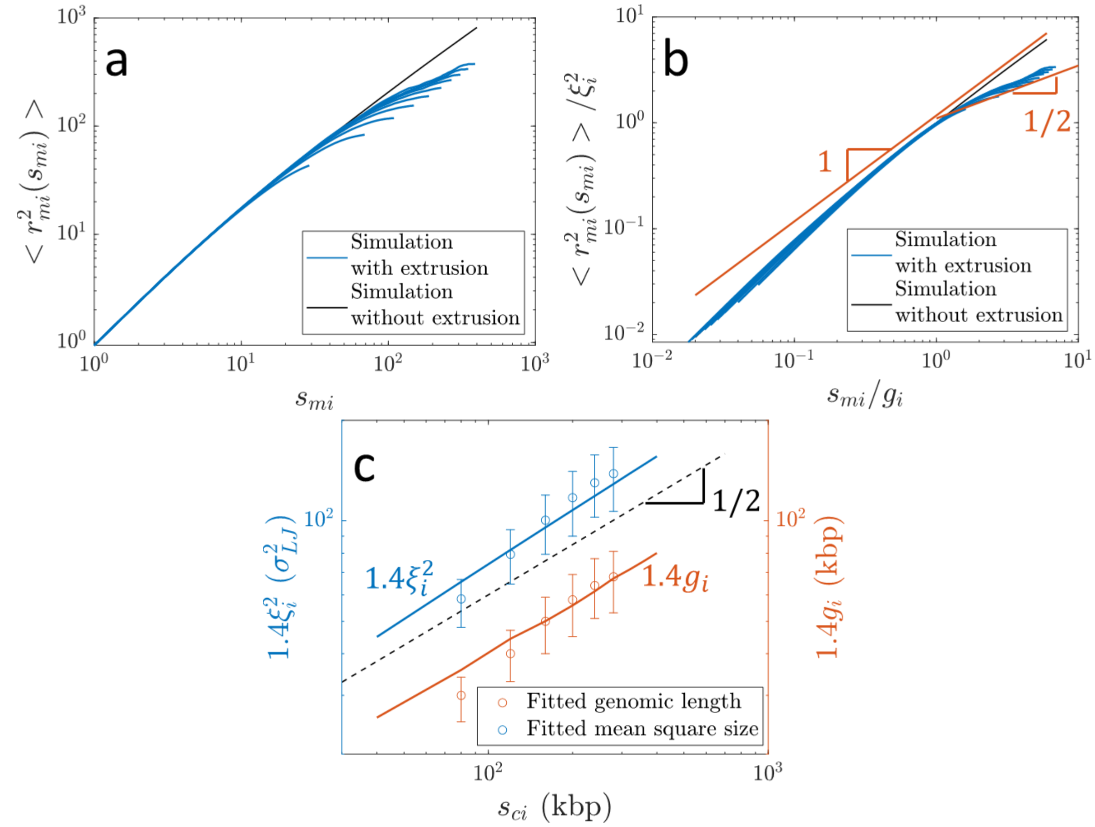


**Figure S3:** Mean square distance $< r_{mi}^2(s_{mi}) >$ between two loci $i$ and $m$ that were extruded by the same cohesin domain from hybrid MD—MC simulations (related to Fig. S1d). a) Comparison of $< r_{mi}^2(s_{mi}) >$ as a function of genomic separation $s_{mi}$ from simulations with and without extrusion. Each blue curve is for a different reference locus $i$ separated from cohesin by $s_{ci}$ =40, 80, 120, …, 400 kbp. b) The same data as a) with the abscissa and ordinate scaled by $g_i$ and $\xi_i^2$, respectively, obtained from simulation. The red lines are power laws $\sim (s_{mi}/g_i)^1$ and $\sim (s_{mi}/g_i)^{1/2}$. C) Crossover locations of $< r_{mi}^2(s_{mi}) >$ as a function of $s_{ci}$, the genomic distance between cohesin and locus $i$. The solid curves are the crossovers obtained by first collapsing all data *via* normalizing by $g_i$ and $\xi_i^2$ as determined from simulation. The circles are the crossovers obtained by individually fitting the blue curves in a). The error bars are the ranges of the intersections between the black curve in a) (data without extrusion) and the simultaneous, observational bounds of the fitted power laws for the large $s_{mi}$ regimes of the blue curves at 95% confidence levels. The dashed black line is a power law $\sim s_{ci}^{1/2}$.

Figure S4a shows $< r_{ij}^2(s_{ij}) >$, the mean square distance between locus $i$ and the cohesin binding site at locus $j$ as a function of the genomic separation between them $s_{ij}$ (see Fig. S1a). Each blue curve represents a different loop length $l$. Each blue curve initially follows the relaxed, unperturbed mean square size of $s_{ij}$ (the black curve), then crosses over to a weaker dependence on $s_{ij}$. The crossover occurs at longer genomic lengths for longer loop lengths. The crossover is predicted to occur at genomic length on the order of $g(l)$ (Eq. (S7)) and mean square size on the order of $\xi^2(l)$ (Eq. (S8)). These are the genomic length and mean square size, respectively, of the largest relaxed chromatin section within an extruded loop of length $l$ (see Fig. 5C in the main text).

We determine $g(l)$ and $\xi^2(l)$ from simulation in a similar manner to how we obtained $g_i$ and $\xi_i^2$ described above: we compare $MSD_{passive}$, the mean square displacement of beads in a simulated polymer *without* extrusion, to internal sizes $< r_{passive}^2(s) >$ *without* extrusion. $g(l)$ is the largest section with $MSD_{passive}(l/2v_{ex}) \geq < r^2_{passive}(s) >$, where $l/2v_{ex}$ is how long it takes to extrude a loop with length $l$. The mean square size $\xi^2(l)$ is $< r^2_{passive}(g(l)) >$, the relaxed, unextruded sizes of sections with genomic length $g(l)$. $g(l)$ and $\xi^2(l)$ were fitted to power laws such that $g(l) \approx (2.0 \pm 0.1) l^{(0.49 \pm 0.01)}$ and $\xi^2(l) \approx (3.04 \pm 0.04) l^{(0.53 \pm 0.01)}$, consistent with the predicted scaling behavior of $\sim l^{1/2}$ (see Eqs. (S7) and (S8)). Errors represent 95% confidence intervals.

We employ the same methods described for $< r^2_{mi}(s_{mi}) >$ to collapse the data for $< r^2_{ij}(s_{ij}) >$ in Figure S4a onto a universal curve. Figure S4b shows the data in Figure S4a collapsed by scaling the abscissa and ordinate by $g(l)$ and $\xi^2(l)$, respectively. The power laws fitted to the two regimes of the collapsed data were $< r^2_{ij}(s_{ij}) > / \xi^2(l) \approx (0.94 \pm 0.01) * [s_{ij}/g(l)]^{(1.01 \pm 0.05)}$ for $0.5 \leq s_{ij}/g(l) \leq 0.9$ and $< r^2_{ij}(s_{ij}) > / \xi^2(l) \approx (1.03 \pm 0.01) * [s_{ij}/g(l)]^{(0.57 \pm 0.01)}$ for $2 \leq s_{ij}/g_i \leq 7$. The intersection of the two curves is at $s_{ij}/g_i \approx 1.2 \pm 0.1$ and $< r^2_{ij}(s_{ij}) > / \xi^2(l) \approx 1.2 \pm 0.1$. The mean square distance between locus $i$ and the cohesin binding site $j$ in our simulations is consistent with a crossover between scaling regimes $\sim s^1_{ij}$ and $\sim s^{1/2}_{ij}$ at genomic length on the order of $1.2g(l)$ and mean square size on the order of $1.2\xi^2(l)$, in agreement with Equation (S10). Although the fitted exponent for the second regime with $2 \leq s_{ij}/g_i \leq 7$ is larger than the predicted value of $1/2$, this could once again be attributed to a wide crossover between two scaling regimes that does not reach the asymptotic behavior. Figure S4c shows the crossover genomic lengths $1.2g(l)$ (solid red curve) and mean square sizes $1.2\xi^2(l)$ (solid blue curve), where $g(l)$ and $\xi^2(l)$ are obtained from simulation. The circles in Figure S4c show the crossover points found by individually fitting the large $s_{ij}$ regimes of individual blue curves in Figure S4a. These individually fitted crossover points are consistent with $1.2g(l)$ and $1.2\xi^2(l)$.

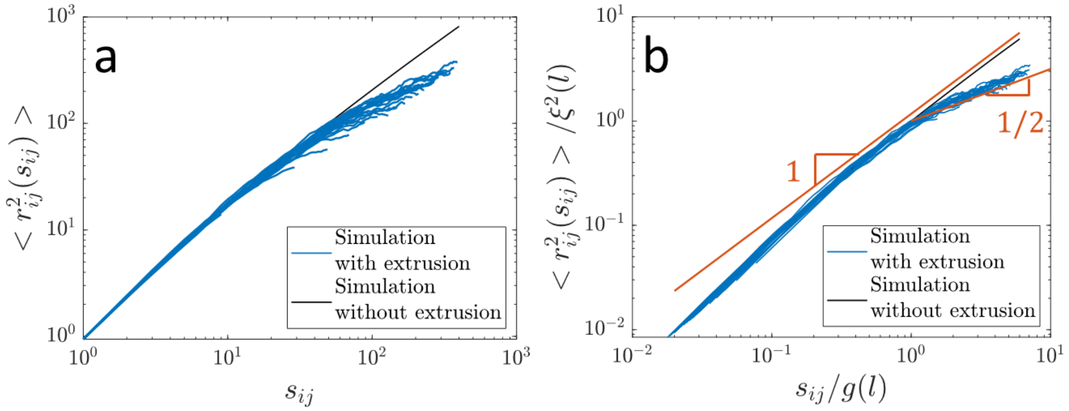

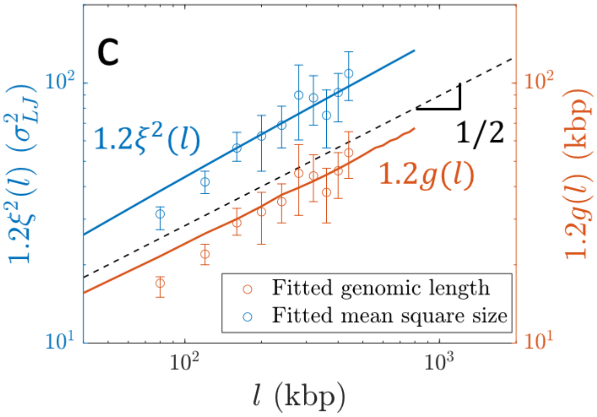


**Figure S4**: Mean square distance $< r_{ij}^2(s_{ij}) >$ between locus $i$ and the cohesin binding site $j$ from hybrid MD—MC simulations (related to Fig. S1e). a) Comparison of $< r_{ij}^2(s_{ij}) >$ as a function of genomic separation $s_{ij}$, with the relaxed, unperturbed mean square size of a section with genomic length $s_{ij}$ without extrusion. Each blue curve represents a different loop length $l$ =40, 80, 120, …, 800 kbp. b) The same data as a) with the abscissa and ordinate scaled by $g(l)$ and $\xi^2(l)$, respectively, determined from simulation. The red lines are power laws $\sim [s_{ij}/g(l)]^1$ and $\sim [s_{ij}/g(l)]^{1/2}$. C) Crossover locations of $< r_{ij}^2(s_{ij}) >$ as a function of loop length $l$. The solid curves are the crossovers found by collapsing all data by normalizing by $g(l)$ and $\xi^2(l)$. The circles are the crossovers determined by individually fitting the blue curves in a). The error bars are the ranges of the intersections between the black curve in a) (data without extrusion) and the simultaneous, observational bounds of the fitted power laws for the large $s_{ij}$ regimes of the blue curves at 95% confidence levels. The dashed black line is a power law $\sim l^{1/2}$.

Figure S5a shows the mean square distance between two loci extruded by the same cohesin domain separated by genomic distance $s$ within a loop of length $l$, averaged over all such loci pairs within the loop. Each blue curve corresponds to a different loop length. In Figure S5b, we collapse the data in Figure S5a onto a universal curve by scaling the abscissa and ordinate by $g(l)$ and $\xi^2(l)$, respectively, determined from simulation. $g(l)$ and $\xi^2(l)$ are the same as used in Figure S4b. The power laws fitted to the two regimes of the collapsed data were $< r_l^2(s) > /\xi^2(l) \approx (0.94 \pm 0.01) * [s/g(l)]^{(1.05\pm0.01)}$ for $0.5 \leq s/g(l) \leq 0.9$ and $< r_l^2(s) > /\xi^2(l) \approx (0.99 \pm 0.01) * [s/g(l)]^{(0.57\pm0.01)}$ for $2 \leq s/g(l) \leq 6$. The intersection of the two curves is at $s/g(l) \approx 1.11 \pm 0.05$ and $< r_l^2(s) > /\xi^2(l) \approx 1.12 \pm 0.08$. The mean square distance between two loci extruded by the same cohesin domain separated by $s$ in our simulations is consistent with a crossover between scaling regimes $\sim s^1$ and $\sim s^{1/2}$ at genomic length on the order of $1.1g(l)$ and mean square size on the order of $1.1\xi^2(l)$, in agreement with Equation (8) in the main text. Similar to the effect described in the preceding paragraphs, the higher exponent in the fitted power law for the regime with $2 \leq s/g(l) \leq 6$ compared to the theoretical value of $1/2$ could be a result of the narrow range of $s/g(l)$ available for fitting and a weak crossover between scaling regimes. Figure S5c shows the crossover genomic lengths $1.1g(l)$ (solid red curve) and mean square sizes $1.1\xi^2(l)$ (solid blue curve). The circles in Figure S5c show the crossover points found by individually fitting the large $s$ regimes of individual blue curves in Figure S5a, consistent with $1.1g(l)$ and $1.1\xi^2(l)$.

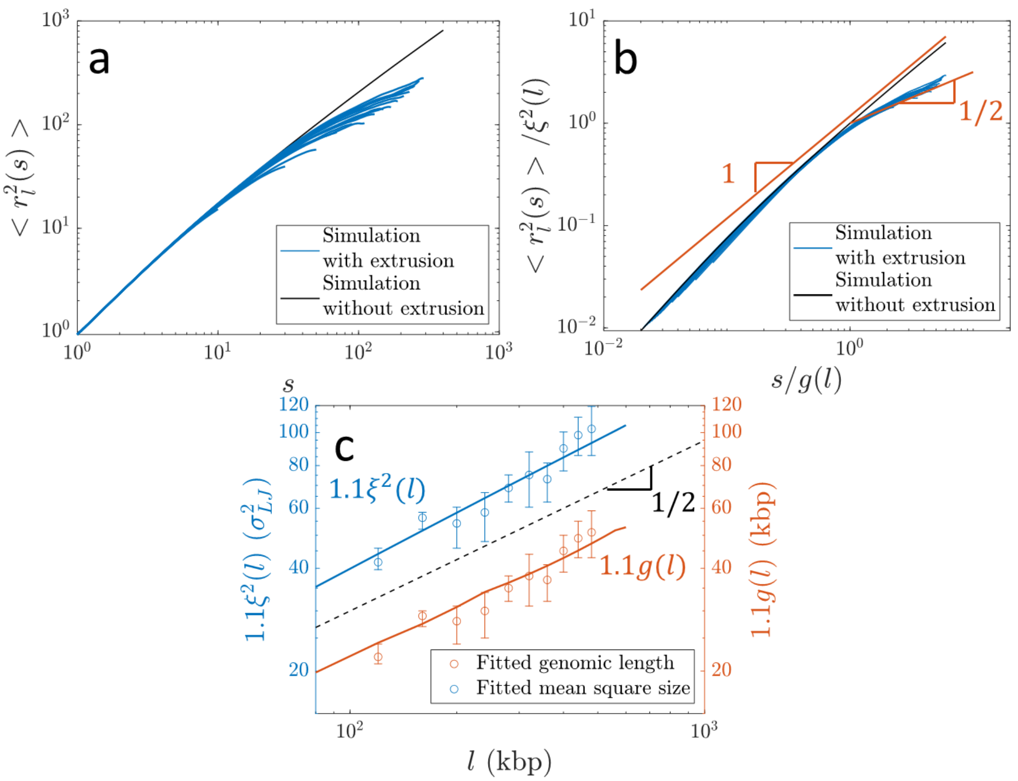


**Figure S5:** Mean square distance $< r_l^2(s) >$ between two loci extruded by the same cohesin domain separated by genomic distance $s$ within a loop of length $l$ from hybrid MD—MC simulations (related to Figs. 5D and 6A in the main text). a) Comparison of $< r_l^2(s) >$ with the relaxed, unperturbed mean square size of a section with genomic length $s$ without extrusion. Each blue curve represents a different loop length $l$ =40, 80, 120, …, 600 kbp. b) The same data as a) with the abscissa and ordinate scaled by $g(l)$ and $\xi^2(l)$, respectively, determined from simulation. The red lines are power laws ~ $[s/g(l)]^1$ and ~ $[s/g(l)]^{1/2}$. c) Crossover locations of $< r_l^2(s) >$ as a function of loop length $l$. The solid curves are the crossovers found by collapsing all data by normalizing by $g(l)$ and $\xi^2(l)$. The circles are the crossovers determined by individually fitting the blue curves in a). The error bars are the ranges of the intersections between the black curve in a) (data without extrusion) and the simultaneous, observational bounds of the fitted power laws for the large $s$ regimes of the blue curves at 95% confidence levels. The dashed black line is a power law ~ $l^{1/2}$.

**b. Steady-state extrusion regimes**

In steady-state extrusion, many cohesins bind to and unbind from a chromatin section. We base our steady-state model on our previous work in ref. (19), outlined here. We consider passive cohesin binding for definiteness, though we expect the results of our model to be consistent with other mechanisms of cohesin binding as well.

The separation $d$ is the average genomic length between two cohesins, and the processivity $\lambda$ is the average loop length extruded by an unobstructed cohesin. We define $\mu\, k_B T$ as the chemical potential per locus associated with cohesin binding. The average cohesin separation is $d \approx ze^{\mu}$. Bound cohesins have an energetic barrier of $h\, k_B T$ for unbinding during a time interval with duration $\tau_c$. During an interval with duration $\tau_c$, each locus is bound by a cohesin with probability $e^{-(\mu+h)}$, and each bound cohesin unbinds with probability $e^{-h}$. Both the residence time and processivity are exponentially distributed with averages $\tau_{res} \approx \tau_c e^h$ and $\lambda \approx 2v_{ex}\tau_{res} \approx 2v_{ex}\tau_c e^h$ respectively. The average time between cohesins binding the same locus is approximately $\tau_B \approx \tau_c e^{\mu+h}$. The average time between cohesins binding the same genomic section with length $s$ is approximately $\tau_{B,s} \approx \tau_B z/s \approx z\tau_c e^{\mu+h}/s$.

The average time between two cohesins binding within a chromatin section of length $\lambda$ is $\tau_{B,\lambda} \approx d/(2v_{ex})$, the time it takes to extrude the separation $d$. As shown in Figure 7 of the main text and related discussion, four regimes of processivity and separation dictate chromatin compaction and loop nesting on genomic length scales shorter than the genomic entanglement length $N_e$. In Regime I with $\lambda \leq 2g_{min} \approx 2z/\kappa$, loops can fully relax during the time it takes to extrude them. Recall that $g_{min}$ is the smallest relaxed section next to a cohesin in an extruded loop. Loops do not significantly impact average contact probabilities.

In Regime II, loops can relax in the time between two cohesins binding the same genomic section with length $\lambda$. This means $\tau_{B,\lambda} > \tau_{res} + \tau_0(\lambda/z)^2$, where the right-hand side is the average cohesin residence time plus the relaxation time of $\lambda$. Rewriting in terms of $\lambda$ and $d$ yields $d > \lambda(1 + 2\kappa\lambda/z)$. Each loop is compact while a cohesin extrudes them but have enough time to relax such that the average conformation of the section is relaxed. In Regime III, loops *do not* have time to relax between two cohesin binding events. The chromatin section is compact, but loops are still unnested. Regime III with compact but unnested chromatin corresponds to separation $d$ between two boundaries $\lambda < d < \lambda(1 + 2\kappa\lambda/z)$. In Regime IV, loops are nested with $\lambda \geq d$ and the chromatin section is compact.

**c. Entanglement suppression**

We define the overlap parameter as the number of chromatin sections with the same genomic length $s$ occupying the same approximately spherical volume with diameter $\approx < r^2(s) >^{1/2}$, the root-mean-square size of $s$:

$$O(s) = \frac{\phi}{\phi_{self}(s)} \approx \frac{\phi \frac{4}{3}\pi \left(\frac{< r^2(s) >^{\frac{1}{2}}}{2}\right)^3}{v\left(\frac{s}{z}\right)}. \tag{S12}$$

The numerator is the volume of chromatin within the sphere assuming an average chromatin volume fraction in the nucleus $\phi$. The denominator is the physical volume of chromatin in $s/z$ loci, where $v$ is the volume of a locus (see Eq. (S1)). Eq. (S12) simplifies to

$$O(s) \approx \phi \frac{\pi z}{6sv} < r^2(s) >^{\frac{3}{2}}. \tag{S13}$$

Entanglements occur when the overlap parameter is on the order of $O_{KN} \approx 10 - 20$ (23). If chromatin has a fractal dimension of $D$ on length scales between $b$ and $< r^2(s) >^{1/2} \approx b(s/z)^{1/D}$, the overlap parameter is

$$O(s) \approx \phi \frac{\pi z}{6sv} b^3 \left(\frac{s}{z}\right)^{\frac{3}{D}} \approx \phi \frac{\pi b^3}{6v} \left(\frac{s}{z}\right)^{\frac{3}{D}-1}. \tag{S14}$$

In the passive case without loop extrusion, chromatin has fractal dimension $D \approx 2$ on scales below entanglements, so Eq. (S14) becomes

$$O_{passive}(s) \approx \phi \frac{\pi b^3}{6v} \left(\frac{s}{z}\right)^{\frac{1}{2}}. \tag{S15}$$

The passive entanglement genomic length is the genomic length $s$ that makes Eq. (S15) equal to $O_{KN}$:

$$N_{e,passive} \approx z\left(\frac{O_{KN}6v}{\phi\pi b^3}\right)^2 \qquad \text{(S16)}$$

which for $O_{KN} = 10$, $\phi = 0.15$, $b = 50$ nm, and $v \approx 7.5\text{x}10^3$ nm$^3$ yields $N_{e,passive} \approx 100$ kbp.

For steady-state active extrusion in interphase, the transition in fractal dimension from $D \approx 2$ to $D \approx 4$ and back to $D \approx 2$ changes $< r^2(s) >$ in Eq. (S13) such that

$$< r^2(s) > \approx \begin{cases} b^2\frac{s}{z}\ , & s \le g(d) \\ b^2\frac{[g(d)s]^{\frac{1}{2}}}{z}\ , & g(d) < s \le C_2\overline{N_{TAD}}\ , \\ b^2\left[\frac{g(d)}{C_2\overline{N_{TAD}}}\right]^{\frac{1}{2}}\frac{s}{z}\ , & C_2\overline{N_{TAD}} < s \le N_e \end{cases} \qquad \text{(S17)}$$

where $g(d)$ is the genomic length of the longest relaxed chromatin section in steady-state extrusion (see Eq. (10) in the main text), $\overline{N_{TAD}}$ is the average TAD length of the genomic section of interest, and $C_2$ is a constant on the order of unity. The overlap parameter is approximately

$$O_{active}(s) \approx \begin{cases} \phi\frac{\pi b^3}{6v}\left(\frac{s}{z}\right)^{\frac{1}{2}}\ , & s \le g(d) \\ \phi\frac{\pi b^3}{6v}\left[\frac{g(d)^3}{sz^2}\right]^{\frac{1}{4}}\ , & g(d) < s \le C_2\overline{N_{TAD}}\ , \\ \phi\frac{\pi b^3}{6v}\left[\frac{g(d)}{C_2\overline{N_{TAD}}}\right]^{\frac{3}{4}}\left(\frac{s}{z}\right)^{\frac{1}{2}}\ , & C_2\overline{N_{TAD}} < s \le N_e \end{cases} \qquad \text{(S18)}$$

see Figure 9A in the main text. The overlap parameter does not reach $O_{KN}$ because the longest relaxed chromatin section length $g(d)$ is shorter than the entanglement length $N_{e,passive} \approx 100$ kbp of chromatin in the passive state without extrusion. In the second regime with fractal dimension four, the overlap parameter $O_{active}(s)$ decreases toward unity. $O_{active}(s)$ starts increasing in the third regime with $s > C_2\overline{N_{TAD}}$.

The active entanglement genomic length is determined by the condition of the overlap parameter $O_{active}(s)$ in this third regime (the last line of Eq. (S18)) reaching $O_{KN}$:

$$N_{e,active} \approx z\left(\frac{O_{KN}6v}{\phi\pi b^3}\right)^2\left[\frac{C_2\overline{N_{TAD}}}{g(d)}\right]^{\frac{3}{2}} \approx N_{e,passive}\left[\frac{C_2\overline{N_{TAD}}}{g(d)}\right]^{\frac{3}{2}}, \qquad \text{(S19)}$$

which for $N_{e,passive} \approx$ 100 kbp, $C_2\overline{N_{TAD}} \approx 400$ kbp, and $g(d) \approx 30$ kbp gives $N_{e,active} \approx 5$ Mbp. Overlap suppression $O_{passive}(s)/O_{active}(s)$ is maximized at $s \approx C_2\overline{N_{TAD}}$, which is the genomic length scale at which chromatin fractal dimension crosses over from $D \approx 4$ back to $D \approx 2$ with active loop extrusion. The maximum overlap suppression is

$$\frac{O_{passive}\left(C_2\overline{N_{TAD}}\right)}{O_{active}\left(C_2\overline{N_{TAD}}\right)} \approx \left[\frac{C_2\overline{N_{TAD}}}{g(d)}\right]^{\frac{3}{4}}, \tag{S20}$$

which for $C_2\overline{N_{TAD}} \approx 400$ kbp and $g(d) \approx 30$ kbp is approximately equal to 7.

**d. "Legs" and cohesin trajectories**

As described in the main text, the conformation of extended sections of chromatin adjacent to cohesins outside of extruded loops ("legs") are determined by cohesin translocation, not polymer relaxation. Legs are extended such that

$$g_{leg}(t) \sim < r_{leg}^2(t) >^{1/2} \sim (v_{ex}t)^{1/D} , \tag{S21}$$

where $< r_{leg}^2(t) >^{1/2}$ is the root-mean-square size of a leg and $t$ is time since the beginning of extrusion. In Figure S6a, we plot the genomic length of legs $g_{leg}(t)$ as a function of the length of chromatin extruded by the cohesin $v_{ex}t$ for $v_{ex} \approx 0.15$ kbp/$\tau_0$, 0.3 kbp/$\tau_0$, and 0.6 kbp/$\tau_0$ from simulations of a single extrusion cycle starting from $D \approx 2$. We fit the data to a power law

$$g_{leg}(t) \approx A * (v_{ex}t)^B \tag{S22}$$

and find $A \approx 2.1 \pm 0.5$ and $B \approx 0.54 \pm 0.05$, consistent with the prediction $g_{leg}(t) \sim (v_{ex}t)^{1/2}$ (Eq. (S21)).

Cohesin trajectories follow the midpoints of the tension fronts produced by extrusion (see Fig. 11A in the main text). The location of the tension fronts at time *t* can be approximated from the initial chromatin conformation on which cohesin binds. As such, the initial chromatin conformation can be used to approximate the cohesin trajectory (see Fig. 11B in the main text and section IIIa in the SI). The fluctuations in the cohesin trajectory compared to tension front midpoints are on the order of $\approx b^2(t/\tau_0)^{1/2}$, which is the mean square distance a locus moves in time *t*. The mean square distance between cohesin trajectories and tension front midpoints from simulations $< \Delta_{coh,tens}^2(t) >$ is consistent with this scaling (see Fig. S6b).

Consider two un-nested, neighboring cohesins actively extruding the same genomic section (see Fig. S6c). Initially, the two cohesins travel independently in space before their tension fronts meet in the intervening genomic section. As described in the main text, once the tension fronts meet, the cohesins continue pulling and move towards each other. After the cohesins meet, they travel together in space; this effect is more pronounced if the cohesins cannot traverse one another (see Fig. S6d), where the two cohesins will follow the midpoint between the outermost tension fronts.

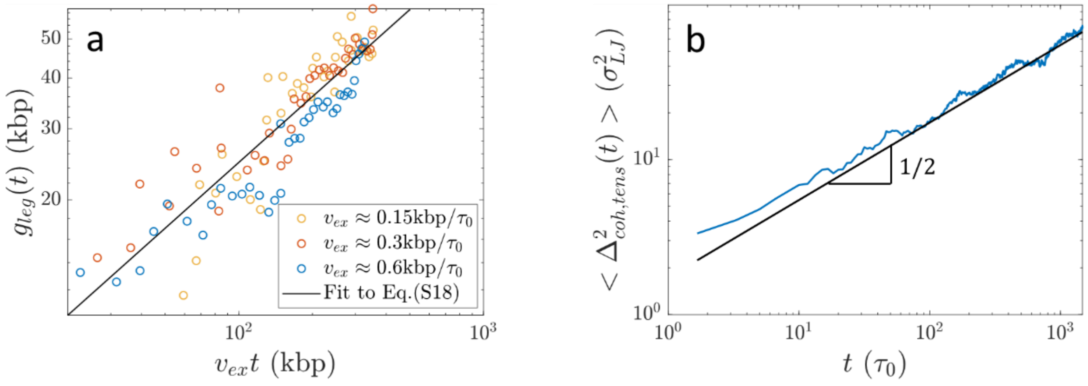

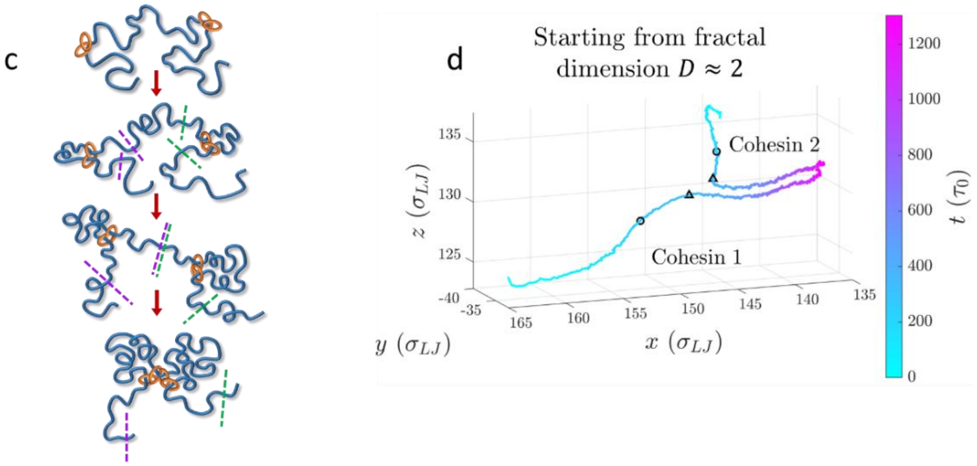


**Figure S6:** Extended legs and cohesin trajectories formed by active loop extrusion. a) Genomic length of extended legs at $t$ after extrusion starts from simulations of a single cohesin extruding an initially relaxed chromatin section with $D \approx 2$ using three different extrusion velocities. The black line is a power law fit (Eq. (S22)) to all points such that $g_{leg}(t) \approx A * (v_{ex}t)^B$ with $A = 2.1 \pm\ 0.5$ and $B = 0.54 \pm\ 0.05$. See section IIIa for leg length measurement method. b) Mean square distance between cohesin trajectories and tension front midpoints at time $t$ after extrusion starts from simulations of a single cohesin extruding an initially relaxed chromatin section with $D \approx 2$ and $v_{ex} \approx 0.3$ kbp/$\tau_0$. The blue curve is the data, and the black curve is $(3t)^{1/2}$. c) Schematic of two un-nested, neighboring cohesins extruding the same genomic section. Purple and green dashed lines indicate tension fronts produced by the two cohesins. d) Average cohesin trajectories of two un-nested cohesins extruding a single polymer chain from 200 simulations with the same initial conformations with $D \approx 2$ in which the cohesins cannot traverse one another. Black circles indicate the approximate time and locations when tension fronts meet; black triangles indicate approximate time and locations when cohesins meet.

##### e. Mean square displacement of cohesins and chromatin loci

<u>Cohesin MSD</u>

Cohesin dynamics are tied to the genomic section on which it is extruding. During short lag times, cohesin motion couples to the relaxation modes of a section with genomic length $g_{min}$. Its mean square displacement $MSD_{coh}$ is the same as if the cohesin were not actively extruding and instead acted like a crosslink. For these short lag times, the cohesin MSD is on the order of

$$MSD_{coh}(\Delta t) \approx D_{coh} b^2 \left(\frac{t}{\tau_0}\right)^{\frac{1}{2}}. \quad \text{(S23)}$$

$D_{coh}$ accounts for effective friction experienced by cohesin (for example, due to size or temporary associations with other proteins), and is approximately the ratio $MSD_{coh}(\tau_0)/MSD_{chr}(\tau_0)$, where $MSD_{chr}(\tau_0)$ is the mean square displacement of a chromatin locus during lag time $\tau_0$. This regime ends at the relaxation time of $g_{min}$ corresponding to lag time

$$\Delta t_{coh}^{+} \approx \tau_0 \kappa^{-2}\ . \quad \text{(S24)}$$

During longer lag times, cohesin trajectories follow the tension fronts that are determined by the chromatin conformation at the time of cohesin binding (see previous section and main text). In Figure S6a, we sketch examples of the predicted cohesin MSD for different initial chromatin conformations:

i) Relaxed with $D \approx 2$, and
ii) Confined to a radius of gyration $R_g$.

In general, we expect $MSD_{coh} \sim \Delta t^{2/D}$ for $\Delta t \geq \Delta t^{+}_{coh}$. For case i), cohesins perform effective diffusion with $MSD_{coh} \sim \Delta t^{1}$. For case ii), cohesin MSD is diffusive before plateauing when $MSD_{coh} \approx R_g^2$. In Figure S6b, we plot cohesin MSDs from hybrid MD—MC simulations of single extrusion cycles corresponding to case i) (purple curve) and case ii) (green curve) with $\kappa \approx 0.3$ (see "Extended methods", section III below). We compare them to the MSD of a cohesin that acts as a crosslink and does not extrude (black curve). As expected, $MSD_{coh}$ for both extrusion cases are approximately equal to the crosslink MSD during short lag times. For our simulations, $MSD_{coh}(\tau_0)/MSD_{chr}(\tau_0) \approx 0.5 \approx D_{coh}$ because the simulated cohesin bond effectively acts as the junction point of a star polymer when not extruding. After lag times $\Delta t^{+}_{coh} \approx 12\tau_0$ shown with the dashed black line in Figure S6b, both the purple and green curves are consistent with $MSD_{coh} \sim \Delta t^{1}$. $\Delta t^{+}_{coh} \approx 12\tau_0$ is the relaxation time of $g_{min}$ determined from simulation (see "Extended methods"). Lastly, the green curve plateaus when the cohesin MSD approximately equals the twice the mean squared radius of gyration of confinement (dashed green line), which is consistent with the scaling prediction of a plateau on the order of $MSD_{coh} \approx R_g^2$.

For the case of steady-state extrusion, chromatin conformation crosses over from fractal dimensions $D \approx 2$ to $D \approx 4$ at genomic lengths on the order of $g(d) \approx [dz^2/(2v_{ex}\tau_0)]^{1/2} \approx [dz/(2\kappa)]^{1/2}$ with mean square size $\xi^2(d) \approx b^2[d/(2\kappa z)]^{1/2}$ (see Eq. (10) in the main text and related discussion). So, the mean square displacement of cohesin $MSD_{coh}$ crosses over between scaling behaviors of $MSD_{coh} \sim \Delta t^{1}$ and $MSD_{coh} \sim \Delta t^{1/2}$ at lag time

$$\Delta t^{++}_{coh} \approx D_{coh}^{-1}\left(\frac{dz}{2\kappa v_{ex}^2}\right)^{\frac{1}{2}} \approx D_{coh}^{-1} g(d)/v_{ex}\,, \tag{S25}$$

(see Fig. S6c). This lag time is how long it takes $MSD_{coh}$ to reach $\xi^2(d)$. It is also approximately the time it takes one cohesin domain to extrude a genomic length of $D_{coh}^{-1}g(d)$. $MSD_{coh}$ for lag times between $\Delta t^{+}_{coh}$ and $\Delta t^{++}_{coh}$ is approximately

$$MSD_{coh}(\Delta t) \approx D_{coh} b^2 \kappa \frac{\Delta t}{\tau_0}\,, \text{ for } \Delta t^{+}_{coh} < \Delta t \leq \Delta t^{++}_{coh}\,. \tag{S26}$$

Note that $MSD_{coh}(\Delta t^{+}_{coh}) \approx D_{coh} b^2 \kappa^{-1} \approx D_{coh}\xi^2_{min}$, where $\xi^2_{min} \approx b^2\kappa^{-1}$ is the mean square size of the smallest relaxed section in a loop with genomic length $g_{min}$ (see Fig. 5B and related discussion in the main text). $MSD_{coh}$ for lag times longer than $\Delta t^{++}_{coh}$ is approximately

$$MSD_{coh}(\Delta t) \approx b^2 \left[\frac{g(d) D_{coh} \kappa \Delta t}{z\tau_0}\right]^{\frac{1}{2}}, \text{ for } \Delta t > \Delta t^{++}_{coh}\,. \tag{S27}$$

The crossover times $\Delta t^{+}_{coh}$ and $\Delta t^{++}_{coh}$ both depend on $\tau_0$, which in turn depends on the $D_{app}$ of the chromatin section of interest (see Eq. (S2)). Note that $\Delta t^{+}_{coh}$ and $\Delta t^{++}_{coh}$ are meaningless for $\kappa \leq 2z/\lambda$. This condition corresponds to Regime I in Figure 7A in the main text, in which entire loops have time to fully relax in the time cohesin extrudes them. In this case, there is no regime with $MSD_{coh} \sim \Delta t^{1}$.

Figure S6d shows the cohesin MSD from a steady-state extrusion simulation with $\kappa \approx 0.15$ (solid blue curve) compared to the MSD of a cohesin that acts as a crosslink (solid black curve). We expect the cohesin MSD with extrusion to crossover between scaling behaviors $MSD_{coh} \sim \Delta t^{1/2}$ (see Eq. (S23)) and $MSD_{coh} \sim \Delta t^{1}$ (see Eq. (S26)) at a lag time of approximately $\Delta t^{+}_{coh} \approx$

$50\tau_0$, the relaxation time of $g_{min}$ obtained from simulation (see "Extended methods"). We then expect a second crossover back to $MSD_{coh} \sim \Delta t^{1/2}$ (see Eq. (S27)) at a lag time of approximately $\Delta t_{coh}^{++} \approx 250\tau_0$. This value of $\Delta t_{coh}^{++}$ was calculated from Eq. (S25), using $D_{coh} \approx 0.5$ and $g(d) \approx 37$ kbp, as determined from simulation *without* extrusion (see "Extended methods"). In Figure S6d we also plot a theoretical crossover function (solid green curve) following this expected scaling behavior:

$$MSD_{coh}(\Delta t) \approx \Delta t^{\frac{1}{2}} \left[1 + \left(\frac{\Delta t}{50\tau_0}\right)^{\frac{1}{3}}\right]^{\frac{3}{2}} \left[1 + \left(\frac{\Delta t}{250\tau_0}\right)^{2}\right]^{-\frac{1}{4}}. \tag{S28}$$

The cohesin MSD from steady-state extrusion simulation (solid blue curve) is qualitatively consistent with Eq. (S28) (solid green curve).

To further characterize the data, we fit the cohesin MSD from the steady-state extrusion simulation to two power laws for lag times longer than $\Delta t_{coh}^{+}$. The fits yielded $MSD_{coh}(\Delta t) \approx (0.73 \pm 0.01) * (\Delta t/\tau_0)^{(0.802\pm0.003)}$ for $50 \leq \Delta t/\tau_0 \leq 150$ and $MSD_{coh}(\Delta t) \approx (5.25 \pm 0.06) * (\Delta t/\tau_0)^{(0.442\pm0.002)}$ for $300 \leq \Delta t/\tau_0 \leq 1000$. We also fit the crosslink MSD (black curve) to $MSD_{coh}(\Delta t) \approx (1.40 \pm 0.01) * (\Delta t/\tau_0)^{(0.570\pm0.001)}$ for $20 \leq \Delta t/\tau_0 \leq 300$. The errors are 95% confidence intervals. We find the intersections between the fitted power laws to fit the crossover locations between scaling regimes, indicated by dashed black and blue lines in Figure S6d. Note that the three fitted power laws do not exactly match the theoretically predicted behaviors of $\sim \Delta t^1$ for $50 \leq \Delta t/\tau_0 \leq 150$ and $\sim \Delta t^{1/2}$ for the interval $300 \leq \Delta \mathrm{t}/\tau_0 \leq 1000$ and the crosslink MSD for $20 \leq \Delta t/\tau_0 \leq 300$. We attribute this discrepancy to a broad crossover between narrow intervals of scaling regimes. Note that the simulation data (solid blue curve in Fig. S6d) is consistent with the theoretical crossover function (solid green curve, Eq. (S28)) that was constructed with the theoretical power laws.

The first crossover was found at lag times of $\approx (0.3 \pm 0.1)\Delta t_{coh}^{+}$ corresponding to mean square displacements of $\approx (0.26 \pm 0.07)\xi_{min}^2$, where $\Delta t_{coh}^{+} \approx 50\tau_0$ and $\xi_{min}^2 \approx 27\sigma_{LJ}^2$ were determined from simulation without extrusion (see "Extended methods"). The errors represent the range of the intersection between the simultaneous, observational bounds of the fitted power laws at 95% confidence intervals. The second crossover was found at lag times of $\approx (0.9 \pm 0.2)\Delta t_{coh}^{++}$ corresponding to mean square displacements of $\approx (0.8 \pm 0.1)\xi^2(d)$, where $\Delta t_{coh}^{++} \approx 250\tau_0$ was calculated using Eq. (S25) (see previous paragraph) and $\xi^2(d) \approx 73\sigma_{LJ}^2$ was determined from simulation (see "Extended methods"). This fitted crossover agrees with the theoretical prediction.

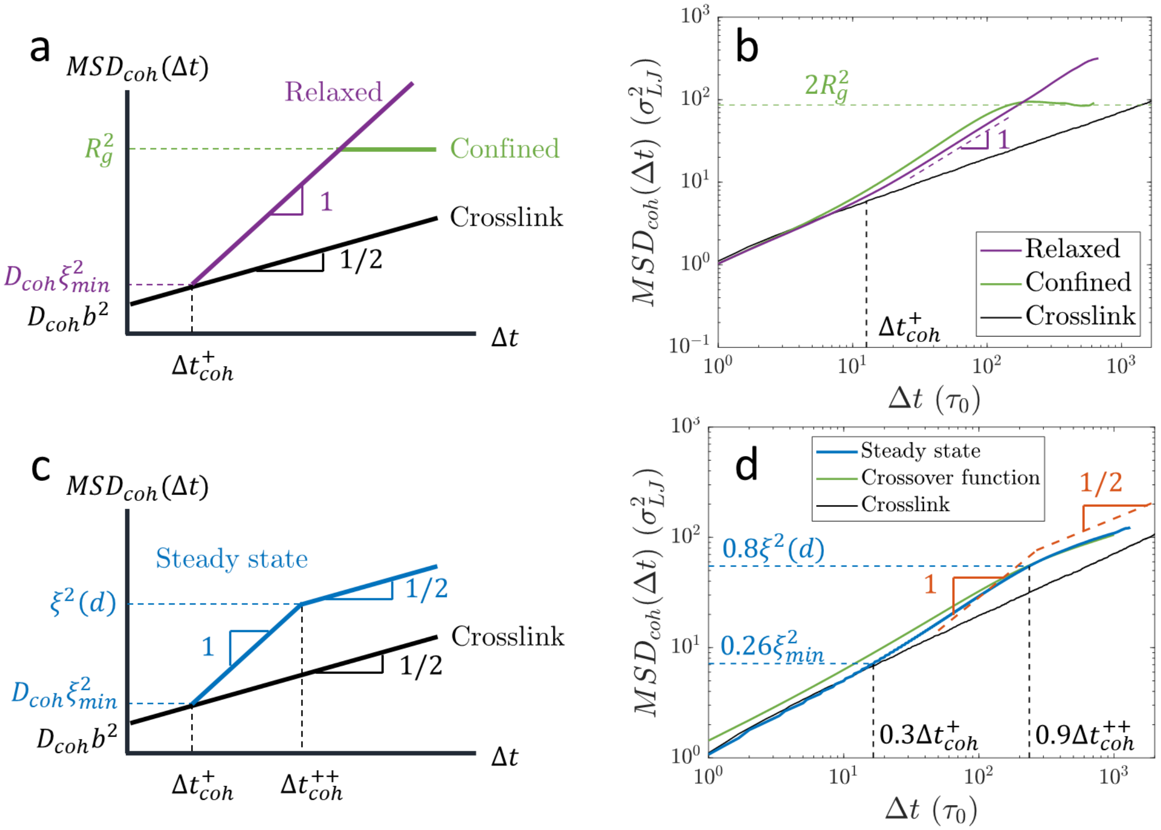


**Figure S6:** MSD of cohesins and chromatin loci during active extrusion, neglecting entanglement effects at long lag times. a) Schematic plot on a log-log scale of the predicted MSD of actively extruding, chromatin-bound cohesins during a single extrusion cycle on an initially relaxed chromatin section with $D \approx 2$ (purple) or a chromatin section initially confined to a radius of gyration $R_g$ (green). The black line shows the MSD of a cohesin that does not extrude, but rather acts as a crosslink on a relaxed chromatin section with $D = 2$. b) Cohesin MSD from simulations of a single extrusion cycle corresponding to a) with $v_{ex} \approx 0.6$ kbp/$\tau_0$ ($\kappa \approx 0.3$). The dashed green line is $2R_g^2$, where $R_g^2$ is the mean square radius of gyration of confinement. The dashed vertical line corresponding to $\Delta t_{coh}^+$ was obtained from simulation as the relaxation time of $g_{min}$ (see "Extended methods"). c) Schematic plot on a log-log scale of unobstructed cohesin predicted MSD in steady-state extrusion without TADs. d) Cohesin MSD from a steady-state active extrusion simulation with $v_{ex} \approx 0.3$ kbp/$\tau_0$ (solid blue curve) compared with the MSD of a cohesin that acts as a crosslink which does not extrude (solid black curve) and a theoretical crossover function (solid green curve, see Eq. (S28)). The dashed red lines show the theoretical scaling predictions $\sim \Delta t^1$ and $\sim \Delta t^{1/2}$. The black dashed vertical lines are the lag times corresponding to the intersections of power laws fitted to the cohesin MSD and crosslink MSD. The blue dashed horizontal lines are the corresponding cohesin MSDs. The first crossover point is at lag times of $\approx (0.3 \pm 0.1)\Delta t_{coh}^+$ and $MSD_{coh}$ of $\approx (0.26 \pm 0.07)\xi_{min}^2$. The second crossover point is at lag times of $\approx (0.9 \pm 0.2)\Delta t_{coh}^{++}$ and $MSD_{coh}$ of $\approx (0.8 \pm 0.1)\xi^2(d)$.

#### Chromatin loci MSD

Monomers in a polymer with fractal dimension $D$ are predicted to follow Rouse-like MSDs that scale as $\sim \Delta t^{2/(D+2)}$ (24). With steady-state active loop extrusion in interphase, the fractal dimension of chromatin crosses over between $D \approx 2$ and $D \approx 4$ and back to $D \approx 2$. The scaling behavior of chromatin MSD thus crosses over between $\sim \Delta t^{1/2}$ and $\sim \Delta t^{1/3}$ and back to $\sim \Delta t^{1/2}$ at lag times of $\Delta t_{chr}^+$ and $\Delta t_{chr}^{++}$ respectively (see Fig. S7a). On short length scales, chromatin has fractal dimension $D \approx 2$, so chromatin loci have MSD

$$MSD_{chr}(\Delta t) \approx b^2 \left(\frac{\Delta t}{\tau_0}\right)^{\frac{1}{2}} \approx D_{app}\Delta t^{\frac{1}{2}} \quad \text{for } \Delta t \leq \Delta t_{chr}^+ \,. \qquad \text{(S29)}$$

The fractal dimension crosses over to $D \approx 4$ on genomic length scales on the order of $g(d) \approx [dz/(2\kappa)]^{1/2}$. The lag time $\Delta t_{chr}^{+}$ at which chromatin locus MSD crosses over to a scaling of $\sim \Delta t^{1/3}$ is when Eq. (S29) is approximately equal to $\xi^2(d) \approx b^2(g(d)/z) \approx b^2[d/(2\kappa z)]^{1/2}$, which is the mean square spatial size of $g(d)$. This yields

$$\Delta t_{chr}^{+} \approx \frac{d}{2v_{ex}} \quad \text{(S30)}$$

corresponding to MSDs on the order of

$$MSD_{chr}(\Delta t_{chr}^{+}) \approx b^2 \left(\frac{d}{2\kappa z}\right)^{\frac{1}{2}} \approx D_{app} \left(\frac{d}{2v_{ex}}\right)^{\frac{1}{2}}. \quad \text{(S31)}$$

The magnitudes of $MSD_{chr}(\Delta t_{chr}^{+})$ and $MSD_{coh}(\Delta t_{coh}^{++})$ are approximately equal: $MSD_{coh}(\Delta t_{coh}^{++}) \approx MSD_{chr}(\Delta t_{chr}^{+}) \approx b^2[d/(2\kappa z)]^{1/2}$.

Using $d \approx 200$ kbp, $v_{ex} \approx 0.1$ kbp/s, and $D_{app} \approx 1.3\text{x}10^{-3}$ μm$^2$ s$^{-1/2}$ gives $\Delta t_{chr}^{+} \approx 15$ minutes and $MSD_{chr}(\Delta t_{chr}^{+}) \approx 0.04$ μm$^2$. The chromatin locus MSD is then approximately

$$MSD_{chr}(\Delta t) \approx D_{app} \left(\frac{d}{2v_{ex}}\right)^{\frac{1}{6}} \Delta t^{\frac{1}{3}} \text{ for } \Delta t_{chr}^{+} < \Delta t \leq \Delta t_{chr}^{++}. \quad \text{(S32)}$$

Note that the two-point MSD of the relative position of two loci would have approximately twice the magnitude of the single-point MSD that we describe here. That is, the two-point MSD at $\Delta t^*$ would be approximately 0.08 μm$^2$.

After $\Delta t_{chr}^{++}$, the MSD returns to a scaling of $MSD_{chr} \sim \Delta t^{1/2}$, reflecting the crossover in chromatin fractal dimension back to $D \approx 2$ on genomic length scales longer than $\approx C_2\overline{N_{TAD}}$ (or $\approx C_1\lambda$ without TAD anchors) with mean square size (with active loop extrusion)

$$< r^2\left(C_2\overline{N_{TAD}}\right) > \approx b^2 \left[\frac{g(d)}{z}\right]\left[\frac{C_2\overline{N_{TAD}}}{g(d)}\right]^{\frac{1}{2}} \approx b^2 \left[\frac{g(d)C_2\overline{N_{TAD}}}{z^2}\right]^{\frac{1}{2}}. \quad \text{(S33)}$$

The crossover time $\Delta t_{chr}^{++}$ is found by equating Eqs. (S32) and (S33). For even longer lag times $\Delta t \geq \Delta t_{e,active}$, chromatin entanglements cause the MSD to exhibit a scaling of $MSD_{chr} \sim \Delta t^{1/4}$ since $D \approx 3$ on genomic length scales longer than $N_{e,active}$ and mean square distances longer than $< r^2(N_{e,active}) >$ with extrusion. For simplicity, in Figure S7a we do not plot this last regime (for neither the active nor passive case).

Figure S7b shows the MSD of the central chromatin locus from a hybrid MD—MC simulation of steady-state extrusion without TAD anchors using $v_{ex} \approx 0.3$ kbp/$\tau_0$ ($\kappa \approx 0.15$). This MSD is found by tracking the position of the center of mass of the middle two beads of the simulated polymer chain. We fit the data to three power laws. The fitted functions were $MSD_{chr}(\Delta t) \approx (2.98 \pm 0.04) * (\Delta t/\tau_0)^{(0.56 \pm 0.01)}$ for $1.5\text{x}10^2\tau_0 \leq \Delta t \leq 3.5\text{x}10^2\tau_0$, $MSD_{chr}(\Delta t) \approx (11.8 \pm 0.2) * (\Delta t/\tau_0)^{(0.348 \pm 0.002)}$ for $1.3\text{x}10^3\tau_0 \leq \Delta t \leq 3\text{x}10^3\tau_0$, and $MSD_{chr}(\Delta t) \approx (2.0 \pm 0.1) * (\Delta t/\tau_0)^{(0.545 \pm 0.005)}$ for $4.4\text{x}10^3\tau_0 \leq \Delta t \leq 1.4\text{x}10^4\tau_0$. The errors represent 95% confidence intervals. While the fitted power laws are close to the theoretical scaling regimes crossing over between $MSD_{chr} \sim \Delta t^{1/2}$ to $MSD_{chr} \sim \Delta t^{1/3}$ and back to $MSD_{chr} \sim \Delta t^{1/2}$, all of the fitted exponents are larger in magnitude than predicted. In the first regime, this could be due to a broad crossover between ballistic MSD at short lag times to subdiffusive behavior. For intermediate lag times, the fitted exponent of $\approx 0.348 \pm 0.002$ is larger than $1/3$, possibly because there is a narrow range (on the order of a decade) of this regime between two other regimes with higher exponents. In the last regime, the discrepancy between the fitted exponent and theoretical value of $1/2$ could be because in our simulation, the MSD at the upper end of this narrow fitting interval $MSD_{chr}(1.4\text{x}10^4\tau_0) \approx 420\sigma_{LJ}^2$ approaches the overall mean square end-to-end distance of the polymer chain of $\approx 450\sigma_{LJ}^2$. As such, our simulated MSD has a crossover to diffusive behavior. This would not be present in biological chromatin, as instead the effect of entanglements would appear.

The crossovers between the three scaling regimes were determined by finding the intersection between the fitted power laws. The first crossover between the fitted power laws $\sim \Delta t^{0.56}$ and $\sim \Delta t^{0.348}$ occurs at a lag time of $\Delta t_{chr,fit}^{+} \approx (640 \pm 40)\tau_0$ corresponding to MSD of $\approx (1.6 \pm 0.1)\xi^2(d)$, where $\xi^2(d) \approx 73\sigma_{LJ}^2$ is obtained from simulation (see "Extended methods"). The error represents the range of the intersection between the simultaneous, observational bounds of the fitted power laws at 95% confidence levels. The second crossover between the fitted power laws $\sim \Delta t^{0.348}$ and $\sim \Delta t^{0.545}$ occurs at a lag time of $\Delta t_{chr,fit}^{++} \approx (4.6 \pm 0.9)\text{x}10^3\tau_0$ with MSD of $\approx (1.0 \pm 0.1) < r^2(2\lambda) >$, where $< r^2(2\lambda) >$ is the mean square size of a chromatin section with genomic length $2\lambda$ with active loop extrusion determined from simulation. As discussed in the main text (see Fig. 8B and related discussion), this simulation causes a change in fractal dimension from $D \approx 4$ to $D \approx 2$ at genomic length scales of $\approx 2\lambda$ (*i.e.*, the constant $C_1 \approx 2$) as evidenced by the change in contact probability scaling from $P(s) \sim s^{-3/4}$ to $P(s) \sim s^{-3/2}$. Recall that this is because extruding proteins rarely extrude loops longer than processivity $\lambda$. Our simulations did not exhibit the crossover to $MSD_{chr} \sim \Delta t^{1/4}$ at the longest time scales because our system is not entangled.

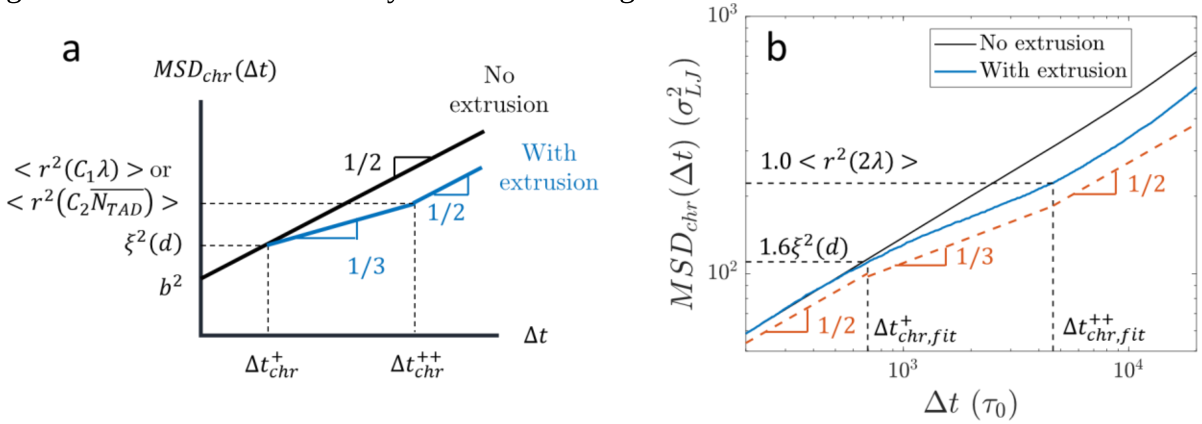


**Figure 7:** Mean square displacement $MSD_{chr}(\Delta t)$ of chromatin loci undergoing active loop extrusion in a steady state. a) Schematic plot on a log-log scale of $MSD_{chr}(\Delta t)$ with and without extrusion, without considering entanglements for simplicity. The MSD of the second crossover is $\approx < r^2(C_1\lambda) >$ without TAD anchors, or $\approx < r^2(C_2\overline{N_{TAD}}) >$ with TAD anchors. b) Chromatin locus MSD during steady-state active loop extrusion without TADs from simulation with $v_{ex} \approx 0.3$ kbp/$\tau_0$ ($\kappa \approx 0.15$). Dashed red lines show the theoretical scaling behaviors of $\sim \Delta t^{1/2}$ and $\sim \Delta t^{1/3}$. The dashed vertical lines are the lag times corresponding to the intersections of power laws fitted to the chromatin locus MSD. The dashed horizontal lines are the corresponding chromatin locus MSDs. The first crossover is at $\Delta t_{chr,fit}^{+} \approx (640 \pm 40)\tau_0$ and MSD of $\approx (1.6 \pm 0.1)\xi^2(d)$, where $\xi^2(d)$ is measured from simulation. The second crossover is at $\Delta t_{chr,fit}^{++} \approx (4.6 \pm 0.9)\text{x}10^3\tau_0$ with MSD of $\approx (1.1 \pm 0.1) < r^2(2\lambda) >$, where $< r^2(2\lambda) >$ is the mean square size of a genomic length $2\lambda$ with active loop extrusion obtained from simulation.

#### III. Extended methods

##### a. Hybrid molecular dynamics – Monte Carlo (MD—MC) simulations

We use the Kremer-Grest bead-spring model (25) to simulate linear polymer chains with $N_{beads} = 1000$ beads, each with diameter $\sigma_{LJ}$ and each representing 1 kbp of chromatin. We use the Large-scale Atomic/Molecular Massively Parallel Simulator (LAMMPS) package (26). Non-bonded beads interact *via* a truncated and shifted Lennard-Jones (LJ) potential

$$U_{LJ}(r;\epsilon,r_c) = \begin{cases} 4\epsilon\left[\left(\frac{\sigma_{LJ}}{r}\right)^{12} - \left(\frac{\sigma_{LJ}}{r}\right)^{6} - \left(\frac{\sigma_{LJ}}{r_c}\right)^{12} + \left(\frac{\sigma_{LJ}}{r_c}\right)^{6}\right] & , \quad r \le r_c \\ 0 & , \quad r > r_c \end{cases} \tag{S34}$$

with $r_c = 2.5\sigma_{LJ}$ and $\epsilon = 0.298\epsilon_{LJ}$. $\epsilon_{LJ}$ is the LJ energy scale. $\tau_{LJ} = \left(m\sigma_{LJ}/\epsilon_{LJ}\right)^{1/2}$ is the LJ time scale, where $m$ is the mass of a bead. This choice of parameters simulates a theta-like implicit solvent (27). We use a finite extensible nonlinear elastic (FENE) potential to model bonded interactions along the backbone:

$$U_B(r) = -0.5KR_0^2 ln\left[1 - \left(\frac{r}{R_0}\right)^2\right] + U_{LJ}\left(r; 1\epsilon_{LJ}, 2^{\frac{1}{6}}\sigma_{LJ}\right), \tag{S35}$$

where $K = 30\epsilon_{LJ}\sigma_{LJ}^{-2}$, $R_0 = 1.5\sigma_{LJ}$, and $U_{LJ}(r;\epsilon,r_c)$ is given by Eq. (S34). We use a Langevin thermostat to maintain a temperature $T = 1\epsilon_{LJ}/k_BT$ with a damping constant $\Gamma = 1\tau_{LJ}^{-1}$. The integration time step was $\Delta t = 0.01\tau_{LJ}$. The Kuhn length for this chain is $\approx 2\sigma_{LJ}$ corresponding to approximately two beads (2 kbp or one locus). The center of mass of the two central beads in the chain moved a mean square displacement on the order of the mean square size of a dimer in $\approx 3\tau_{LJ}$, so the diffusion time of a Kuhn segment is approximately $\tau_0 \approx 3\tau_{LJ}$.

Chains were initialized in one of three ways:

1) Random walks in a cubic box with length 500 $\sigma_{LJ}$ and then equilibrated for $10^7\tau_{LJ}$,
2) Hairpin with two parallel straight lines of 500 beads each, or
3) Random walks in a cubic box, confined with a potential $U_{conf}(R_g) = 15\epsilon_{LJ}\sigma_{LJ}^{-2}\left(R_g - 5\sigma_{LJ}\right)^2$ and equilibrated for $2\text{x}10^6\ \tau_{LJ}$.

Conditions 1) and 3) were used for simulations with a single extrusion cycle (when only one cohesin extrudes the chain one time). Conditions 1) and 2) were used for steady-state simulations. The initial box sizes were chosen to be much larger than the initial polymer size. Simulations used non-periodic, shrink-wrapped boundaries such that the box adjusts during the simulations to ensure beads do not cross the boundaries. The confining potential for condition 3) caused an average radius of gyration $R_g \approx (6.6 \pm 0.1)\sigma_{LJ}$ before extrusion, which is approximately three times smaller than the unconfined radius of gyration. The confining potential was maintained throughout extrusion.

We then used hybrid MD—MC to simulate loop extrusion implemented in LAMMPS and C (MD in LAMMPS followed bead trajectories; MC in C decided cohesin binding, unbinding, and translocation). One MC step was performed every $\tau_{MC} = 2\tau_{LJ}$, $5\tau_{LJ}$, or $10\tau_{LJ}$. Single extrusion cycle simulations were run with all three $\tau_{MC}$, while steady-state extrusion simulations were run with $\tau_{MC} = 5\tau_{LJ}$. Cohesins were modeled by additional bonds with the potential

$$U_c(r) = -0.5K_c R_{0_c}^2 ln\left[1-\left(\frac{r}{R_{0_c}}\right)^2\right] + U_{LJ}\left(r; 1\epsilon_{LJ}, 2^{\frac{1}{6}}\sigma_{LJ}\right), \quad \text{(S36)}$$

where $K_c = 1\epsilon_{LJ}\sigma_{LJ}^{-2}$ and $R_{0_c} = 4\sigma_{LJ}$. To simulate extrusion, this bond can form, break, and change binding partners during MC steps. We randomize the order of cohesin binding, unbinding, and translocation during each MC step. We denote one side of the cohesin bond the (-) domain, and the other the (+) domain. Steady-state simulations were run for $10^7\tau_{LJ}$, which is more than $10^3$ times the end-to-end vector autocorrelation decay time (see Fig. S8a). We compared three different initial chromatin conformations (two different relaxed conformations, and a hairpin) to ensure that the initial conformation did not affect the contact probabilities in the simulation. The average contact probabilities across all three initial conformations were consistent (see Fig. S8b). Two beads were considered to be in contact if the distance between their centers was shorter than or equal to $1.5\sigma_{LJ}$.

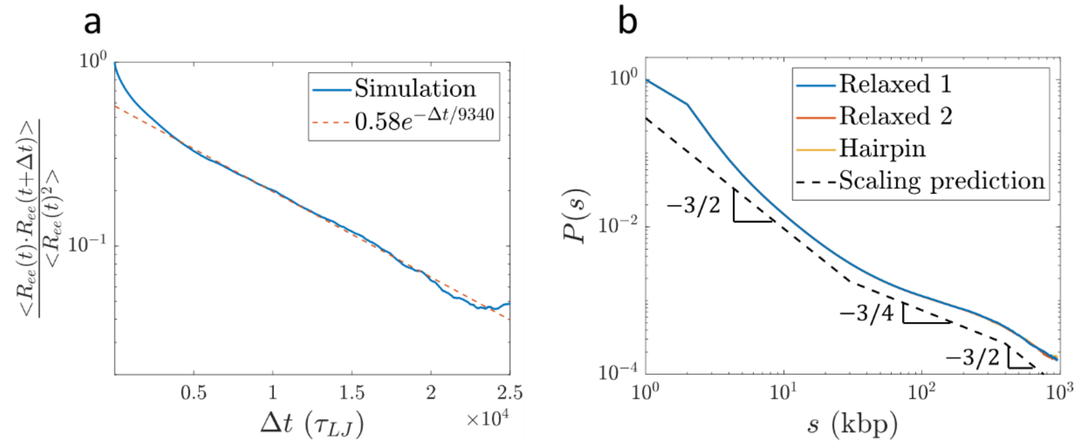


**Figure S8:** Equilibration of hybrid MD—MC simulations. a) Autocorrelation of the end-to-end vector of a steady-state extrusion simulation with $\tau_{MC} = 5\tau_{LJ}$. The dashed red line is a fit in the range $\Delta t = 5000\tau_{LJ} - 25000\tau_{LJ}$. The decay time is approximately $(9340 \pm 20)\tau_{LJ}$. b) Average contact probabilities of steady-state extrusion simulations for three initial conformations, all using $\tau_{MC} = 5\tau_{LJ}$.The dashed black lines indicate our scaling prediction shifted vertically. The blue, red, and yellow curves mostly overlap one another.

<u>Translocation</u>

Let the (-) and (+) cohesin domains be located at beads $a$ and $b$ respectively such that $1 \leq a < b \leq N_{beads}$. The two domains move independently and cannot move past TAD anchors in the correct orientations. During each MC step, the (-) domain attempts a move to bead *a-1* with probability $p = 0.5$. The (+) domain attempts a move to bead $b$+*1* with probability $p = 0.5$. Moves are accepted if the domains are not at TAD anchors and the attempted bond length is shorter than $R_{0_c} - 0.005 = 3.995\sigma_{LJ}$ (to prevent over-stretched FENE bonds). If the (-) domain is at bead index 1 and attempts to move, the cohesin unbinds. Similarly, if the (+) domain is at bead index $N_{beads} = 1000$ and attempts to move, the cohesin unbinds. For almost all simulations, multiple cohesin domains are allowed to bind the same bead (*i.e.*, unimpeded cohesin traversal is allowed). Using these parameters, the average extrusion velocity per domain with $\tau_{MC} = 5\tau_{LJ}$, $2\tau_{LJ}$, and $10\tau_{LJ}$ were $v_{ex} \approx 0.2$ kbp/$\tau_{LJ} \approx 0.6$ kbp/$\tau_0$, $v_{ex} \approx 0.1$ kbp/$\tau_{LJ} \approx$ 0.3 kbp/$\tau_0$, and $v_{ex} \approx 0.05$ kbp/$\tau_{LJ} \approx 0.15$ kbp/$\tau_0$ respectively, where $\tau_0$ is the diffusion time of a Kuhn segment.

Cohesin binding and unbinding

For simulations of a single extrusion cycle, we place a single cohesin bond halfway along the polymer chain with the (-) and (+) domains at bead indices 500 and 501, respectively. Recall that we represent cohesin by a single bond between two beads (see Eq. (S36) and related discussion). The cohesin translocates until reaching the ends of the polymer chain. Only one cohesin is allowed on the chain. For steady-state simulations, we place no limit on the number of cohesins on the chain. We randomly choose a pair of beads with indices *x* and *x+1* such that $1 \leq x < N_{beads}$. A cohesin bond is formed with probability $p_{bind}$, which we set to $p_{bind}$ =2.8x10$^{-5}$. During a single MC step, we cycle through all possible bead pairs *x* and *x+1* in random order. Also, during each MC step, all bound cohesins attempt to unbind with probability $p_{unbind}$, which we set to $p_{unbind} = 0.004$. These parameters with $\tau_{MC} = 5\tau_{LJ}$ without TAD anchors yielded a mean residence time of $\tau_{res} \approx (960 \pm 10)\tau_{LJ}$, processivity of $\lambda \approx (190 \pm 2)$ kbp, and separation of $d \approx (190 \pm 20)$ kbp.

Simulations with immobile cohesins

For Figure S2a and Figure 10b in the main text, we ran simulations with cohesins that do not move in space. To do this, we include a harmonic potential to the beads held together by the cohesin bond. This potential tethers the beads' center of mass (COM) to the initial COM of the cohesin binding site. This potential is given by $U_{tether}(r_{COM}) = K_t {r_{COM}}^2$, where $K_t = 1\epsilon_{LJ}\sigma_{LJ}^{-2}$ and $r_{COM}$ is the distance between the COM of the cohesin beads and the tether point. During each integration time step, the tethering potential is only applied to beads held together by cohesin.

Simulations with two cohesins

For Figure S3d, we simulate two unnested cohesins actively extruding the same polymer. These simulations are performed like the single extrusion cycle simulations, except we initially place one cohesin at bead indices 332 and 333, and the other at indices 666 and 667. In these simulations, we only allow one cohesin domain per bead at a time, meaning cohesin traversal is not allowed.

Determining $g_{min}$ and $g(d)$ from simulation

To measure $g_{min}$ and $g(d)$, we compared the mean square displacements of beads $MSD_{passive}(\Delta t)$ *without* extrusion to internal sizes $< r_{passive}^2(s) >$ *without* extrusion. $< {r_{passive}}^2(s) >$ is the mean square size of a section with *s* beads averaged over different sections within the polymer chain. The MSD at lag time $\Delta t$ is the mean square distance a bead moves in the time it takes a cohesin domain to extrude $v_{ex}\Delta t$ beads. $< r_{passive}^2(v_{ex}\Delta t) >$ is the relaxed size of a section with $v_{ex}\Delta t$ beads. $g_{min}$ is the largest section such that $MSD_{passive}(v_{ex}\Delta t) \geq < r_{passive}^2(v_{ex}\Delta t) >$. The mean square size of $g_{min}$ is $\xi_{min}^2 \approx < r_{passive}^2(g_{min}) >$. The lag time at which this occurs is the relaxation time of a section with genomic length $g_{min}$ which is also $\Delta t_{coh}^+$ (see Eq. (S24) and related discussion). Figure S9 shows an example for $v_{ex} \approx 0.3$ kbp per $\tau_0$ where $g_{min} \approx (15 \pm 1)$ kbp, $\xi_{min}^2 \approx (27 \pm 2)\sigma_{LJ}^2$, and $\Delta t_{coh}^+ \approx (50 \pm 3)\tau_0$ obtained from simulation (the intersection of the red and blue curves in Figure S9). Errors represent the range of the intersection found by using the simultaneous, observational bounds at 95% confidence intervals of power law fits to the red and blue curves. For $v_{ex} \approx 0.6$ kbp per $\tau_0$ (used in Fig. S6b), $g_{min} \approx (7 \pm 1)$ kbp, $\xi_{min}^2 \approx (12 \pm 2)\sigma_{LJ}$, and $\Delta t_{coh}^+ \approx (12 \pm 1)\tau_0$.

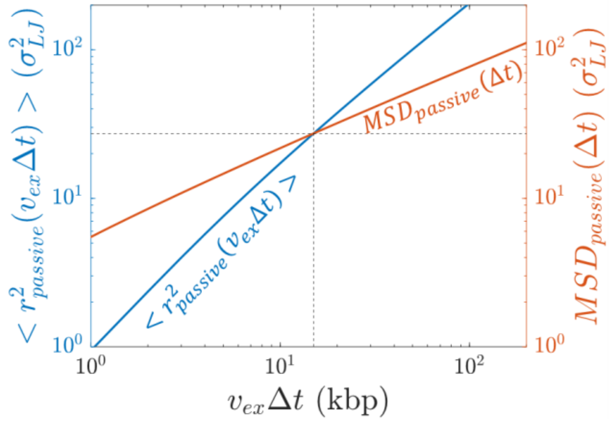


**Figure S9:** Bead MSD and mean square internal sizes of a simulation without extrusion, used to obtain $g_{min}$. This example uses $v_{ex} = 0.3$ kbp/$\tau_0$. The dashed black lines indicate the intersection of the red and blue curves and represent $g_{min}$ and the relaxed, mean square size of $g_{min}$ from simulation.

$g(d)$ is the largest chromatin section that relaxes during time $\tau_{B,\lambda} \approx d/(2v_{ex})$ (see section IIb). $\tau_{B,\lambda}$ is the time between cohesin binding events within a chromatin section of genomic length $\lambda$. In our steady-state simulations, $\tau_{B,\lambda} \approx (940 \pm 10)\tau_{LJ}$. We obtain $g(d)$ from simulation as the largest section such that $MSD_{passive}(940\tau_{LJ}) \geq < r^2_{passive}(s) >$. $\xi^2(d)$ is $< r^2_{passive}(g(d)) >$, the relaxed, unextruded mean square size of a section with genomic length $g(d)$. We find $g(d) \approx (37 \pm 1)$ kbp and $\xi^2(d) \approx (73 \pm 2)\sigma^2_{LJ}$.

<u>Approximating $g_{leg}$ and tension fronts from simulation</u>

To approximate the number of beads in a leg produced by extrusion, we simulate a single extrusion cycle using three extrusion velocities. Each simulation started from a relaxed polymer conformation with $D \approx 2$. We track $< d_0^2(t;i) >$, the mean square distance between the initial position of bead *i* and at time $t$ after extrusion begins (see Fig. S10a). Before the tension front reaches bead $i$, $< d_0^2(t;i) >$ follows the bead MSD from a passive chain without extrusion $MSD_{passive} \sim t^{1/2}$. After the tension front reaches bead *i* at time $t_{feel}$, $< d_0^2(t;i) >$ deviates from $MSD_{passive}$ and grows faster than $t^{1/2}$ (see Fig. S10b). To find $t_{feel}$, we fit the second regime of $< d_0^2(t;i) >$ averaged over 100 simulations per extrusion velocity to a power law and find the time at which $< d_0^2(t;i) >$ intersects $MSD_{passive}$ from a simulation without extrusion. Each bead *i* thus has an associated $t_{feel}$. From each simulation, we calculate $g_{leg}(t_{feel})$, the number of beads in a leg at time $t_{feel}$, by finding the number of beads between *i* and the bead being extruded at $t_{feel}$ (see Fig. S10a). On average we expect $g_{leg}(t_{feel}) \approx s_{bind,i} - v_{ex}t_{feel}$, where $s_{bind,i}$ is the number of beads between the cohesin binding site and the bead *i* at the end of the tension front at time $t_{feel}$.

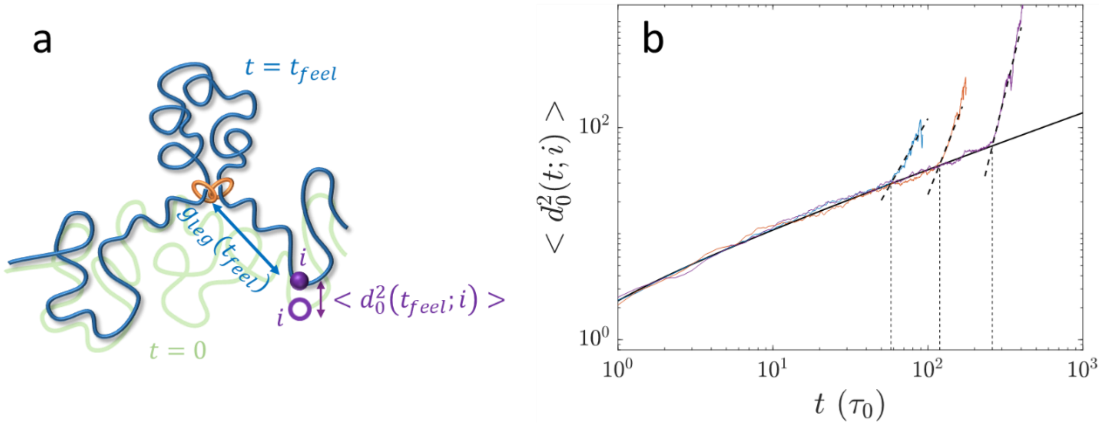


**Figure S10:** Approximating the length of extended chromatin legs from simulations. a) Schematic showing bead $i$ moving in space from its initial position (open purple circle) to its position at time $t_{feel}$ (filled purple circle) corresponding to a mean square distance of $< d_0^2(t_{feel}; i) >$. The green and blue curves represent the chromatin conformation at $t = 0$ when cohesin starts and at $t = t_{feel}$ respectively. $g_{leg}(t_{feel})$ is the number of beads between cohesin (orange rings) and $i$ at time $t_{feel}$. b) Mean square distance $< d_0^2(t; i) >$ between a bead's position at time $t$ and its initial position from simulations of a single extrusion cycle on an initially relaxed chain ($D \approx 2$) using $v_{ex} \approx 0.6$ kbp/$\tau_0$. The blue, red, and purple curves correspond to beads that are 50 kbp, 100 kbp, and 200 kbp away from the cohesin binding site, respectively. The thick black dashed lines are fits to the second regimes of the $< d_0^2(t; i) >$ curves. The vertical dashed lines are the intersections between the estimated $t_{feel}$ for the different beads.

<u>Cohesin trajectories and tension fronts from simulations</u>
Each cohesin produces two tension fronts. As described in the main text and section IId above, cohesin trajectories are determined by the midpoint between the two tension fronts. The tension front midpoint is determined by the initial conformation of the chromatin section of interest. To test this, we simulate a single extrusion cycle on an initially relaxed polymer chain. We run 200 simulations, all with the same initial chain conformation and cohesin binding site, but with different random number generator seeds. The cohesin trajectory is averaged over the 200 simulations.

From the initial chain conformation, we estimate the positions of the tension fronts at time $t$ as the positions of the beads $g_{leg}(t) \approx [2.1(v_{ex}t)^{0.54}]$ (see Eq. (S22)) past the cohesin binding site on either side, where $[\ldots]$ denotes the integer part (see Fig. S11a). As described in the previous section and the discussion related to Eq. (S22), we found the expression for $g_{leg}(t)$ by fitting the calculated genomic lengths of legs at time $t$ to a power law $g_{leg}(t) \approx A * (v_{ex}t)^B$ resulting in $A = 2.1 \pm 0.5$ and $B = 0.54 \pm 0.05$. The fit was performed simultaneously on data from 100 simulations per extrusion velocity. Our objective is to approximate the average path of the tension front midpoints produced by any cohesin that extrudes a given chromatin section with a given initial conformation and a given binding site (see Fig. 11B in the main text). One initial chain conformation has one set of tension front midpoints.

To smooth the tension front midpoints, we average the positions of all beads $i$ that satisfy

$$\max[0, g_{leg}(t) - 2g_{smooth}(t)] \le s_{ij} \le \min[g_{leg}(t) + 2g_{smooth}(t), N_{beads} - j], \qquad \text{(S37)}$$

where $s_{ij}$ is the genomic distance between $i$ and the cohesin binding site at $j$ and $N_{beads} - j$ is the maximum genomic distance between the polymer chain end and the cohesin binding site (see

Fig. S11a). $g_{smooth}(t)$ is the number of beads in a section of a relaxed, unperturbed chain with mean square size approximately equal to the mean square displacement of a single bead in the relaxed, unperturbed chain during time $\Delta t$. For each time $t$, we measure $g_{smooth}(t)$ from a simulation of a relaxed polymer chain *without* extrusion and fit a power law $g_{smooth}(t) \approx (2.10 \pm 0.01) * (t/\tau_0)^{(0.504 \pm 0.001)}$ as shown in Figure S11b. Errors represent 95% confidence intervals. Smoothing over $2g_{smooth}(t)$ effectively averages the locations of two tension fronts over thermal fluctuations of several polymer sections that relax during time *t*. The max[…] and min[…] functions in Eq. (S37) ensure that the smoothing does not extend past the polymer chain ends nor the cohesin binding site.

Figure S11c and S11d show the unsmoothed and smoothed tension front midpoints from the same initial chain conformation, respectively. The smoothed midpoints are used for Figure 11B in the main text. Figure S11e plots the mean square distance between cohesin trajectories and the tension front midpoint trajectory $< \Delta^2_{coh,tens}(t) >$ (see Fig. S6b) for both the smoothed (blue curve) and unsmoothed (red curve) tension front midpoint. Smoothing the tension front midpoint trajectory suppresses fluctuations in $< \Delta^2_{coh,tens}(t) >$ but maintains the same scaling of $\sim t^{1/2}$.

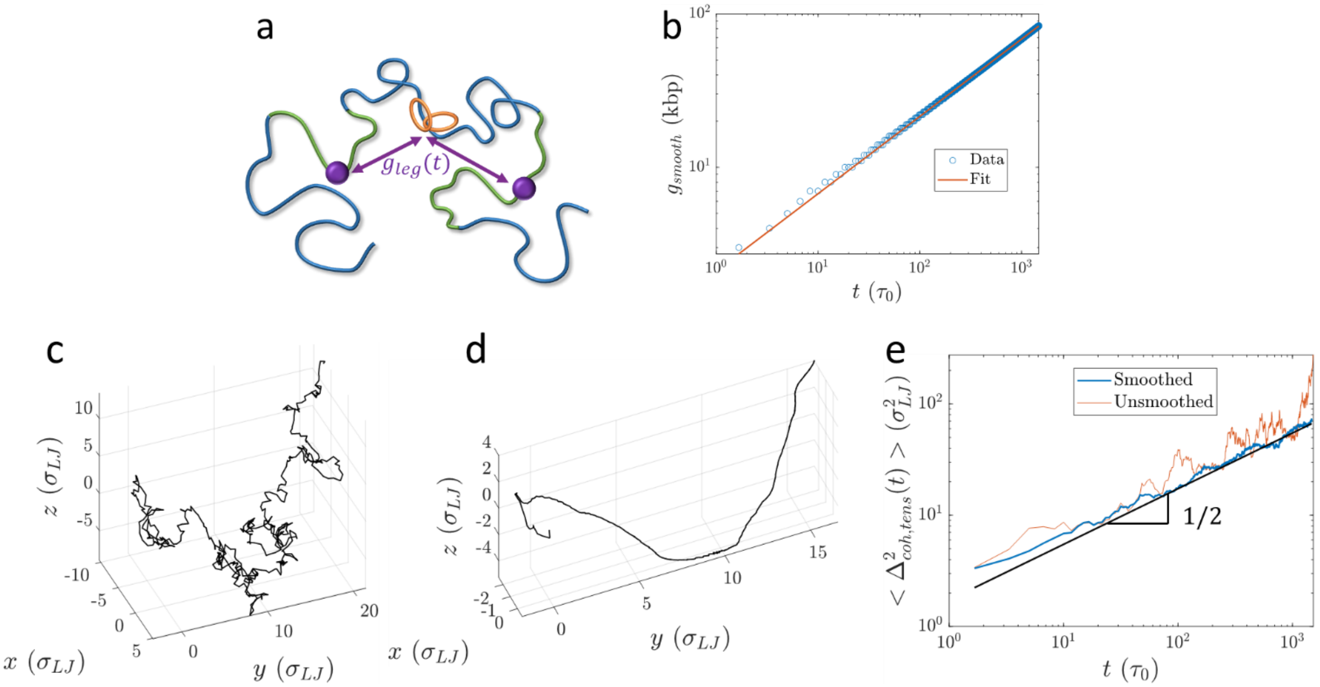


**Figure S11:** Trajectories of tension front midpoints from initial chain conformations. a) The tension front positions are approximated from the initial chain conformation before extrusion begins as the positions of the beads $g_{leg}(t)$ past the cohesin binding site (purple circles) when extrusion begins. To smooth the tension fronts, we take the average position of all beads that satisfy Eq. (S37) (green curves) when extrusion begins. b) The number of beads $g_{smooth}(t)$ in a section of a relaxed polymer chain that has approximately the same mean square size as the mean square displacement of a single bead during time $\Delta t$. The red line is a power law fit to the data yielding $g_{smooth}(t) \approx (2.10 \pm 0.01) * (t/\tau_0)^{(0.504 \pm 0.001)}$. c) Unsmoothed tension front midpoint trajectories taken from a single initial chain conformation. d) Smoothed tension front midpoint trajectories taken from the same initial chain conformation as in c). e) Mean square distance between cohesin trajectories and tension front midpoints at time *t* after extrusion starts from simulations of a single cohesin extruding an initially relaxed chromatin section with $D \approx 2$ and $v_{ex} \approx 0.3$ kbp/$\tau_0$. The blue and red curves use the smoothed and unsmoothed tension front midpoints, respectively. The black curve is $(3t)^{1/2}$.

**b. Contact probability plots from Micro-C data**

We used publicly available Micro-C data from HFF and mESC cells. HFF data was from Ref. (28), accessed in the 4D Nucleome Data Portal with accession number 4DNFI9FVHJZQ. mESC data was from Ref. (29), accessed in the Gene Expression Omnibus with accession number GSE178982, file GSE178982_CTCF-UT_pool.mcool. cooltools (30) was used to calculate smoothed contact probabilities $P(s)$ averaged over all chromosomes.

**c. Inter-TAD and intra-TAD contacts**

TADs in HFF cells were identified with the Arrowhead algorithm in Juicer tools version 1.22.01 (31) with 5 kbp resolution. TADs in mESC cells were taken from Supplemental Table 3 in (29), which were also identified using Arrowhead. Contacts were accessed from .mcool files at 1 kbp resolution using the matrix selector in cooler (32) with the balance=True option. Consider a TAD as depicted by the dark red square in the center of the contact map schematic in Figure S12.

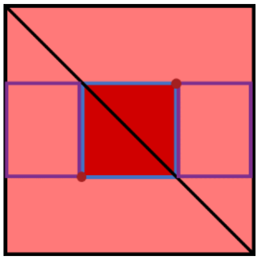

**Figure S12:** Schematic contact map with a TAD in the center, represented with the dark red square. Purple and blue rectangles represent inter-TAD and intra-TAD contacts, respectively.

The inter-TAD contacts $C_{inter}$ are calculated as the sum of all contacts outlined by the purple rectangles, extending to the chromosome ends. These are all contacts between loci in the TAD with loci outside of the TAD but within the same chromosome. The intra-TAD contacts $C_{intra}$ are calculated as the sum of all contacts outlined by the blue square, not including the main diagonal entries. These are the contacts between loci in the TAD with other loci in the TAD. $C_{tot} = C_{inter} + C_{intra}$ is the total number of contacts made by loci within the TAD. We bin the TADs in the analyzed datasets by genomic length $N_{TAD}$ into 100 log-spaced bins. The bin-averaged fraction of inter-TAD and intra-TAD contacts ($F_{inter}(N_{TAD})$ and $F_{intra}(N_{TAD})$, respectively) are plotted in Figure 9 of the main text.

Given an average contact probability function $P(s)$, the number of inter-TAD contacts should be approximately proportional to

$$I_{inter}(N_{TAD}) \approx \int_{1}^{N_{TAD}} sP(s)ds + \int_{N_{TAD}+1}^{\overline{L_{chr}}-N_{TAD}} N_{TAD}\, P(s)ds\,, \tag{S38}$$

where $\overline{L_{chr}}$ is the average genomic length of a chromosome which we take to be approximately 100 Mbp. We calculate $I_{inter}$ using the smoothed crossover function for contact probabilities shown in Figure 8c of the main text:

$$P_1(s) = s^{-\frac{3}{2}}\left[1+\left(\frac{s}{30}\right)^2\right]^{\frac{3}{8}}\left[1+\left(\frac{s}{400}\right)^2\right]^{-\frac{3}{8}}. \tag{S39}$$

We recognize that Eq. (S39) does not capture the scaling behavior past the entanglement genomic length, after which the contact probabilities may be proportional to $s^{-1}$, but assume that large genomic distances $s$ contribute minimally to the integrals in Eq. (S38). An example of a crossover function that has contact probabilities $\sim s^{-1}$ for $s \geq N_{e,active} \approx 5000$ kbp is

$$P_2(s) = s^{-\frac{3}{2}}\left[1 + \left(\frac{s}{30}\right)^2\right]^{\frac{3}{8}}\left[1 + \left(\frac{s}{400}\right)^2\right]^{-\frac{3}{8}}\left[1 + \left(\frac{s}{5000}\right)^2\right]^{\frac{1}{8}}. \quad \text{(S40)}$$

The values of $I_{inter}(N_{TAD})$ using Eq. (S39) and Eq. (S40) differ by ≤ 10% for TADs shorter than 1 Mbp, and ≤ 15% for TADs between 1 Mbp and 4 Mbp. Numerical integration was performed using the *trapz()* function in MATLAB. Figure S13a shows the calculated $I_{inter}(N_{TAD})$ scaled to match $C_{inter}(N_{TAD})$ at $N_{TAD} = 200$ kbp $\approx \lambda$ from Micro-C data in HFF and mESC cells. We call this scaled integral $\widetilde{I_{inter}}(N_{TAD})$.

Ideally, $\overline{C_{loc}} \approx C_{tot}(N_{TAD})/N_{TAD}$ or the average number of contacts made per locus, should be independent of TAD length. However, as shown in Figure S13b, the Micro-C data shows a weak dependence of $\overline{C_{loc}}$ on $N_{TAD}$. This discrepancy could be attributed to inefficient cross-linking during the Micro-C experiments, minor sequencing errors, artifacts of the downstream processing to produce contact matrices, or unexplored biological phenomena. To correct for this effect, we fit separate power laws to the data in HFF and mESC cells (see Fig. S13b). We call the fitted power laws $\overline{C_{loc,fit}}(N_{TAD})$. Our predicted inter-TAD and intra-TAD fractions are then calculated as

$$F_{inter,predict}(N_{TAD}) \approx \frac{\widetilde{I_{inter}}(N_{TAD})}{N_{TAD}\overline{C_{loc,fit}}(N_{TAD})} \quad \text{(S41)}$$

and

$$F_{intra,predict}(N_{TAD}) \approx 1 - F_{inter,predict}(N_{TAD})\,, \quad \text{(S42)}$$

respectively. Figures S13c and S13d show $F_{inter}(N_{TAD})$ and $F_{intra}(N_{TAD})$, respectively, from Micro-C data (circles) as well as from Eqs. (S41) and (S42) (blue and green curves). The purple curves show the inter- and intra-TAD fractions predicted *without* scaling the integral $I_{inter}(N_{TAD})$ and *without* correcting for the $N_{TAD}$ dependence of $\overline{C_{loc}}$. The scaling and correcting steps are necessary to match the magnitudes and shapes of the data more accurately.

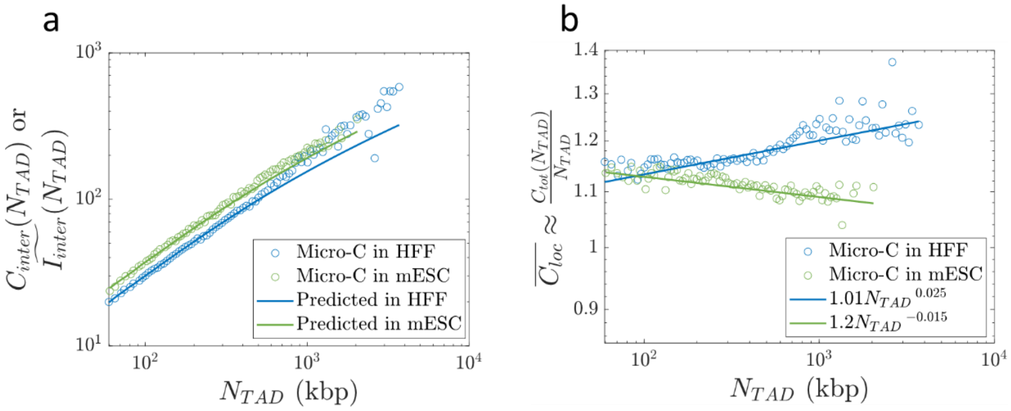

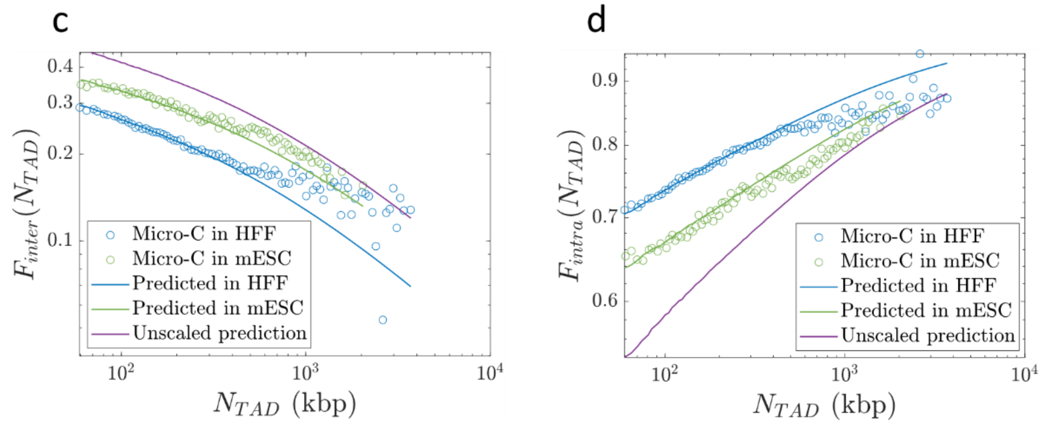


**Figure S13:** Inter-TAD and intra-TAD contacts. a) Sum of inter-TAD contacts for a given TAD length $C_{inter}(N_{TAD})$ from Micro-C experiments compared to the predicted number $I_{inter}(N_{TAD})$ (Eq. (S38)) scaled to be equal to $C_{inter}$ at $N_{TAD} = 200$ kbp. b) Total number of intra-chromosomal contacts made by a TAD $C_{tot}(N_{TAD})$ normalized by TAD length $N_{TAD}$. The data for HFF and mESC cells were fit to separate power laws (solid lines). c) Inter-TAD fraction of intra-chromosomal contacts made by a TAD with length $N_{TAD}$ from experimental data (circles) compared to predicted values (blue and green curves, Eq. (S41)) and unscaled predictions (purple curve). d) Same as c) but for the intra-TAD fraction of intra-chromosomal contacts, where the blue and green curves use Eq. (S42).

### IV. Supplementary tables

Table S1: Description of notations used in the main text.

| Parameter | Description |
|---|---|
| $b$ | Locus size |
| $z$ | Genomic length per locus |
| $v_{ex}$ | Extrusion speed (in units of genomic length/time) |
| $\tau_0$ | Diffusion time of a locus |
| $\kappa = v_{ex}\tau_0 z^{-1}$ | Extrusion ratio |
| $l = 2v_{ex}t$ | Genomic length of an extruded loop |
| $s$ | Genomic length |
| $s_{c,i}$ | Genomic distance between cohesin and locus *i* within a loop |
| $s_{c,out}$ | Genomic distance between cohesin and locus of interest outside a loop |
| $g_{min}$ | Shortest genomic length of a relaxed chromatin section within an extruded loop |
| $g(l)$ | Longest genomic length of a relaxed chromatin section within a loop of length *l*, predicted to be Eq. (S7) |
| $g(d)$ | Longest genomic length of a relaxed chromatin section in steady state extrusion, predicted to be Eq. (10) in the main text |
| $\xi(l)$ | Size of the longest relaxed chromatin section within a loop of length *l* |
| $< R^2(l) >$ | Mean square size of a loop with length *l* |
| $< {r_l}^2(s) >$ | Mean square size of a section with genomic length *s* within a loop of length *l* |
| $\lambda$ | Processivity (genomic length) |
| $d$ | Separation (genomic length) |
| $\overline{N_{TAD}}$ | Average genomic length of a TAD |
| $\tau_{res}$ | Residence time of cohesin |
| $P(s)$ | Contact probability |
| $\phi_{self}(s)$ | Volume fraction of a section with length *s* within its pervaded volume where the section is part of a genomic section of length *l* |
| $\phi_z$ | Volume fraction of chromatin (histones and DNA) within the pervaded volume of a locus with *z* base pairs |
| $\phi$ | Average volume fraction of chromatin in nucleus |
| $N_{e,passive}$ | Entanglement genomic length in the absence of activity |
| $N_{e,active}$ | Entanglement genomic length with active loop extrusion |
| $O(s)$ | Overlap parameter |
| $D$ | (Apparent) fractal dimension |
| $g_{leg}$ | Genomic length of leg outside of loop |
| $< r_{leg}^2 >$ | Mean square size of leg |

Table S2: Range of parameter estimates given an extrusion ratio range of $\kappa \approx 0.003 - 0.3$. All estimates use a locus discretization of $z = 2$ kbp, locus size $b = 50$ nm, extrusion velocity $v_{ex} = 0.1$ kbp per second, and cohesin processivity and separation $\lambda = d = 200$ kbp.

| Parameter | $\kappa \approx 0.003$ | $\kappa \approx 0.2$ (used in main text) | $\kappa \approx 0.3$ |
|---|---|---|---|
| $g_{min}$ | 700 kbp | 10 kbp | 7 kbp |
| $g(\lambda)$ in a loop with processivity $\lambda$ | 200 kbp[1] | 30 kbp | 26 kbp |
| $g(d)$ | N/A[2] | 30 kbp | 26 kbp |

[1] Using $\kappa \approx 0.003$, entire loops are relaxed up to loop lengths $g_{min} \approx 700$ kbp. Loops with $\lambda = 200$ kbp are fully relaxed.

[2] Since entire loops are relaxed with these parameters, $g(d)$ does not exist as the system would be in the "relaxed" Regime I in Figure 7a of the main text.

Table S3: Range of the passive and active entanglement genomic lengths for $O_{KN} = 10 - 20$ and average chromatin volume fraction $\phi = 0.06 - 0.4$. All estimates use a locus discretization of $z = 2$ kbp, locus size $b = 50$ nm, physical volume of a locus $v \approx 7.5\text{x}10^3$ nm$^3$, average TAD length $\overline{N_{TAD}} \approx 200$ kbp and $g(d) = 30$ kbp.

| | **Lower bound** | **Main text estimate** | **Upper bound** |
|---|---|---|---|
| $N_{e,passive}$ | 16 kbp | 100 kbp | 3 Mbp |
| $N_{e,active}$ | 800 kbp | 5 Mbp | 150 Mbp |
